# Limits and constraints on mechanisms of cell-cycle regulation imposed by cell sizehomeostasis measurements

**DOI:** 10.1101/720292

**Authors:** Lisa Willis, Henrik Jönsson, Kerwyn Casey Huang

## Abstract

High-throughput imaging has led to an explosion of observations regarding cell-size homeostasis across the kingdoms of life. Among bacteria, “adder” behavior in which a constant size appears to be added during each cell cycle is ubiquitous, while various eukaryotes show other size-homeostasis behaviors. Since interactions between cell-cycle progression and growth ultimately determine size-homeostasis behaviors, we developed a general model of cell proliferation to: 1) discover how the requirement of cell-size homeostasis limits mechanisms of cell-cycle control; 2) predict how features of cell-cycle control translate into size-homeostasis measurements. Our analyses revealed plausible cell-cycle control scenarios that nevertheless fail to regulate cell size, conditions that generate apparent adder behavior without underlying adder mechanisms, cell-cycle features that play unintuitive roles in causing deviations from adder, and distinguishing predictions for extended size-homeostasis statistics according to the underlying control mechanism. The model thus provides holistic insight into the mechanistic implications of cell-size homeostasis measurements.

## Introduction

One of the most fundamental questions in biology is how cells regulate cell-cycle progression, which is intimately tied to myriad processes such as cell-size determination (Schmoller et al., 2015), drug sensitivity (Shi et al., 2017), and transcription (Padovan-Merhard et al., 2015). In all organisms, cell-cycle control must be coupled to growth to ensure cell-size homeostasis, the maintenance of a fixed average size in steady conditions. Measurable size-homeostasis behaviors are determined by interactions between cell-cycle control and growth. Single-cell lineage tracking and cellcycle reporters have led to a rapid proliferation in size homeostasis measurements across bacteria, yeast, mammalian cells, and plant cells. Among bacteria (Campos et al., 2014; Taheri-Araghi et al., 2015; Wallden et al., 2016; Willis and Huang, 2017) and an archaeon (Eun et al., 2018), a common theme has emerged: cells appear to regulate their size via an “adder” behavior whereby a fixed volume is added between birth and division. Among eukaryotes, budding yeast and mammalian cells can deviate from adder behavior over the G1 and S/G2 cell-cycle stages while maintaining apparent adder or near-adder behavior between birth and division (Cadart et al., 2018; Chandler-Brown et al., 2017; Di Talia et al., 2007; Schmoller et al., 2015), with the smallest mammalian cells switching to approximately “sizer” behavior with no correlation between birth and division sizes (Varsano et al., 2017). Similarly, small fission yeast exhibit sizer behavior at division while large fission yeast exhibit near-adder behaviour (Facchetti et al., 2019; Fantes, 1977; Pan et al., 2014). By contrast, stem cells of *Arabidopsis thaliana* exhibit intermediate adder-sizer behavior (Willis et al., 2016).

Despite the recent explosion of size-homeostasis measurements, there is no clarity as to the implications of these similarities and differences for mechanisms of cell-cycle control and its coupling to growth. Furthermore, despite the centrality of these concepts, how the necessity for size homeostasis limits mechanisms of cell-cycle control is not understood. We sought to develop a theoretical framework to address two major questions: how does the requirement for cell-size homeostasis limit cell-cycle regulator dynamics and mechanisms of cell-cycle checkpoint progression, and to what extent are size-homeostasis measurements informative about underpinning mechanisms of cellcycle regulation? Importantly, it is unresolved whether the widely observed “adder” behavior implies a common mechanism across diverse organisms.

Seminal studies have revealed how cell-cycle progression is coupled to growth in several model organisms. In budding yeast, the G1/S inhibitor Whi5 is produced throughout S/G2/M and then diluted out by growth during G1 to trigger G1/S upon reaching a threshold minimum concentration (Schmoller et al., 2015). Mathematical models showed that for budding yeast-like proliferation dynamics, this “inhibitor-dilutor” G1/S regulation imparts adder behavior between birth and division (ChandlerBrown et al., 2017; Heldt et al., 2018; Soifer et al., 2016). Whi5 has functional homologs in mammals (Rb) and plants (RBR1), suggesting that an inhibitor-dilutor mechanism may regulate G1/S. In the bacterium *Escherichia coli*, the division protein FtsZ is a “master regulator” of division, with newly synthesized FtsZ accumulating at midcell proportionally with cell growth to trigger division at a total intracellular threshold level (Sekar et al., 2018; Si et al., 2019), a mechanism that recapitulates the observed adder behavior. Similarly, active DnaA, which accumulates at the origins of replication, effects adder behavior both between consecutive G1/Ss and between consecutive divisions if it is produced proportionally with growth and triggers replication initiation (G1/S) at a threshold level per origin when it is inactivated while a fixed time or added-size increment elapses between G1/S and division (Amir, 2014; Barber et al., 2017; Ho and Amir, 2015; Logsdon et al., 2017). DnaA followed by a fixed time interval and FtsZ mediated division may operate simultaneously, with the slower process triggering cell division (Micali et al., 2018a; Micali et al., 2018b; Si et al., 2019). DnaA and FtsZ are broadly conserved among bacteria but details of their dynamics and other proliferation factors are likely to vary; the extent to which size homeostasis limits their dynamics, and how they can account for the apparent universality of adder behavior across bacteria are unclear. Master regulators also control cell cycle-checkpoint progression in eukaryotes: the broadly conserved CDK1-cyclin (Harashima et al., 2013) accumulates during growth to trigger G1/S then G2/M at successive threshold activity levels in engineered fission yeast (Coudreuse and Nurse, 2010). The CDK1-cyclin regulatory network is complex, but data indicate that it may result in a simple scaling relating active CDK1-cyclin accumulation to cell size (Keifenheim et al., 2017; Patterson et al., 2019).

Here, we develop a general model of cell proliferation and thus predict the size-homeostasis behaviors produced by a wide range of cell-cycle control mechanisms. Instances of the model focus on cells with two phases partitioned by the major eukaryotic cell-cycle checkpoints: G1/S and G2/M (assuming that G2/M and division are coincident), and on two rate-limiting mechanisms of irreversible checkpoint progression: master regulators like CDK1-cyclin or FtsZ/DnaA that accumulate to threshold activity levels, and Whi5-like inhibitor dilutors. Previous models have focused on particular organisms with specific cell-cycle and growth regimes, and thus do not provide a comprehensive framework connecting proliferation dynamics to size-homeostasis measurements, or do not consider the mechanism coupling growth and cell-cycle progression and therefore lack predictive power for how genetic perturbations will affect size-homeostasis behavior. We systematically identify apparently plausible cell-cycle control scenarios that nevertheless fail to regulate cell size and are thus impossible. We describe how growth, noise origins, cell cycle checkpoint criteria, and cell-cycle regulator dynamics differentially impact size homeostasis measurements, and how additional size homeostasis measurements may be useful to discriminate among different underlying mechanisms that cause robust deviation from adder, as observed in *A. thaliana*. Taken together, this framework and the insights it provides should be broadly useful for interpreting and motivating cellsize homeostasis measurements across all organisms.

## Results

### A general model of cell proliferation with two cell-cycle checkpoints

Our models consider two types of checkpoint regulators motivated by present understanding of the eukaryotic cell cycle (Fig. 1A): 1) a master regulator (e.g., CDK1-cyclin) that accumulates from zero and triggers G1/S or G2/M progression upon reaching a total intracellular threshold level (absolute number of molecules), when it is immediately degraded; 2) an inhibitor dilutor (e.g. Whi5) that accumulates during one phase and is diluted out in the subsequent phase, triggering progression upon reaching a threshold minimum concentration with no subsequent degradation. Master regulators can accumulate through one phase, as is common for cyclins in eukaryotes, or two phases, as for CDK1-cyclin in an engineered model of fission yeast (Coudreuse and Nurse, 2010; Hochegger et al., 2008) and FtsZ/DnaA in slow-growing bacteria without multiple replication forks. Regulator production rates (*dC/dt*) can be cell-size dependent and may differ between phases according to

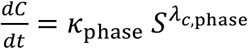

where *C* is the number of proteins, *S* is cell size, and *λ*_*c*,phase_, *k*_phase_ are parameters (Fig. 1A). Production rates change with cell size continually through the phase if *λ*_*c*,phase_ ≠ 0. The majority of proteins are thought to be maintained at constant concentrations during steady-state growth and thus likely are produced at a fixed rate proportional to cell size in exponentially growing cells (*λ*_*c*,phase_ = 1) (Newman et al., 2006; Padovan-Merhard et al., 2015; Schmoller and Skotheim, 2015), while Whi5 is produced independently of size through S/G2/M in budding yeast (*λ*_*c*,S/G2/M_ = 0) (Schmoller et al., 2015), and in fission yeast the activity of CDK1-cyclin may increase with a stronger size-dependence (*λ*_*c*,phase_ > 1) that results from multiple regulators with cell size-dependent levels (Keifenheim et al., 2017). The ratio of regulator production rates (*r*_S/G2/M_ = *k*_S/G2/M_/*k*_G1_) represents two extreme scenarios: either production is gene-copy number limited, meaning that the production rate doubles in S/G2/M upon gene duplication regardless of ploidy (*r*_S/G2/M_ = 2), or production is unaffected by gene-copy number (*r*_S/G2/M_ = 1) because another factor such as ribosome abundance is limiting (Heldt et al., 2018; Schmoller and Skotheim, 2015; Schmoller et al., 2015) (Fig. 1A). Proteins are assumed to be stable, consistent with measurements of key regulators, aside from targeted degradation (Hochegger et al., 2008; Schmoller et al., 2015). If a regulator persists through cell divisions, we assume it is inherited in proportion to daughter cell sizes without noise.

**Figure 1:**
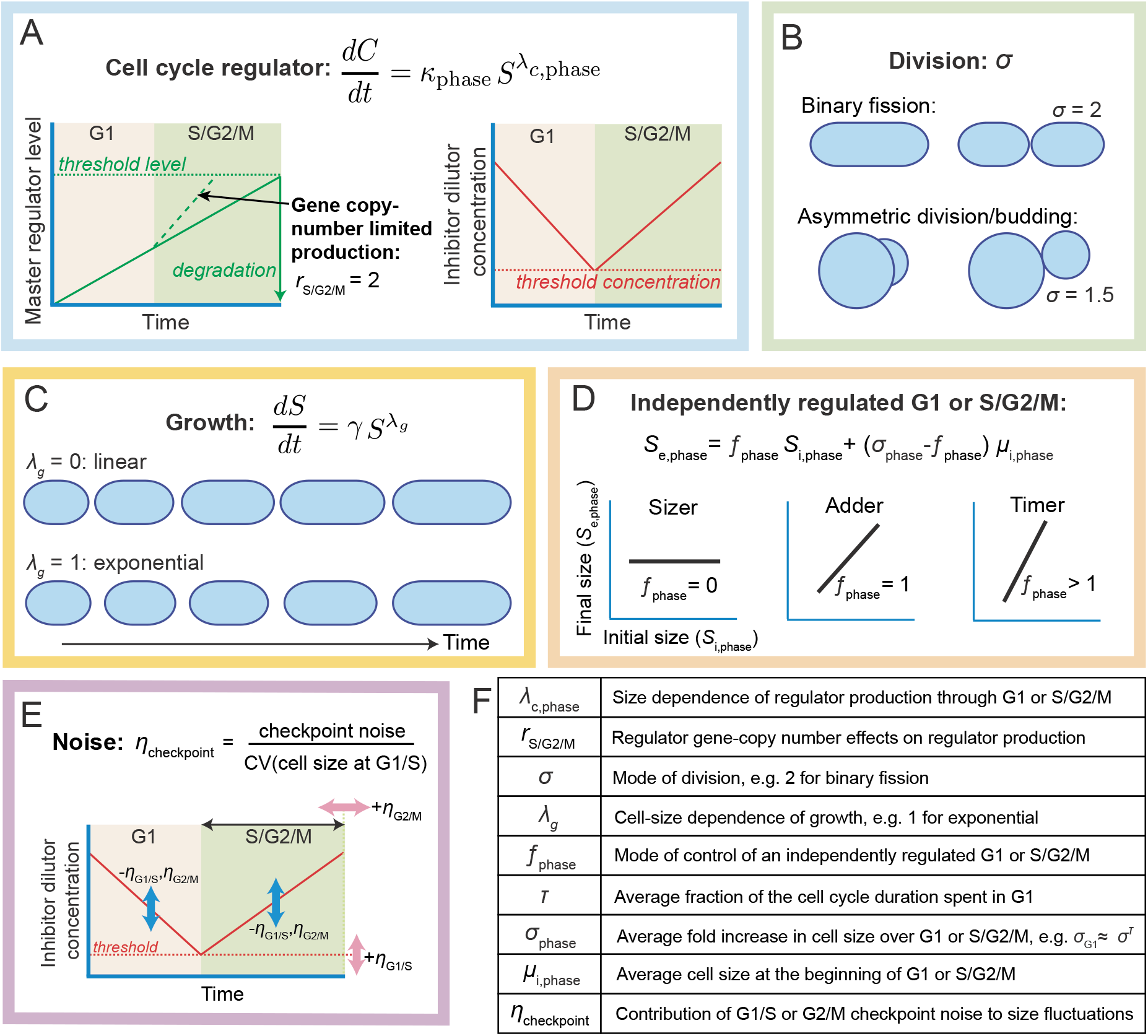
A general model of cell proliferation with two cell-cycle checkpoints. (A) The dependence of cell-cycle regulator production rate 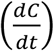 on cell size (*S*) is dictated by *λ*_*c*,phase_, with the phase corresponding to either G1 or S/G2/M. *λ*_*c*,phase_ = 1 corresponds to size-proportional production. Master regulators (left) are produced throughout one or both phases at a rate that may increase with gene-copy number (corresponding to *r*_S/G2/M_ = *k*_S/G2/M_/*k*_G1_ = 2) to trigger G1/S or G2/M at a threshold total intracellular level, and then are degraded. Inhibitor dilutors (right) are produced throughout one phase only (*k*_G1_ = 0) and then diluted out in the next, triggering G1/S or G2/M at a threshold concentration. (B) Cell division may occur through binary fission (*σ* = 2) or asymmetrically (*σ* < 2 or *σ* > 2). (C) Cell growth is exponential, linear, or intermediate (*λ_g_* = 1,0, or otherwise, respectively). (D) Cell cycle regulators can operate in combination with an independently regulated G1 or S/G2/M phase, meaning that the size at the end of the phase (*S*_i,phase_) depends only on the size at the beginning of the phase (*S*_i,phase_) and not on prior sizes, with the mode of regulation dictated by *f*_phase_: *f*_phase_ = 0,1 or depends on the growth behavior for critical size (“sizer”), adder, or timer regulation, respectively. *σ*_phase_ is the steady state average fold increase in cell size over the phase; *σ*_G1_ ≈ *σ^τ^*, where *τ* is the fraction of the cell cycle taken up by G1, and *σ*_S/G2/M_ ≈ *σ*^1−*τ*^. The average initial size *μ*_i,phase_ can be expressed in terms of other model parameters (Table S1, SI). (E) Cell-size fluctuations are due to noise in regulator dynamics and cell-cycle checkpoints. Noise effects are summarized by *η*_G1/S_ and *η*_G2/M_, corresponding to the noise in the G1/S and G2/M checkpoint criteria, respectively, divided by the coefficient of variation (CV) in G1/S size. For example, the G2/M checkpoint noise of S/G2/M timer control is generated by noise in the fixed time period between G1/S and G2/M (horizontal pink arrow); for G1/S inhibitor dilutors, noise in the minimum threshold concentration generates the G1/S checkpoint noise (vertical pink arrow). Noise sources that increase CV(cell size at G1/S) without affecting noise in the checkpoint criteria, for example noise in the production or dilution of the inhibitor (blue arrows), reduce *η*_G1/S_ and *η*_G2/M_. (F) Definitions of key parameters determining size-homeostasis behaviors.

In our model, cells divide into sisters with size-ratio 1:(*σ* – 1). Thus, binary fission and asymmetric division are accounted for by *σ* = 2 and *σ* ≠ 2, respectively (Fig. 1B), and at steady state cells increase their average birth size by an average factor *σ* over the cell cycle. The growth rate (*dS/dt*) can be cell-size dependent according to

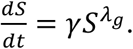

While many organisms grow exponentially (*λ_g_* = 1) (Di Talia et al., 2007; Osella et al., 2014; Soifer et al., 2016; Taheri-Araghi et al., 2015; Wang et al., 2010; Willis et al., 2016), there is some evidence of linear growth in certain regimes (*λ_g_* = 0) (Lin and Amir, 2018). *γ* sets the average timescale for growth; ln *σ /γ* is the average cell cycle duration for exponential growth (Fig. 1C).

We consider master regulators or inhibitor dilutors of G1/S or G2/M in combination with various phenomenological controls over S/G2/M or G1, respectively, including sizer, adder, or timer control, meaning that over the phase in question cells reach a critical size, add a fixed size increment, or a fixed time period elapses. Specifically, cell size at the end of the phase (*S*_e,phase_) is determined by cell size at the beginning of the phase (*S*_i,phase_) according to

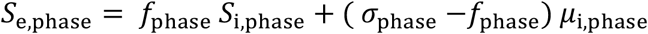

where *f*_phase_ is the mode of control (*f*_phase_ = 0, 1, or *σ*_phase_ for sizer, adder, or timer control and exponential growth, respectively), *σ*_phase_ > 1 is the average fold-size increase, and *μ*_i,phase_ is the average initial size at steady state (Fig. 1D). We call phases that follow this size-determination rule independently regulated. The average fraction of the cell cycle spent in G1 at steady state (*τ*, which equals the G1 duration × *σ*/ ln *γ* for exponential growth) and the mode of division (*σ*) determine *σ*_G1_ ≈ *σ^τ^* and *σ*_S/G2/M_ ≈ *σ*^1-*τ*^ because *σ* = *σ*_G1_ *σ*_S/G2/M_ (the approximations are exact for exponential growth, Methods). Average sizes at birth (*μ*_i,/G1_) and G1/S (*μ*_i,S/G2/M_) are determined by a combination of parameters governing the average regulator dynamics (*λ*_*c*,phase_, *k*_phase_) and threshold levels or concentrations, G1 duration (*τ*), growth type (*λ_g_, γ*), and division behavior (*σ*) (Table S1, SI). Cell-size fluctuations emerge from noise in regulator dynamics, noise in the critical levels or concentrations that trigger cell-cycle progression, and noise in sizer/adder/timer mechanisms. The impact of noise on size-homeostasis behavior is encapsulated by two parameters (*η*_G1/S_ and *η*_G2/M_; Methods and SI) according to

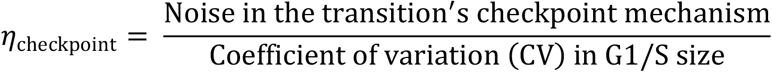

(Fig. 1E). For example, assuming typical values of the CV of G1/S size of ~13% (Cadart et al., 2018; Taheri-Araghi et al., 2015; Willis et al., 2016), a typical error of ~7% in a G1/S threshold concentration checkpoint gives *η*_G1/S_~0.5, and a typical error of ~20% in a S/G2/M timer checkpoint gives *η*_G2/M_~0.5 assuming exponential growth, binary fission, and *τ* = 0.5 (Methods and SI). Fig. 1F summarizes the parameters affecting size-homeostasis. In later sections, motivated by findings in *A. thaliana*, mammalian cells, and bacteria (Cadart et al., 2018; Ginzberg et al., 2018; Nordholt et al., 2019; Willis et al., 2016), growth and production rates are allowed to depend on cell birth size, and the threshold mechanisms for cell-cycle checkpoint progression are generalized.

Together, this model represents a broad framework for interrogating the requirements and molecular bases for cell-size homeostasis measurements.

### G1/S inhibitor dilutors combined with S/G2/M sizer or timer regulation fail to achieve size homeostasis in plausible cell proliferation scenarios

It is well known that master regulators of G2/M produced at a constant, size-independent rate throughout both phases fail to achieve size homeostasis in exponentially growing cells: a fixed time elapses between divisions, and cells multiply their birth size by a constant factor on average prior to division, so that cells born large get larger while small cells get smaller (Willis and Huang, 2017). We applied our model to identify other cell proliferation scenarios that fail to achieve G1/S or G2/M size homeostasis. We temporarily set noise sources to zero to determine whether homeostasis is lost independently of noise. To maintain homeostasis at steady G1/S and G2/M mean sizes, G1/S and G2/M size fluctuations must regress to their respective means, which requires that the absolute value of the slope between sizes at consecutive G1/S and G2/M transitions is <1 (Fig. S1). For example, the G2/M size-homeostasis requirement fails if a fixed time period *T* elapses between birth and division while cells grow exponentially:

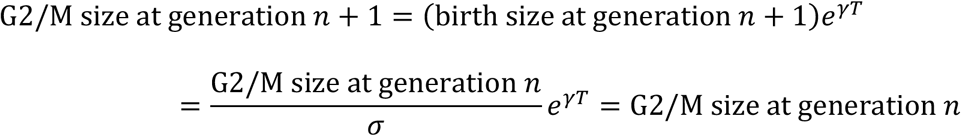

because at steady state *e^γT^* = *σ*; hence the slope between consecutive G2/M sizes is 1 and fluctuations do not decay to the mean. The loss of size homeostasis occurs despite the fixed cell cycle duration. Below, we readily identify more complex conditions that fail to regulate size by analytically deriving these slopes in terms of model parameters (Methods, SI).

In exponentially growing budding yeast, Whi5 executes inhibitor-dilutor control of G1/S, with Whi5 produced at a constant rate through S/G2/M, and an approximately fixed interval elapses between G1/S and G2/M (Schmoller et al., 2015). If S/G2/M is under timer control and growth is exponential, our model predicts that an inhibitor produced in proportion to cell size or growth (*λ*_*c*,S/G2/M_ ≥ 1; Fig. 1A,C) or with a stronger size-dependency (*λ*_*c*,S/G2/M_ ≥ 1) fails to regulate both G1/S and G2/M sizes (Fig. 2Ai,ii, Table S2). Single-cell trajectories with realistic noise levels illustrate this loss of size control as *λ*_*c*,S/G2/M_ varies from 0 (constant production rate) to 1 (proportional to size) (Fig. 2Bi-iv) while the G1 and S/G2/M durations and thus the ordering of G1/S and G2/M are maintained so cells contain the correct number of genome copies. By contrast, if S/G2/M were under adder control, size homeostasis is achieved for *λ*_*c*,S/G2/M_ = 0 and 1 (Fig. 2Aiii,iv).

**Figure 2:**
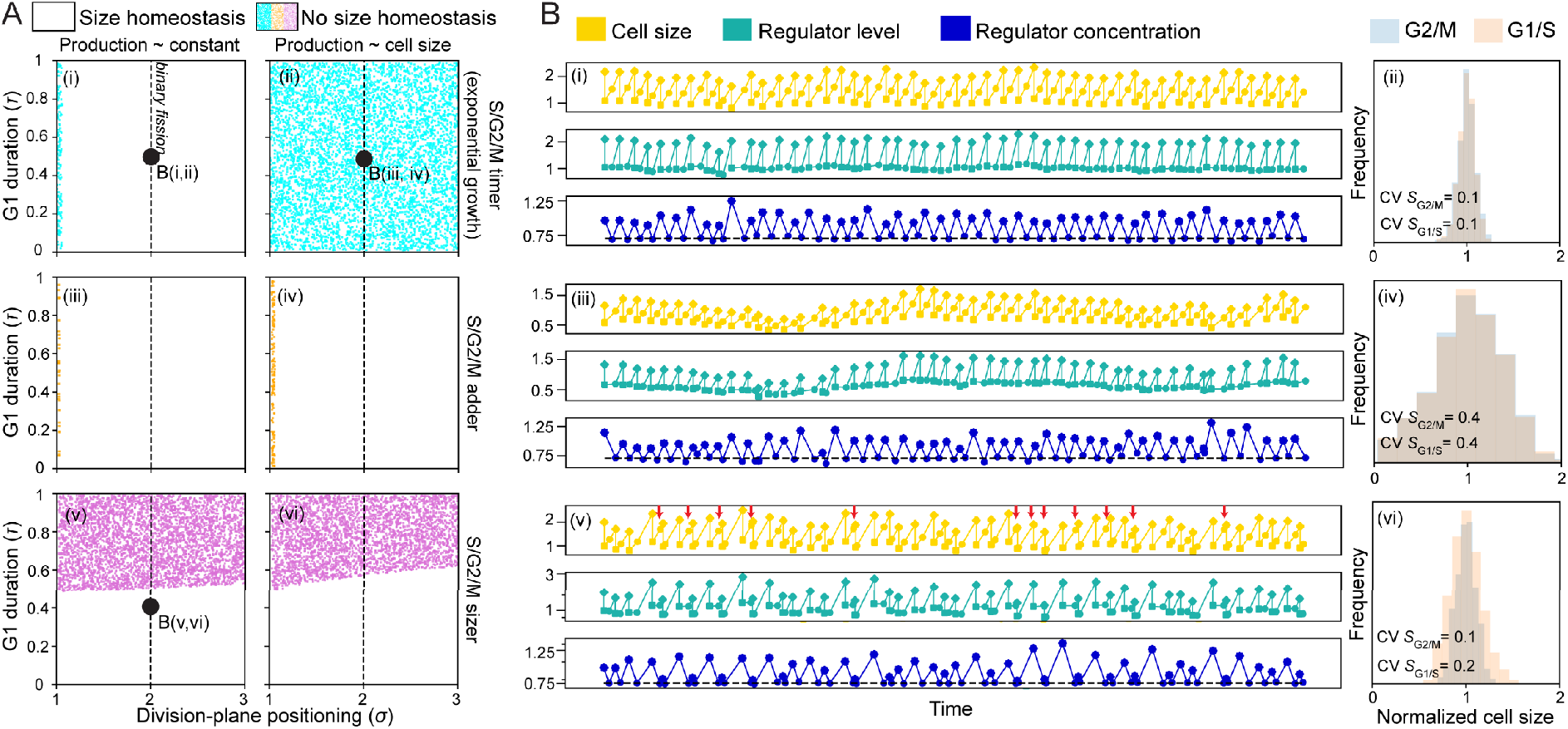
G1/S inhibitor dilutors combined with S/G2/M sizer or timer regulation can fail to achieve size homeostasis. (A) Colored regions indicate where G1/S size homeostasis is lost (as defined by analytical results for exponentially growing cells (*λ_g_* = 1) with G1/S inhibitor-dilution control in which the absolute value of the slope of G1/S size in generation *n* vs. generation *n*+1 is >0.95) for a range of division-plane positions (*σ*; x-axis), G1 durations (*τ*; y-axis), different modes of S/G2/M regulation (rows), and inhibitor production being constant (left) or proportional to cell size (right). If S/G2/M is under timer regulation, size homeostasis is predicted to be achieved if production is not size-dependent (*λ*_*c*,S/G2/M_ = 0) (i) but generally lost if inhibitor production is proportional to cell size (*λ*_*c*,S/G2/M_ = 1) (ii). Regardless of growth and production patterns, G1/S inhibitor dilutors are also predicted to be incompatible with the combination of G2/M sizer regulation and long G1 durations (*τ* ≥ 0.5) (v,vi). S/G2/M adder regulation generally maintains size homeostasis (iii,iv). Black ccircles correspond to single cell lineages simulated in (B). (B) Simulations of single cell lineages with realistic noise levels confirm analytical results. (i,ii) Cell size, inhibitor level, and inhibitor concentration control are achieved for a budding yeast-like inhibitor dilutor (*λ*_*c*,S/G2/M_ = 0, *σ* = 2, *τ* = 0.5) in exponentially growing cells (*λ_g_* = 1) with S/G2/M timer regulation and realistic noise levels (coefficient of variation (CV) of G2/M size ≈ 0.1, produced by noise terms *ξ*_C,G1_, *ξ*_C,S/G2/M_, *ξ*_G1/S_, *ξ*_G2/M_ = 0.03, Methods), as demonstrated by a cell lineage (left) and the corresponding G1/S and G2/M size distributions (right). In the cell lineage, squares, circles, and diamonds denote birth, G1/S, and G2/M (division), respectively. (iii, iv) For the same noise levels as in (i,ii), cell-size control was nearly lost when the inhibitor’s production was *λ*_*c*,S/G2/M_ = 0.95 ≈ *λ_g_* = 1 (the same size dependence as growth). The size distributions (iv) show that transitions frequently occur at ~10% or ~200% of the usual size. (v,vi) G2/M sizer control causes loss of G1/S size homeostasis if the G1 duration is long (*σ* = 2, *τ* = 0.4 are at the boundary of G1/S size homeostasis, as indicated in Fig. 2A(v); CV of G2/M size ≈ 0.1, produced by *ξ*_C,G1_, *ξ*_C,S/G2/M_, *ξ*_G1/S_ = 0.01, *ξ*_G2/M_ = 0.1, Methods). Red arrows point to cell cycles where G1/S occurred near the end of the cell cycle; in the following cell cycle, G1/S tends to occur early. The G1/S size distribution (vi) shows G1/S frequently occurs at ~60% or ~140% of the average G1/S size, and therefore at ~0.6 *σ*_G1_ = 0.6 2^*τ*^ = 0.8 or ~1.4 *σ*_G1_ = 1.4 2^*τ*^ = 1.8 of the average birth size.

Regardless of the type of growth and inhibitor production, the inhibitor threshold concentration for G1/S, and the division pattern, we also found that G1/S inhibitor dilutors are incompatible with G2/M sizer mechanisms and long G1 durations (*τ* ≥ 0.5): for *τ* ≈ 0.5, fluctuations below the average G1/S size are overcompensated for in the subsequent generation (Fig. 2Av,vi), consistent with analytical predictions of a slope between consecutive G1/S sizes ≤-1 (Fig. S1, Table S2), so G1/S size homeostasis is lost and transitions frequently alternate between the beginning and end of the cell cycle (arrows in Fig. 2Bv,vi). Analogous results apply for inhibitor dilutors that trigger G2/M rather than G1/S with timer/adder/sizer control over G1 (SI). The model’s generality and analytical tractability enabled these results, which demonstrate how cell proliferation scenarios that are *a priori* biologically plausible necessarily fail to achieve size homeostasis and thus can be ruled out.

### Master regulators can lose cell-size control when production is gene copy-number limited or strongly size-dependent, or when a threshold concentration triggers phase progression

CDK1-cyclin and DnaA/FtsZ cell-cycle regulation are thought to be widely conserved in eukaryotes and bacteria, respectively, but details of the regulators’ dynamics and other proliferation factors have yet to be quantified in nearly all species. Motivated by CDK1-cyclin and DnaA/FtsZ, we focused on G2/M and G1/S master regulators produced through both phases (“two-phase” master regulators) and applied our model to identify limits on proliferation dynamics imposed by size homeostasis.

For exponentially growing cells, when regulator production is gene copy-number limited (*r*_S/G2/M_ = 2; Fig. 1A), G2/M two-phase master regulators produced at a size-independent rate (*λ*_*c*,G1/S_ = *λ*_c,S/G2/M_ = 0) fail to execute size homeostasis for G1 timer control, because then a fixed time elapses over the cell cycle which is incompatible with exponential growth regardless of other parameters (Fig. 3Ai). However, we found that size homeostasis is restored for G1 sizer or adder control (Fig. 3Aii,iii, Table S2) largely regardless of the average G1 duration (*τ*). By contrast, G1/S two-phase master regulators that are gene copy-number limited are incompatible with G2/M sizer regulation: this combination fails to implement G1/S size homeostasis for binary fission or asymmetric division (*σ* ≤ 2) (Fig. 3B, Table S2). This finding depends on the increased rate of regulator production during S/G2/M due to gene-copy number doubling and, conditional upon no growth rate (*γ*) increase during S/G2/M, holds regardless of the average G1 duration and the size-dependencies of production and growth (*λ_g_* and *λ*_*c*,phase_, assuming *λ*_*c*,G1_ = *λ*_*c*,S/G2/M_) (SI). Here, the loss of G1/S size homeostasis while G2/M size homeostasis is enforced implies that the proper ordering of G1/S followed by G2/M is also lost (Fig. S2). G1/S two-phase master regulators produced at a strongly size-dependent rate throughout the cell cycle without being gene copy-number limited (*λ_c_* = *λ*_*c*,G1_ = *λ*_*c*,S/G2/M_ ≥ 2 + *λ_g_*, *r*_S/G2/M_ = 1) fail to execute G1/S size homeostasis for S/G2/M timer regulation combined with binary fission or asymmetric division (*σ* ≥ 2) and long S/G2/M durations (1 – *τ* ≥ 0.6) (Fig. 3Ci, iv). For 2 *λ_c_* = 2 + *λ_g_* but not *λ_c_* = 3 + *λ_g_*, size homeostasis is restored by S/G2/M adder or sizer regulation (Fig. 3C, Table S2). Thus, under exponential growth, G1/S two-phase master regulators such as DnaA robustly achieve size homeostasis only when 0 < *λ_c_* < 3.

While size homeostasis is achieved by two-phase master regulators produced from an initial level of zero in proportion to growth (*λ*_*c*,G1_ = *λ*_*c*,S/G2/M_ = *λ_g_*, *r*_S/G2/M_ = 1) to trigger phase progression at a threshold level (Amir, 2014), we found that if instead progression is triggered at a critical concentration (or a local threshold density in an intracellular region that scales proportionally with cell size), size homeostasis is generally lost (Fig. 3D, SI). This mechanism fails because a threshold concentration means that cell size at the checkpoint is proportional to the regulator’s level, which is proportional to the added size since production is proportional to growth. Thus, cell size at the checkpoint is proportional to the added size, and ultimately cells multiply their birth size by a constant factor on average prior to division, so there is no negative feedback on size fluctuations. However, if the size-dependence of regulator production exceeds that of growth (*λ*_*c*,phase_ > *λ_g_*), size homeostasis can be restored (SI). Thus, to maintain size homeostasis in bacteria, the division-initiating FtsZ midcell bands must not increase in width proportionally as the cell grows.

In all cases above, analytical predictions were confirmed by simulations of single-cell trajectories with realistic size fluctuations (Fig. 2B, 3D, S2). A recurring theme is that switching to adder regulation of S/G2/M or G1 restored size homeostasis when sizer or timer regulation failed (white regions in Fig. 2A, 3A,B,C). Thus, these scenarios illustrate robustness of adder regulation, regardless of its molecular origin.

### Connecting cell-cycle regulation mechanisms to linear regression slopes between birth and division sizes

When taking noise into account, we can predict how regulatory factors affect size-homeostasis behaviors at steady state by deriving linear regression slopes between birth and division sizes and among other size variables (Methods). We present particular scenarios motivated by experiment (see SI for the general case).

At least in *E. coli*, DnaA appears to operate as a two-phase master regulator between consecutive G1/Ss (Si et al., 2019), while in slow growth conditions S/G2/M is independently regulated by a timer or adder mechanism or a mechanism whereby both a minimum time period elapses and a minimum size increment is added between divisions (Logsdon et al., 2017; Micali et al., 2018b). Then, the linear regression slope of birth vs. division size is

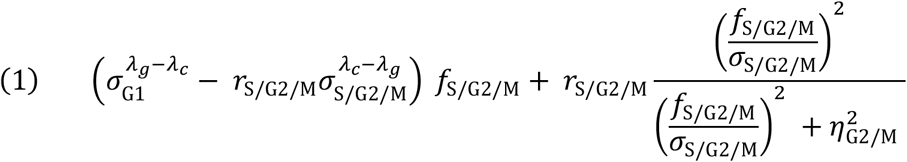

where *λ_c_* = *λ*_*c*,G1_ = *λ*_*c*,S/G2/M_ represents the regulator’s size-dependent production, which we assume to be constant throughout the cell cycle, and recalling *σ*_G1_ ≈ *σ^τ^* ≈ *σ/σ*_S/G2/M_ (Fig. 1F, Methods). From Eq. 1, and in agreement with other studies (Amir, 2014; Ho and Amir, 2015), a production rate that is proportional to growth (*λ*_*c*_ = *λ_g_*, *r*_S/G2/M_ = 1) combined with low G2/M checkpoint noise vs. other sources of cell size noise (*η*_G2/M_ ≪ 1) and supra-sizer control of G2/M (*f*_S/G2/M_ > 0) gives a slope ~1, which means that apparent adder behavior (slope=1) is predicted regardless of further constraints on the S/G2/M control mechanism (*f*_S/G2/M_) (Fig. 4A). *e*_G2/M_ is small when cell size fluctuations are primarily from noise sources other than the G2/M checkpoint, for example, from fluctuations in regulator production (Fig. 1E). Under exponential noiseless growth, a typical error *e*_G2/M_ in a S/G2/M timer checkpoint (CV of S/G2/M duration, SI) dictates that *η*_G2/M_ = *e*_G1/M_ ln *σ*_S/G2/M_ /(CV of G1/S size), so for *e*_G2/M_ = 10%, *σ*_S/G2/M_ ≈ *σ*_1-*τ*_ ≈ 1.4, and a CV of G1/S size = 13%, then 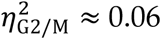 has a nominal effect on the slope in Eq. 1 (Fig. 4A).

**Figure 3:**
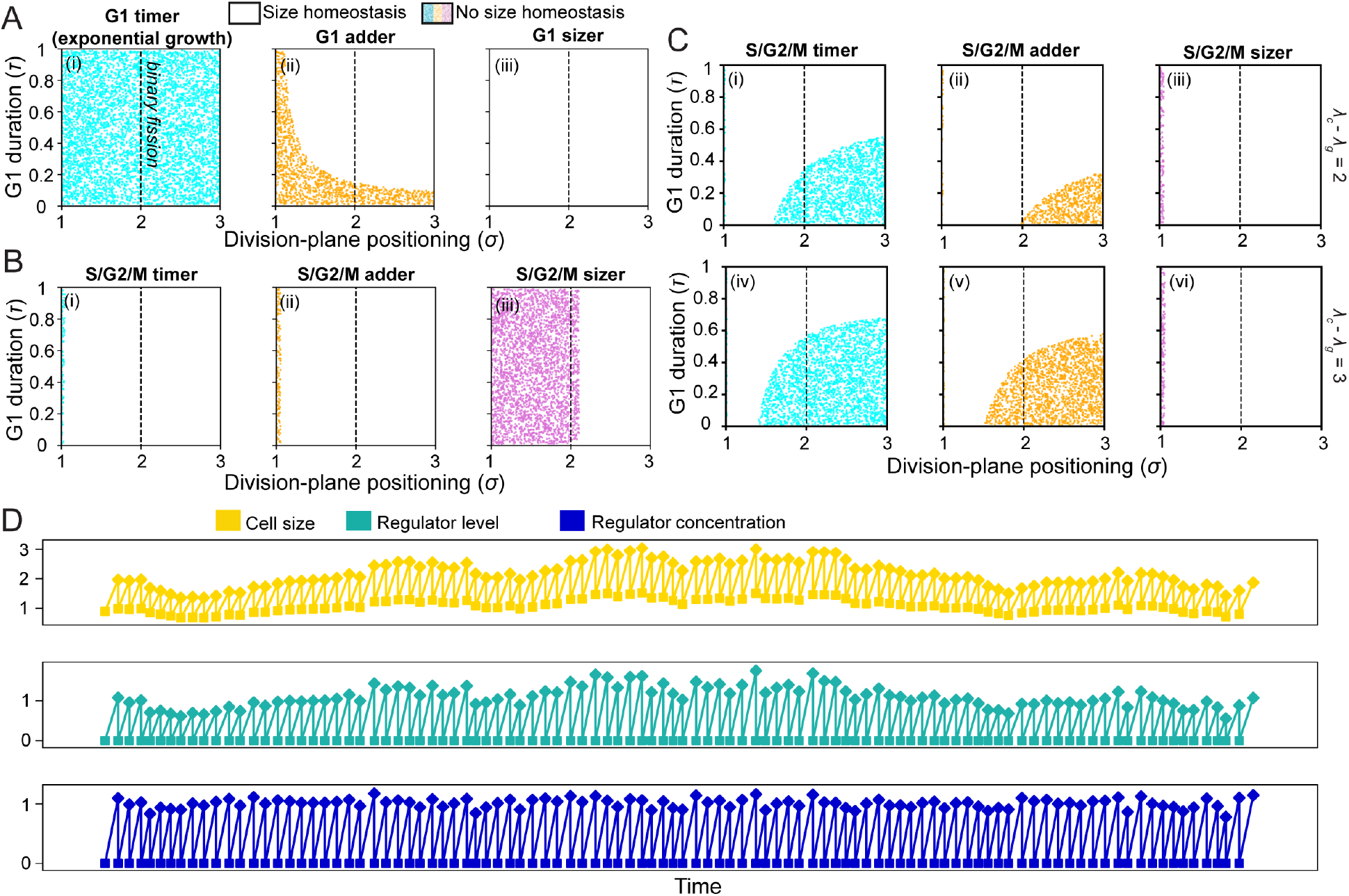
Master regulators can lose cell-size control when production is gene copy-number limited or strongly size-dependent, or a threshold concentration rather than level triggers phase progression. (A) G2/M two-phase master regulators with size-independent production rates (*Λ*_*c*,G1_ = *λ*_*c*,S/G2/M_ =0) can achieve cell-size regulation in exponentially growing cells (*Λ_g_* = 1) if their production rate is gene copy-number limited (*r*_S/G2/M_ = 2) while G1 is under sizer (iii) or adder (ii) regulation, but not timer regulation (i). Colored regions indicate parameters in which G2/M size homeostasis is lost. (B) G1/S two-phase master regulators with gene copy-number limited production that have the same size-dependencies as growth (*λ*_*c*,G1_ = *λ*_*c*,S/G2/M_ = *λ_g_*) achieve size homeostasis for S/G2/M adder (ii) and timer (i) regulation, but are incompatible with G2/M sizer regulation when *σ* ≤ 2 (iii). Colored regions indicate parameters in which G1/S size homeostasis is lost. (C) G1/S two-phase master regulators with strongly size-dependent production (*λ_c_* – *λ_g_* = 2 (top) or 3 (bottom) where *λ_c_* = *λ*_*c*,G1_ = *λ*_*c*,S/G2/M_) and no gene-copy number effects fail to achieve size homeostasis for S/G2/M timer (i,iv) and adder (ii,v) regulation for long S/G2/M durations (1 – *τ* > 0.5) and binary fission (*σ* ≥ 2). Colored regions indicate parameters in which G1/S size homeostasis is lost. (D) The size (i), regulator level (ii), and regulator concentration (iii) trajectories of a cell lineage governed by a two-phase master regulator that is produced from an initial level of zero and triggers G2/M at a noisy threshold concentration instead of a noisy threshold total intracellular level, with size-dependent production *λ_c_* = *λ*_*c*,G1_ = *λ*_*c*,S/G2/M_ = 1.05 similar to the size-dependence of growth *λ_g_* = 1. Cell size fluctuates dramatically as *λ_c_* approaches *λ_g_*, and size control is lost in the limit that *λ_c_* = *λ_g_*. Squares and diamonds denote birth and G2/M (division), respectively.

**Figure 4:**
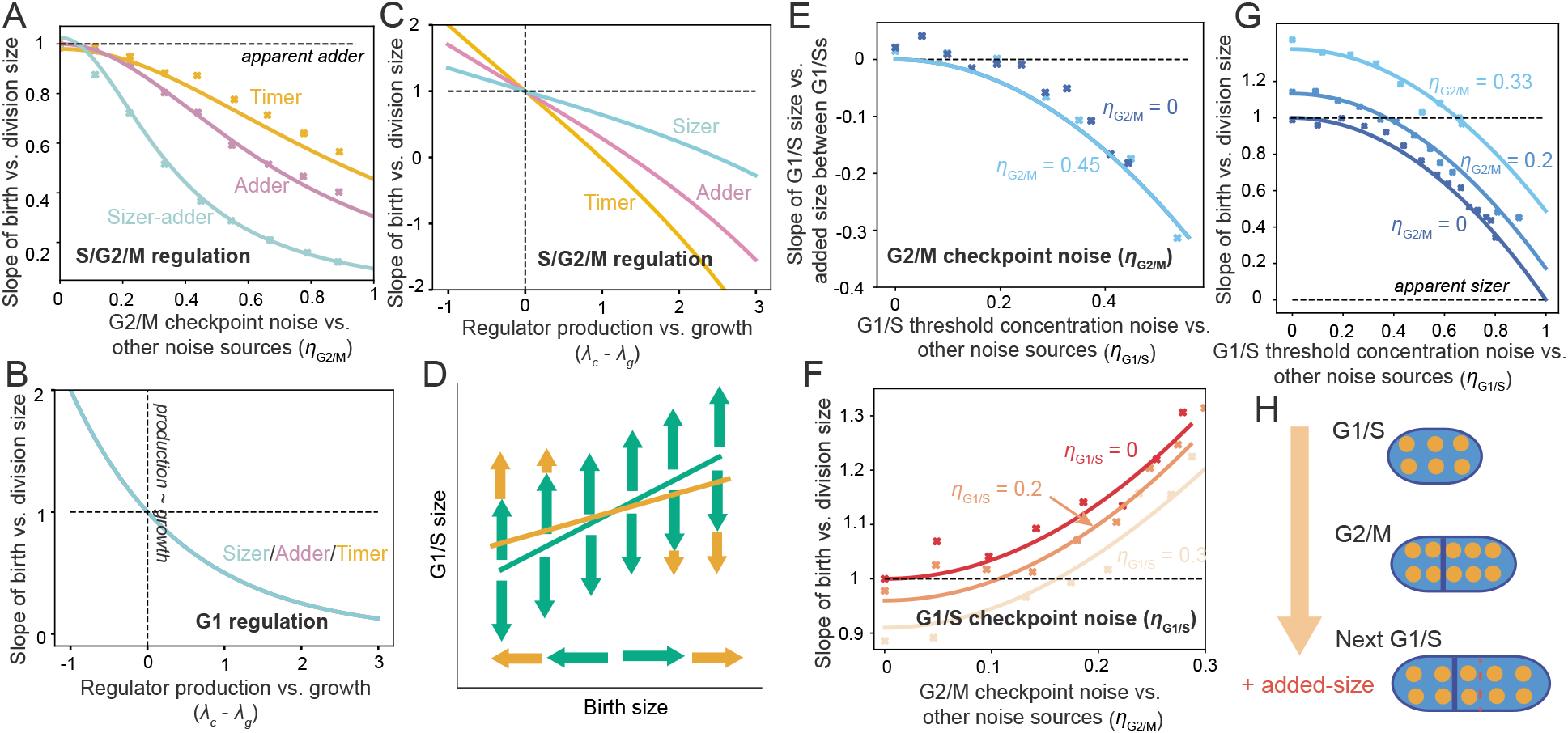
Cell-cycle regulation mechanisms can produce various apparent size homeostasis behaviors depending on factors such as noise and regulator production. (A) G1/S two-phase master regulators that are produced in proportion to growth (*λ_c_* = *λ_g_, r*_S/G2/M_ = 1 in Eq. 1) exhibit a linear regression slope of ~1 between birth and division regardless of the S/G2/M regulatory mode, representing apparent adder behavior, as long as the G2/M checkpoint makes a weak contribution to size fluctuations (*η*_G2/M_ ≪ 1). Analytical approximations (solid lines, SI) agree with exact simulations (symbols) for realistic noise levels (CV of G1/S size ≈ CV of G2/M size ≈ 0.1). In the plots, *σ*_S/G2/M_ = 1.4 was assumed. (B) G2/M two-phase master regulators produce slopes ~0 and thus near-sizer behaviors when production is strongly size-dependent with no gene copy-number effects (*λ_c_* > *λ_g_, r*_S/G2/M_ = 1 in Eq. 2) regardless of the G1 regulatory mode (the curves for G1 timer, adder, and sizer regulation are all identical) and noise levels. Here, binary fission (*σ* = 2) was assumed. (C) By contrast, G1/S two-phase master regulators tend to produce negative slopes for strongly size-dependent production with no gene copy-number effects (*λ_c_* > *λ_g_, r*_S/G2/M_ = 1 in Eq. 1). Here, weak G2/M checkpoint noise (*η*_G2/M_ ≪ 1), *σ*_S/G2/M_ = 1.4, and *σ* = 2 were assumed. (D) Among G1/S two-phase master regulators, relatively high noise in the G2/M checkpoint mechanism results in small or large cells at birth with low or high regulator levels, respectively. Small cells then grow more and thus produce more regulator to achieve the surplus regulator level for G1/S progression (orange arrows). Noise sources that impact the regulator’s production are uncoupled from initial birth size, and thus do not affect the size homeostasis behavior between birth and G1/S (green arrows). (E-G) Depending on which processes make the largest contributions to cell-size fluctuations, a budding yeast-like G1/S inhibitor-dilutor with S/G2/M timer regulation can generate a variety of size homeostasis behaviors as represented by the linear regression slopes among size variables: if noise in inhibitor dynamics primarily generates G1/S size fluctuations (*η*_G1/S_ ≪ 1, *η*_G2/M_ ≪ 1 in Eq. 4), apparent adder behavior results between consecutive G1/Ss (E) and birth and division (F,G). *η*_G1/S_ ≪ 1 requires stringent control of G1/S. (H) Addition of a constant amount of inhibitor (orange spots) over S/G2/M achieves adder size homeostasis between consecutive G1/Ss, because added size scales with amount of inhibitor produced owing to the threshold inhibitor concentration requirement for G1/S progression.

In engineered fission yeast, CDK1-cyclin is produced through G1 and S/G2/M to trigger G2/M at a threshold prior to degradation (Coudreuse and Nurse, 2010), while in *E. coli* FtsZ accumulates at midcell up to a threshold level to trigger G2/M or division (Si et al., 2019); in both cases, G2/M is likely controlled by a two-phase master regulator. The corresponding linear regression slope of birth vs. division size is

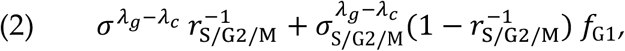

producing adder behavior when production and growth rates are proportional throughout the cell cycle (*λ_c_* = *λ_g_*; *r*_S/G2/M_ = 1) regardless of other parameters. For strongly size-dependent production (*λ_c_* > *λ_g_*), from Eq. 2, G2/M master regulators asymptote to slopes of ~0 and thus to apparent sizer behaviors regardless of other parameters (Fig. 4B). By contrast, from Eq. 1, for *λ_c_* ≈ *λ_g_* + 2, G1/S two-phase master regulators tend to produce strongly negative slopes corresponding to oscillatory size behaviors where the average is overshot then undershot (Fig. 4C).

Eq. 1 indicates that high G2/M checkpoint noise 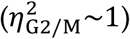 invariably reduces the slope of G1/S two-phase master regulators, an effect that can be understood qualitatively. High *η*_G2/M_ and a non-noisy coupling between growth and regulator dynamics entail that cells born small contain less master regulator and therefore must produce more regulator over G1 and correspondingly grow more to achieve the threshold level for G1/S (Fig. 4D). For timer/adder/intermediate sizer-adder regulation of S/G2/M, any compensatory growth over G1 is inherited as a positive fluctuation in G2/M-added size. Thus, the slope between birth and division is reduced by noisy G2/M checkpoint control, and this effect can be masked by other processes that contribute to G1/S size fluctuations without coupling G1/S size to birth size, such as noisy production of the G1/S regulator. By contrast, the slope in Eq. 2 is unaffected by noise levels (SI): noise impacts the size-homeostasis behaviors between birth and division of G1/S regulators because the production and persistence of the regulator through G2, mitosis, and division correlates birth-size fluctuations with fluctuations in birth-regulator levels; G2/M regulators are degraded or used up prior to birth, so there is no mechanism to generate such a correlation.

These findings showcase the interplay of factors contributing to size-homeostasis statistics and opposing predictions for G1/S vs. G2/M two-phase master regulators concerning the effects of noise and size-dependent production.

### Noise levels dictate whether inhibitor dilutors achieve adder-like, sizer-like, or supra-adder behaviors

In budding yeast, various studies have identified adder-like behavior between birth and division (Chandler-Brown et al., 2017; Di Talia et al., 2007; Soifer et al., 2016). However, deletion of *CLN3*, which leads to prolonged G1 and increased average size (Cross, 1988), or an additional copy of *WHI5* caused behavior closer to sizer for cells that were born small (Chandler-Brown et al., 2017). Experiment and theory have shown that in the absence of noise, the observed combination of constant production of the inhibitor dilutor *WHI5* controlling G1/S, exponential cell growth, and timer control of S/G2/M can result in apparent adder behavior (Chandler-Brown et al., 2017; Di Talia et al., 2007; Heldt et al., 2018; Schmoller et al., 2015; Soifer et al., 2016), despite the lack of a molecular adder underpinning the phenomenological behavior between birth and division. Moreover, noise was suggested to have the potential to disrupt adder behavior (Barber et al., 2017).

We applied our model to determine how noise and gene copy-number effects impact size-homeostasis behaviors of budding yeast-like G1/S inhibitor dilutors. The linear regression slope of G1/S size vs. overall added size until the next G1/S (ignoring the intervening division) most easily discerns the size-homeostasis behavior between consecutive G1/Ss: a slope of 0 indicates adder behavior. For exponential growth, S/G2/M timer control, and constant inhibitor production through S/G2/M as in budding yeast, the linear regression slope of G1/S size vs. overall added size until the next G1/S is

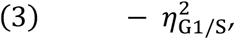

and the slope of birth vs. division size is

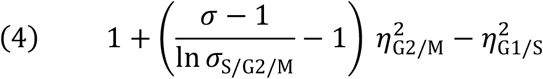

(Methods). From Eqs. 3 and 4, noise contributions affect size-homeostasis behaviors (Fig. 4E-G) while the inhibitor’s production rate (*k*_S/G2/M_) and threshold concentration for G1/S, which are feasibly altered by *WHI5* and *CLN3* copy numbers, have no impact. Apparent adder behaviors, corresponding to slopes of ~0 and ~1 in Eqs. 3 and 4, respectively, are observed only for relatively stringent G1/S checkpoint control: for a typical error in the inhibitor’s G1/S threshold concentration of *e*_G1/M_ = 10%, *η*_G1/S_ = *e*_G1/S_/CV(G1/S size) ≈ 0.8 (assuming CV(G1/S size) ≈ 13%), so the slopes in Eqs. 3 and 4 are perturbed by –0.8^2^ = –0.64 producing sub-adder behaviors (Fig. 4E,G); whereas for a comparable typical error in the S/G2/M timer checkpoint of *e*_G2/M_ = 10%, then *η*_G2/M_ = *e*_G2/M_ ln *σ*_S/G2/M_ /CV(G1/S size) = 0.25 (assuming *σ* = 2 and *σ*_S/G2/M_ = *σ*^1–*τ*^ = 2^1.5^ = 1.4) and the slope in Eq. 4 is perturbed considerably less (by only +0.12, Fig. 4F) and near-adder behavior results. Among inhibitor dilutors, similar to G1/S two-phase master regulators (Eq. 1), adder behaviors require that size fluctuations are primarily from noise sources other than cell-cycle checkpoint mechanisms, such as fluctuations in inhibitor production and dilution.

The size-homeostasis behaviors in Eqs. 3 and 4 can be understood qualitatively. The key feature that results in apparent adder behavior between G1/Ss is the constant amount of inhibitor produced over S/G2/M: the critical concentration at G1/S means that every inhibitor molecule corresponds to a small unit size *s*_0_, so if *n* inhibitor molecules are produced over S/G2/M, the cell adds a size *n* × *s*_0_ to achieve the threshold concentration at the next G1/S (Fig. 4H) (Schmoller et al., 2015). Noise in either the inhibitor dynamics or the S/G2/M timer phase alters the number of inhibitor molecules produced during S/G2/M, and therefore causes the added size between consecutive G1/Ss to fluctuate, but this fluctuation is independent of the initial G1/S size. By contrast, fluctuations in the threshold concentration (the G1/S checkpoint) entail the opposite scenario; a high inhibitor threshold concentration corresponds to small cells at G1/S, so small cells must grow more than the average to dilute out the surplus inhibitor, leading to sub-adder behavior between G1/Ss. If G1/S size fluctuations arise entirely from G1/S checkpoint noise, then the relatively small noise in inhibitor production and S/G2/M interval means that daughter cells inherit a constant amount of inhibitor at birth, and the threshold concentration necessary for G1/S translates into a threshold cell size, resulting in apparent sizer regulation. These arguments apply to any inhibitor production rate and S/G2/M regulatory mode that together produce a constant amount of inhibitor during S/G2/M regardless of growth pattern and G1 duration, under the assumption of no inhibitor degradation following G1/S. Rapid inhibitor degradation following G1/S would produce sizer behavior between G1/Ss, since regardless of noise levels, the G1 regulator level would be uncoupled from the preceding G1/S size.

These findings indicate that G1/S inhibitor dilutors can exhibit adder, supra-adder, or sizer behavior depending on which processes make the primary contributions to cellsize fluctuations (Schmoller et al., 2015). Stringent control of the inhibitor’s threshold concentration for G1/S is necessary for apparent adder behavior, and noise in this control may lead to sizer-like behaviors.

### Deviations in regulator localization patterns cause supra-adder behaviors

Cell-cycle regulators can accumulate in a particular region of the cell, triggering checkpoint progression at a local threshold density. For example, in fission yeast the cell cycle regulator cdr2 accumulates at midcell to trigger division at a local threshold density (Facchetti et al., 2019; Pan et al., 2014). In this case, the threshold density mechanism effects a total intracellular threshold level criterion, because fission yeast grow as rods and the cdr2 localization region at midcell does not scale with size. Division in rod-shaped *E. coli* is regulated in an analogous way by FtsZ (Shi et al., 2017; Si et al., 2019). By contrast, the midcell diameters of coccoid bacteria and plant stem cells are not constant but increase with cell size (Willis et al., 2016), and in general cellular geometries and growth patterns vary among organisms.

To address the implications of heterogeneous localization, we derived the size-homeostasis behaviors of G2/M two-phase master regulators that accumulate to a threshold density within a cellular region that increases with cell size as ~*S*^λ_*T*_^. The region grows in proportion to size if *λ_T_* = 1 (as do most nuclei), and (depending on geometry) with surface area or midcell perimeter if *λ_T_* ≈ 2/3 or 1/3. Then, assuming *λ_c_* = *λ*_*c*,G1_ = *λ*_*c*,S/G2/M_ and *r*_S/G2/M_ = 1 (Fig. 1F), the linear regression slope of birth vs. division size is

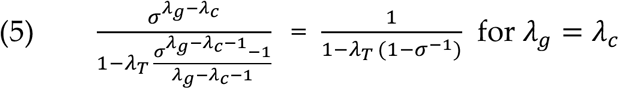

(SI). From Eq. 5, because 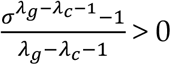, localization to a region that increases with size (*λ_T_* > 0) invariably increases the slope (Fig. 5A), predicting supra-adder behavior when growth and regulator production are proportional (*λ_g_* = *λ_c_*) and intermediate sizeradder behavior when production is strongly size-dependent (*λ_c_* ≫ 1). Adder behavior is achieved when a threshold concentration rather than level of regulator triggers G2/M (*λ_T_* = 1 vs. *λ_T_* = 0 in Eq. 5) under the requirements that *λ_g_* < *λ_c_* and 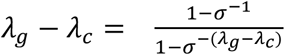 (≈ −0.7 for *σ* = 2) (Fig. 5B). We have been unable to identify plausible mechanisms producing this relation, suggesting that threshold concentration mechanisms are unlikely to underlie adder behavior.

**Figure 5:**
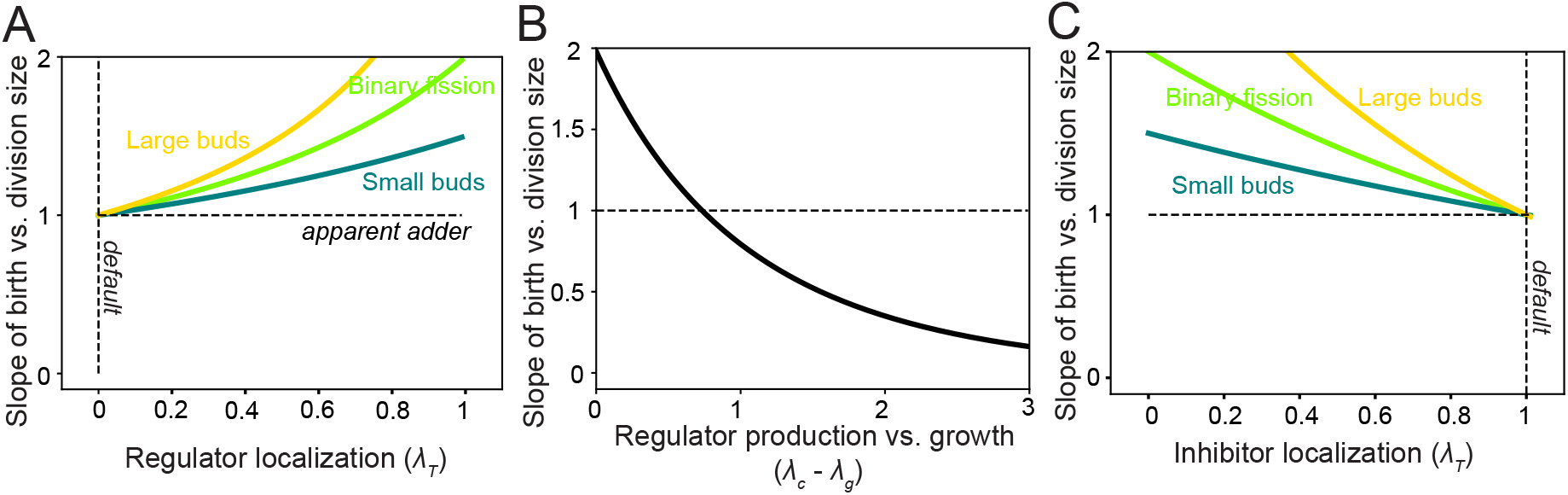
Deviations in regulator localization patterns cause supra-adder behaviors. (A) G2/M two-phase master regulators that localize to a region that scales with cell size as *S^λ_T_^* to trigger checkpoint progression at a threshold density produce a slope between birth and division size of 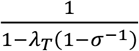, generating supra-adder behavior as *λ_T_* deviates from zero (the model’s default scenario), assuming regulator production is proportional to growth (*λ_g_* = *λ_c_* in Eq. 5). (Yellow, light green, or dark green lines are for *σ* = 3, 2, or 1.5). (B) For a G2/M threshold concentration rather than threshold level, the slope between birth and division size is 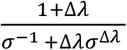 where Δ*λ* = *λ_c_* − *λ_g_* (assuming *r*_S/G2/M_ = 1), which gives adder behavior when Δ*λ* ≈ 0.7 for binary fission (*σ* =2). (C) Budding yeast-like G1/S inhibitor dilutors that localize to a region that scales with cell size as *S*^*λ_T_*^ to trigger phase progression at a threshold density produce a slope between birth and division size of *σ*^1−*λ_T_*^, generating supra-adder behavior as *λ_T_* deviates from one (the model’s default scenario), assuming a fixed average amount of inhibitor is produced during S/G2/M and noisy inhibitor dynamics (*η*_G1/S_, *η*_G2/M_ ≪ 1 in Eq. 6). (Yellow, light green, or dark green lines are for *σ* = 3, 2, or 1.5, respectively).

Cell cycle inhibitors can also localize to subcellular regions, such as the localization of Whi5 to the nucleus for part of G1 in budding yeast (Di Talia et al., 2007). If an inhibitor dilutor localizes to a region that increases with cell size as ~*S*^λ_T_^ and triggers G1/S upon reaching a threshold density within that region, the linear regression slope of birth vs. division size is

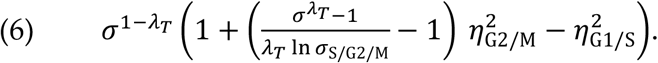

From Eq. 6, for stringent checkpoint control or noisy inhibitor dynamics (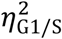 and 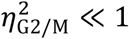), localization to any region that does not scale proportionally with size (*λ_T_* < 1) increases the slope (Fig. 5C), again predicting supra-adder behavior (SI).

In summary, deviations in regulator localization patterns from the default scenarios (localization to a region that is proportional to size in inhibitor dilutors or independent of size in master regulators) increases the linear regression slope between birth and division sizes, tending to generate supra-adder behavior.

### Apparent adder regulation between birth and division can be achieved with non-adder regulation over G1 and S/G2/M via independently regulated G1 or S/G2/M phases

Mammalian cells and budding yeast behave as apparent adders or near-adders between birth and division. However, the linear regression slopes between birth and G1/S sizes (*l*_G1_), and G1/S and G2/M or division sizes (*l*_S/G2/M_), are significantly different from 1 (Cadart et al., 2018; Chandler-Brown et al., 2017; Schmoller et al., 2015), indicating deviations from adder behavior over G1 and S/G2/M individually. In some strains, the G1 size-homeostasis behavior is sub-adder, compensating for a supra-adder timer mode of S/G2/M regulation to achieve an overall adder between birth and division (Cadart et al., 2018).

To consider how cell proliferation mechanisms can give rise to G1 compensatory behaviors, it was important to establish the consequences of independent regulation of a phase (Fig. 1D). Eukaryotic cyclin-like regulators that are produced through only one phase and degraded outside of this phase are likely to produce independently regulated phases, because only size fluctuations at the beginning of the phase impact the regulator’s dynamics and thus size fluctuations at the end of the phase (Fig. 6A). For independently regulated phases with control mode *f*_phase_, the linear regression slope between cell sizes at the beginning and end of the phase is *l*_phase_ = *f*_phase_ (Methods). If S/G2/M is independently regulated, the steady state linear regression slope of birth vs. division size is

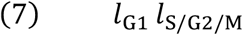

(SI). From Eq. 7, any mechanism that achieves apparent adder behavior between birth and division must exhibit a G1 behavior that compensates for the S/G2/M regulation, that is, *l*_G1_ = 1/*l*_S/G2/M_, which causes a negative correlation between birth size and G1 duration (Fig. 6B, SI). Failure to satisfy Eq. 7, signifying deviation from G1 compensatory behavior, implies that S/G2/M is not independently regulated, likely because progression through S/G2/M is limited by molecular species that persist through multiple cell-cycle phases. Otherwise, if the independently regulated S/G2/M phase is noisy, the contribution of G2/M checkpoint noise to cell size fluctuations may alter the apparent size-homeostasis behavior between birth and G1/S (*l*_G1_), because *η*_G2/M_ can affect linear regression slopes between birth and division (e.g. *η*_G2/M_ > *f*_phase_/*σ*_phase_ in Eqs. 1 and 4) without affecting the mode of independently regulated S/G2/M control (*f*_S/G2/M_ = *l*_S/G2/M_). By contrast, a stringently regulated independent phase (*η*_checkpoint_ ≪ *f*_phase_/*σ*_phase_) displays the same size-homeostasis behaviors between consecutive G2/Ms and consecutive G1/Ss at steady state (Fig. 6C, SI), regardless of other modeling assumptions. Then, Eq. 7 holds regardless of whether G1 or S/G2/M is the independently regulated phase, and any scenario that produces adder behavior over the cell cycle, between either birth and division or G1/Ss, produces the apparent G1 vs. S/G2/M compensatory behavior.

**Figure 6:**
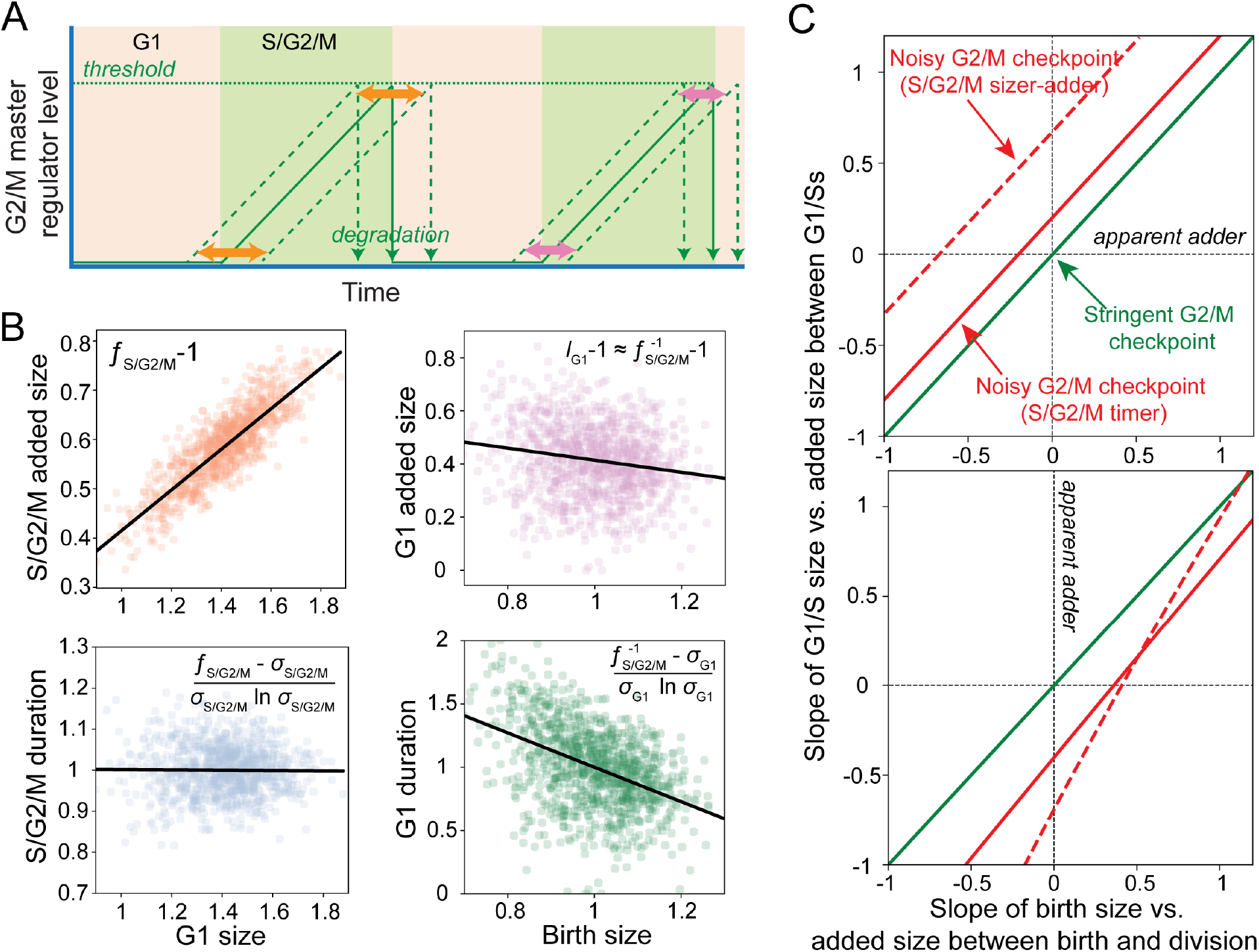
Apparent adder regulation between birth and division with non-adder regulation over G1 and S/G2/M can be achieved by independently regulated G1/S and G2/M transitions. (A)In independently regulated phases, size fluctuations (orange and pink) depend on events during but not prior to that phase. If G2/M is triggered by a cyclin-like master regulator that accumulates only through S/G2/M and is degraded outside of this phase, then S/G2/M is likely to be independently regulated from G1. (B) Simulations demonstrate a compensatory behavior in which the size added during G1 offsets the opposite trend over S/G2/M. This behavior is naturally achieved by control mechanisms that give apparent adder behavior between birth and division combined with an independently regulated phase. Panels: Simulated scatter plots of size-homeostasis statistics for a budding yeast-like G1/S inhibitor dilutor with low checkpoint noise (*η*_G1/S_,*η*_G2/M_ ≪ 1 in Eq. 4) and the corresponding analytical expressions for the linear regression slopes (SI). Noise terms: *ξ*_C,G1_, *ξ*_C,S/G2/M_ = 0.05, *ξ*_G1/S_, *ξ*_G2/M_ = 0.02 give realistic size fluctuations CV of G1/S size ≈ CV of G2/M size ≈0.1. (C) The apparent size-homeostasis behaviors between consecutive G2/Ms and consecutive G1/Ss are identical if the cell cycle is controlled by one independently regulated phase with low checkpoint noise (either G1/S or G2/M). Top: For a G1/S two-phase master regulator combined with an independently regulated S/G2/M phase and realistic noise levels (CV of G1/S size ≈ 0.1), the same size-homeostasis behavior is observed between birth and division and consecutive G1/Ss when the G2/M checkpoint is stringent (green line); the equality is lost as G2/M checkpoint noise contributions increases (red lines). Different size homeostasis behaviors were achieved by varying the size dependency of the regulator’s production rate (*λ_c_*). Bottom: results are similar for G1/S inhibitor dilutors combined with an independently regulated S/G2/M phase.

Size homeostasis measurements in the *A. thaliana* apical stem cell niche, an expanse of tissue at the plant apex that gives rise to all above-ground organs, established a linear regression slope of birth vs. division size ≈ 0.5 (Willis et al., 2016) (Fig. 7A). No mechanistic model has previously been proposed to explain this intermediate behavior between sizer and adder. CDK1-cyclin species are highly conserved as major G1/S and G2/M regulators throughout eukaryotes, including *A. thaliana* (Scofield et al., 2014). Whi5 has no structural *A. thaliana* homolog, but the *A. thaliana* human retinoblastoma (RBR1) homolog plays a functional role similar to Whi5 (Harashima and Sugimoto, 2016; Turner et al., 2012), raising the possibility that *A. thaliana* G1/S is regulated by an inhibitor dilutor.

**Figure 7:**
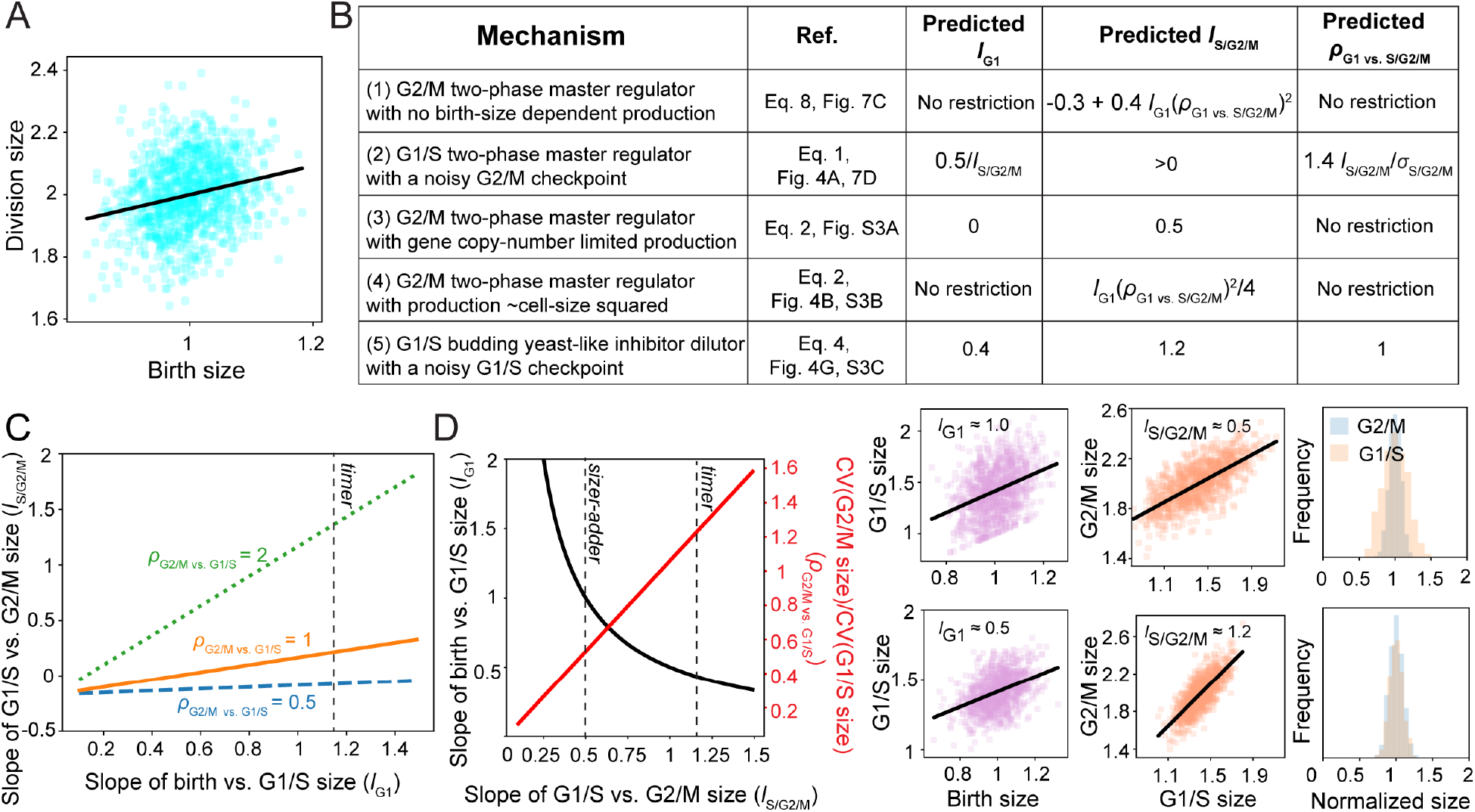
Extended size-homeostasis statistics can discriminate among different possible mechanisms underlying intermediate sizer-adder behavior. All simulations are based on realistic noise levels (*ρ*_G1/S_ ≈ (*ρ*_G2/M_ ≈ 0.1). Solid black lines in scatter plots correspond to linear regression fits. (A) A linear regression slope of ≈ 0.5 between birth and division sizes corresponds to the intermediate sizer-adder behavior identified in *A. thaliana* stem cells. (B) A summary of mechanisms (1)–(5) and the relevant equations and figures that generate intermediate sizer-adder behavior, in some cases with distinguishing predictions for the linear regression slopes between birth and G1/S sizes (*l*_G1_) and G1/S and G2/M sizes (*l*_S/G2/M_), and the ratio of the CVs in G2/M size vs. G1/S size (*ρ*_G2/M vs.G1/S_). (C) Mechanism 1: A G2/M two-phase master regulator produced in proportion to cell size with no birth-size or gene copy-number. Predictions for *l*_S/G2/M_ depend on *l*_G1_ and *ρ*_G2/M vs. G1/S_: there is apparent near-sizer behavior over S/G2/M (*l*_S/G2/M_ ≈ 0.0) if the CV in G1/S size exceeds or equals that of G2/M size (*ρ*_G2/M vs. G1/S_ ≤ 1). (D) Mechanism 2: A G1/S two-phase master regulator produced in proportion to growth with noisy supra-sizer S/G2/M regulation such that *η*_G2/M_ ≈ *f*_S/G2/M_/*σ*_S/G2/M_. Different modes of independently regulated S/G2/M control (*f*_S/G2/M_ = *l*_S/G2/M_) predict different *l*_G1_ s and *ρ*_G2/M vs. G1/S_ (left); for example, S/G/M intermediate sizer-adder vs. timer regulation (*l*_S/G2/M_ = 0.5 vs. *l*_S/G2/M_ = 1.2) predict *l*_G1_ ≈ 1.0 and *ρ*_G2/M vs. G1/S_ ≈ 0.6 (right, top row) vs. *l*_G1_ ≈ 0.5 and *ρ*_G2/M vs. G1/S_ ≈ 1.5 (right, bottom row), respectively. In the simulations, noise terms *ξ*_C,G1_, *ξ*_C,S/G2/M_, *ξ*_G1/S_ = 0.12; *ξ*_G2/M_ = 0.06 (top); *ξ*_C,G1_, *ξ*_C,S/G2/M_, *ξ*_G1/S_, *ξ*_G2/M_ = 0.06 (bottom) produce realistic CVs of cell size ≈ 0.1. Mechanisms (3–5) in panel (B) are represented in Fig. S3.

Since our model can generate homeostasis behaviors that vary continuously depending on factors such as regulator dynamics and noise levels, we applied Eqs. 1–7 to identify control mechanisms that could account for intermediate sizer-adder behavior. Here, we highlight five mechanisms taking into account that *A. thaliana* stem cells increase their average size by 1.5-fold over G1 (*σ*_G1_ = 1.5) (Dewitte et al., 2003) and by 2-fold over the cell cycle (*σ* = 2), implying a 1.3-fold increase over 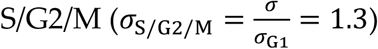, and grow exponentially throughout the cell cycle (*λ_g_* = 1) at a per unit size rate that correlates negatively with birth size (Willis et al., 2016). To account for the negative correlation, we model growth rate dependence on birth size fluctuations as *γ*(1 + *α_g_*Δ*S*_i,G1_) and regulator production rate dependence on initial size fluctuations as *k*_phase_(1 + *α*_*c*, phase_Δ*S*_i,phase_), where 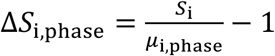 is the scaled deviation from the average size at the beginning of the phase (*μ*_i,phase_) (SI). In *A. thaliana* stem cells, *α_g_* ≈ −0.5 (Willis et al., 2016) while *α*_*c*,phase_ is unknown. This negative correlation is not unique to plants; it has recently been observed in mammalian and bacterial cell lines (Cadart et al., 2018; Nordholt et al., 2019). The hypothesized mechanisms are described below, with distinguishing predictions for other measurable statistics: the linear regression slopes between birth and G1/S sizes (*l*_G1_), between G1/S and G2/M sizes (*l*_S/G2/M_), and the relative CVs in G2/M size vs. G1/S size (*ρ*_G2/M vs.G1/S_) (Fig. 7B).

(1) A G2/M two-phase master regulator produced in proportion to cell size with no gene-copy number or birth-size dependence (*λ*_*c*,phase_ = 1, *r*_S/G2/M_ = 1, *α*_*c*,phase_ = 0) generates a linear regression slope of birth vs. division size of

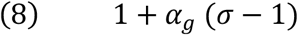

regardless of the mode of G1/S regulation and noise levels, producing intermediate sizer-adder behavior for *A. thaliana* stem cells where *α_g_* = –0.5, *σ* = 2. If *ρ*_G2/M vs.G1/S_ ≈ 1, as is the case for mammalian cells (Cadart, 2018), apparent near-sizer behavior between G1/S and G2/M (*l*_S/G2/M_ ≈ 0) is a key prediction (Fig. 7C, SI).

(2) A G1/S two-phase master regulator produced in proportion to growth with no gene-copy number effect on regulator production (*λ_c_* = 1, *r*_S/G2/M_ = 1 in Eq. 1, *α*_*c*,phase_ = –0.5 (SI)) with a noisy, independently regulated S/G2/M such that *η*_G2/M_ ≈ *f*_S/G2/M_/σ_S/G2/M_. Key predictions are supra-sizer S/G2/M control *f*_S/G2/M_ = *l*_S/G2/M_ > 0, *l*_G1_ ≈ 0.5/*l*_S/G2/M_ and 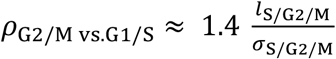 (Fig. 7D).

(3) A G2/M two-phase master regulator produced with the same size dependence as growth (*λ_c_* = 1 in Eq. 2, *α*_*c*,phase_ = −0.5, SI), but, unlike growth, the regulator’s production rate is gene copy-number limited and doubles during S/G2/M (*r*_S/G2/M_ = 2). Key predictions are *l*_G1_ ≈ 0 and *l*_S/G2/M_ ≈ 0.5 (Fig. S3A).

(4) A G2/M two-phase master regulator produced in proportion to the square of cell size throughout the cell cycle (*λ_c_* = 2 in Eq. 2) with the same birth-size dependence as growth and no gene copy-number effect (*α*_*c*,phase_ = −0.5, *r*_S/G2/M_ = 1). Predictions are similar to those of mechanism (1) (Fig. 7C vs. S3B), so it is necessary to measure regulator dynamics to distinguish between mechanisms (1) and (4).

(5) A budding yeast-like G1/S inhibitor dilutor produced at a constant birth-size independent rate through S/G2/M while S/G2/M exhibits timer regulation and a noisy G1/S threshold concentration (*η*_G1/S_ ≈ 0.7 in Eq. 4). Predictions are *l*_G1_ ≈ 0.4, *l*_S/G2/M_ ≈ 1.2, and *ρ*_G2/M vs.G1/S_ ≈ 1 (Fig. S3C).

## Discussion

Here, we developed a general, mechanistic model of cell proliferation with two cell cycle phases, aiming to achieve a pragmatic tradeoff between the representation of cell cycle complexity and model analyzability (Fig. 1). We applied the model to determine how size homeostasis is broken without necessarily disrupting the proper ordering of G1/S and G2/M or division by changing mechanisms of cell-cycle regulation (Fig. 2,3). For example, size homeostasis would break: (1) if the width of FtsZ bands were to increase proportionally with growth to trigger division at a local threshold density; (2) in slow-growing bacteria without multiple replication forks, if DnaA activity were to scale independently of or strongly with cell size while S/G2/M were under timer control; (3) in budding yeast, if Whi5 was produced in proportion to cell size rather than at a constant rate; (4) in many cases, with gene-copy number effects on regulator production. These findings reveal unintuitive constraints on cell-cycle regulation imposed by size homeostasis requirements. They explain why certain patterns of cell cycle regulation have been observed and not others, and suggest a breadth of cellular designs for the loss of size homeostasis, potentially enabling experimentalists to probe the physiological implications of a transient loss of size control.

General analytical expressions connect cell cycle control mechanisms to measured size homeostasis statistics (Eqs. 1-8, Methods, SI), thus providing a linchpin that ultimately connects genotype to size-homeostasis phenotype. In some cases, unintuitive implications were revealed, such as the potential enhancement of adder behavior by noisy regulator production (Fig. 4), potential deviations from adder by regulator localization patterns (Fig. 5), and the curious statistical signatures of independently regulated phases (Fig. 6). We have inevitably approximated or omitted certain details of cell proliferation, yet the analytical expressions are a powerful basis for generating and testing hypotheses across a range of scenarios. To exemplify this power, we enumerated five mechanisms that account for the intermediate sizer-adder behavior between birth and division observed in *A. thaliana* apical stem cells with distinguishing predictions for other size-homeostasis statistics (Fig. 7). One plausible mechanism assumes CDK1-cyclin behaves as a master regulator triggering G2/M at a threshold level and is produced proportionally with cell size rather than growth rate throughout the cell cycle, thus implying that regulator production scales with bulk synthetic capacity of the cell (which presumably scales with size) rather than being directly coupled to growth. Then, if the CVs in cell size at G1/S and G2/M are similar, apparent near-sizer behavior over S/G2/M, corresponding to a zero correlation between G1/S size and G2/M size, is predicted (Fig. 7C). These predictions can be readily tested by quantitative time-lapse imaging of *A. thaliana* apical stem cells in strains containing extant G1/S and membrane reporters (Jones et al., 2017; Willis et al., 2016).

Why is adder behavior ubiquitous? Adder regulation may not be evolutionary advantageous *per se*, but may tend to result from evolutionary pressures on underlying regulator and growth dynamics with the necessity that catastrophic loss of size control is avoided. In fission yeast, widely conserved CDK-cyclin-like master regulators trigger multiple ordered events at successive threshold levels, operating as an “arrow of time” for the cell cycle (Swaffer et al., 2016). This ordering activity may generate pressures to degrade the regulator, thus preventing events in previous cell cycles from affecting future events, and to match the regulator production rate to the biosynthetic capacity of the cell to ensure events occur only when cellular machineries are sufficiently plentiful. Then, events are necessarily triggered at a threshold level rather than concentration to avoid a loss of size control and, for regulators of DNA replication initiation which persist between intervening divisions such as active DnaA, size-homeostasis robustness precludes a production rate that deviates strongly from being proportional to growth. These mechanisms result in apparent adder behavior between birth and division without implementing a “molecular adder” that specifically effects a fixed added-size increment. The use of cell size statistics alone to conclude “molecular adders” may lead to incorrect inferences about the basis of cell-cycle regulation, recalling flawed conclusions about intrinsic noise derived from a dual reporter system (Hilfinger and Paulsson, 2011). In slow growing cells where the constraints on the timing of events may be relatively relaxed, deviations from these mechanisms and thus from adder behavior may arise. Alternatively, adder regulation of S/G2/M may be advantageous in conferring a robustness to size control against variations in other cell proliferation factors (Fig. 2A, 3A-C), especially the dynamics of G1/S regulation; in many conditions and organisms, G1/S occurs shortly after G2/M or division, so the observed adder regulation between birth and division may be generated by S/G2/M adder regulation.

In general, our findings exemplify how the model combined with quantitative time-lapse measurements of cell size dynamics and cell cycle reporters across species, mutants, and conditions should not only help to establish the mechanisms of cell cycle regulation, but further illuminate their necessity for size control. Such a fundamental understanding will inform whether the frequently observed adder behavior emerges from a common mechanism, is a product of convergent evolution due to selective pressures on size homeostasis, or is merely a manifestation of specific regimes of mechanisms without molecular adders that happened to have been the focus of experiments thus far. The intimate connections between size control and other cellular processes should also be an important factor in probing the response of cells to non-steady-state conditions and to the future design of artificial cells.

## Acknowledgements

The authors thank members of the Huang Laboratory and Clotilde Cadart, Po-Yi Ho, Benjamin Knapp, and Fred Chang for helpful discussions. This work was supported by the Allen Discovery Center at Stanford on Systems Modeling of Infection (to K.C.H.) and the Gatsby Charitable Foundation under Grant GAT3395/PR4 (to H.J.). K.C.H. is a Chan Zuckerberg Biohub Investigator. This work was also supported in part by the National Science Foundation under Grant PHYS-1066293 (to K.C.H.) and the hospitality of the Aspen Center for Physics.

## Author Contributions

Conceptualization, L.W., H.J., and K.C.H.; Methodology, L.W. and K.C.H.; Formal analysis, L.W.; Writing-Original draft, L.W. and K.C.H.; Writing-Review and editing, L.W., H.J., and K.C.H.; Funding acquisition, L.W., H.J., and K.C.H.

## Supplemental Figures

**Figure S1:**
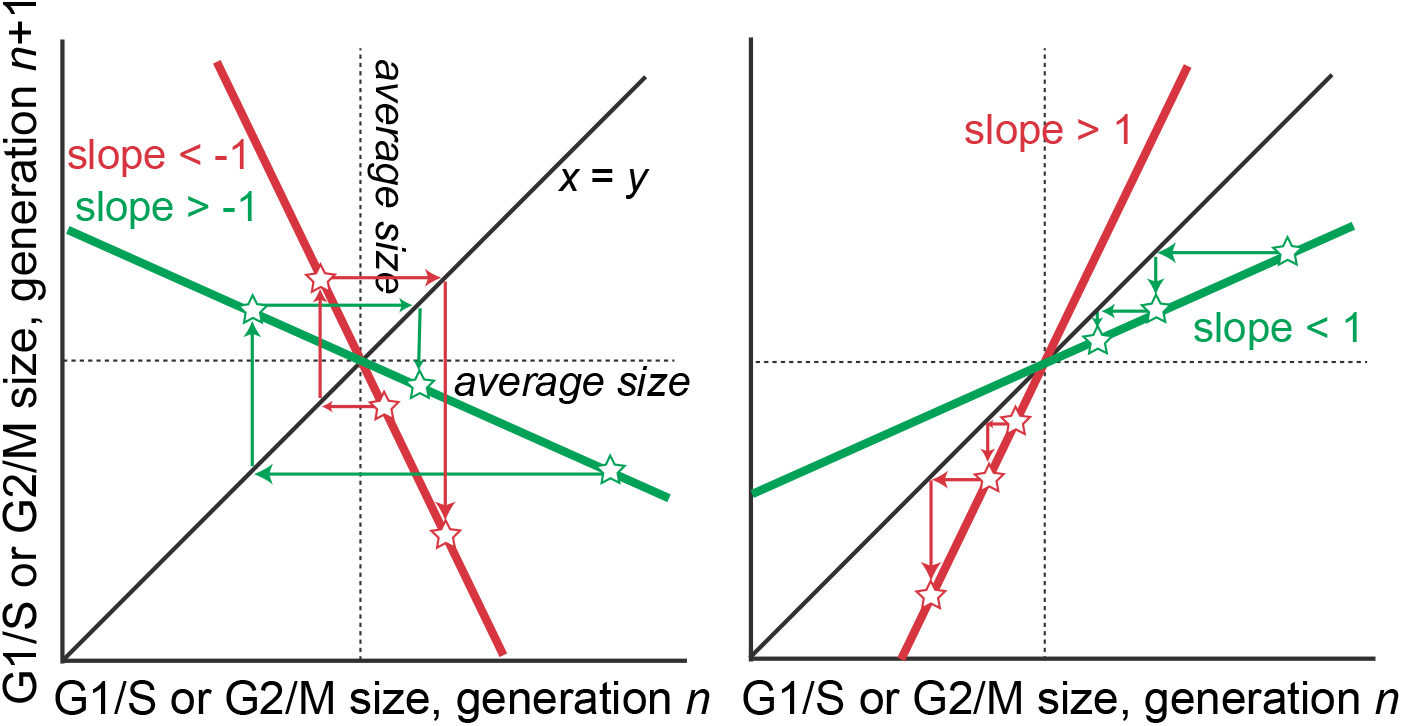
Conditions for loss of cell-size homeostasis. Related to Figures 2 and 3. For cells to achieve control of G1/S or G2/M size, the linearized relationship between cell sizes at G1/S or G2/M in consecutive cell cycles must have a slope between −1 and 1 (green trajectories). Size fluctuations then regress to the average, as opposed to diverging (red trajectories).

**Figure S2:**
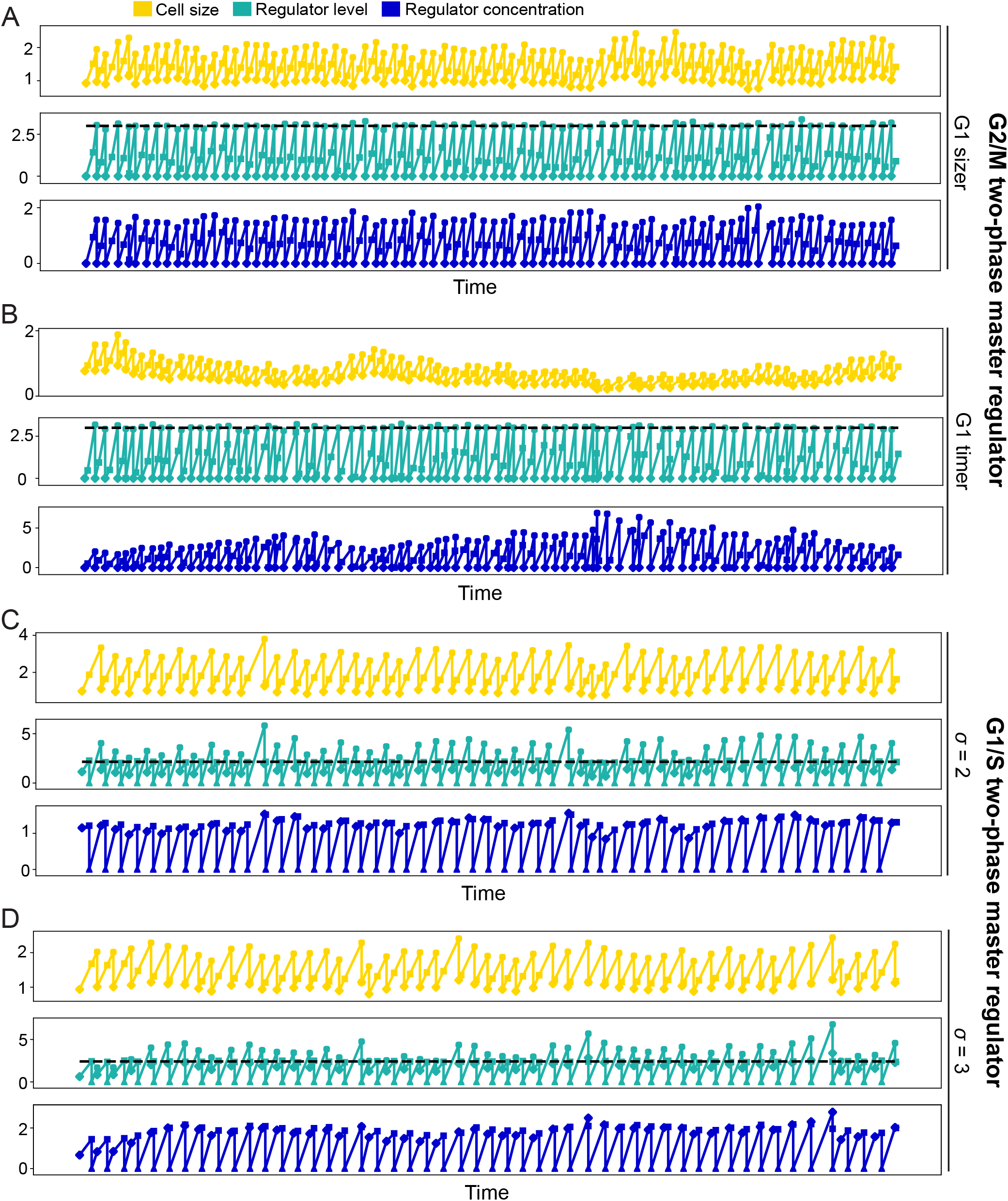
Simulations of single-cell lineage trajectories support analytical findings. Related to Figure 3. G2/M and G1/S master regulators produced through both phases (two-phase master regulators) fail to implement size homeostasis under certain conditions. (A,B) For exponentially growing cells (*λ_g_* = 1), when regulator production is gene copy-number limited (*r*_S/G2/M_ = 2), G2/M two-phase master regulators produced at a size-independent rate (*λ*_*c*,G1/S_ = *λ*_*c*,S/G2/M_ = 0) achieve size homeostasis for G1 critical size or adder control regardless of the average G1 duration (*τ*), but fail to achieve size homeostasis for G1 timer control (Fig. 3A, Table S2). (A) shows size and regulator level homeostasis for G1 critical size regulation (*f*_G1_ = 0, *τ* = 0.5, *σ* = 2; noise terms *ξ*_C,G1_, *ξ*_C,S/G2/M_ = 0.05, *ξ*_G1/S_, *ξ*_G2/M_ = 0.1 produce a CV of cell size ≈ 0.1); (B) shows the loss of homeostasis when G1 control approaches timer (*f*_G1_ = 0.99 *σ^τ^*) while other parameters remain fixed. Diamonds, squares, and discs denote values at birth, G1/S, and G2/M or division, respectively. (C,D) G1/S two-phase master regulators that are gene copy-number limited fail to implement G1/S size homeostasis when combined with G2/M sizer regulation (*f*_S/G2/M_ = 0) for binary fission or asymmetric division with *σ* ≤ 2, regardless of G1 duration, the size dependence of regulator production, and growth pattern (Fig. 3B, Table S2); (C) shows size homeostasis for gene copy-number limited (*r*_S/G2/M_ = 2) G1/S two-phase master regulators when *σ* = 3 (*τ* = 0.5, linear growth pattern *λ_g_* = 0; regulator production *λ*_*c*,G1/S_ = *λ*_*c*,S/G2/M_ = 0; and noise terms *ξ*_C,G1_, *ξ*_C,S/G2/M_, *ξ*_G1/S_ = 0.02, *ξ*_G2/M_ = 0.1 producing a CV of cell size ≈ 0.1); (D) shows the loss of G1/S size homeostasis when *σ* = 2 while other parameters remain fixed. Diamonds, squares, triangles, and discs denote values at birth, G1/S, immediately after G1/S when the regulator is degraded, and G2/M or division, respectively. Horizontal dashed lines show the threshold regulator level triggering phase progression.

**Figure S3:**
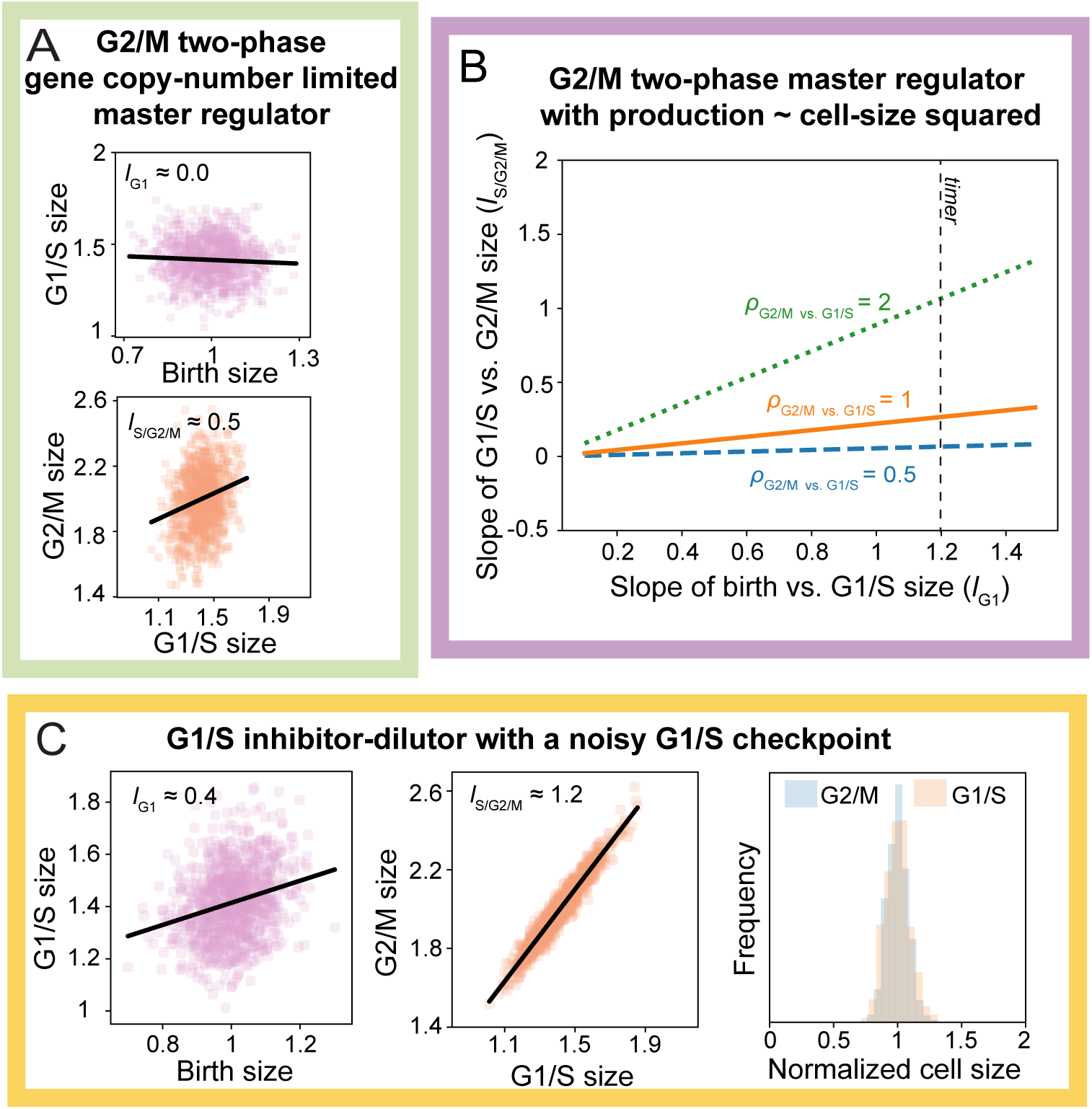
Extended size-homeostasis statistics can discriminate among different possible mechanisms underlying intermediate sizer-adder behavior. Related to Figure 7; the panels represent mechanisms (3-5) listed in Fig. 7B. In all simulations, noise terms were set to produce realistic CVs of cell size ≈ 0.1. (A)Mechanism 3: A G2/M two-phase master regulator produced in proportion to cell size (*λ*_*c*,phase_ = 1) with the same birth-size dependence as growth (*α*_*c*,phase_ = −0.5) (SI) and gene copy-number limited production (*r*_S/G2/M_ = 2), with the latter implying that regulator production is not proportional to growth throughout the cell cycle. Predictions are G1/S sizer regulation (*l*_G1_ ≈ 0; top) and apparent sizer-adder regulation over S/G2/M (*l*_S/G2/M_ ≈ 0.5; bottom). In the simulation, noise terms were *ξ*_C,G1_, *ξ*_C,S/G2/M_, *ξ*_G1/S_, *ξ*_G2/M_ = 0.08. (B) Mechanism 4: A G2/M two-phase master regulator produced in proportion to cell-size squared (*λ*_*c*,G1_ = *λ*_*c*,S/G2/M_ = 2) with no gene copy-number effect (*r*_S/G2/M_ = 1) and the same birth-size dependence as growth (*α*_*c*,G1_ = *α*_*c*,S/G2/M_ = −0.5) (SI). Predictions are similar to mechanism (1) (Fig. 7C). (C) Mechanism 5: A G1/S budding yeast-like inhibitor-dilutor with high G1/S checkpoint noise. Further predictions are intermediate sizer-adder behavior over G1 (*l*_G1_ ≈ 0.4), apparent near adder behavior over S/G2/M (*l*_S/G2/M_ ≈ 1.2), and similar coefficients of variation in G1/S and G2/M sizes (*ρ*_G2/M vs. G1/S_ ≈ 1). In the simulation, noise terms were *ξ*_C,G1_, *ξ*_C,S/G2/M_, *ξ*_G2/M_ = 0.02; *ξ*_G1/S_ = 0.08.

**Table S1.** Average birth size (*μ*_i,G1_) is determined by model parameters. Left: Average birth size for a G2/M two-phase master regulator where growth and regulator production have the same size dependences in both phases *λ_g_* = *λ*_*c*,G1_ = *λ*_*c*,S/G2/M_; Fig. 1A,C), Θ is the threshold level for G2/M, and Θ_deg_ is the fixed level that the regulator is degraded to following G2/M. Middle: Average birth size for a G1/S two-phase master regulator where growth and regulator production have the same size dependencies in both phases (*λ_g_* = *λ*_*c*,G1_ = *λ*_*c*,S/G2/M_). Right: Average birth size for a G1/S inhibitor dilutor that triggers G1/S at a minimum concentration *k*_thres_ with a growth size dependence that exceeds the regulator production’s size dependence as in budding yeast (*λ_g_* – *λ*_*c*,S/G2/M_ = 1). See SI for similar expressions in more general cases.

| Average birth size<br>G2/M two-phase master regulator | Average birth size<br>G1/S two-phase master regulator | Average birth size<br>G1/S inhibitor dilutor |
| --- | --- | --- |
| $\frac{\Theta - \Theta_{deg}}{\frac{\kappa_{G1}}{\gamma}(\sigma^\tau - 1) + \frac{\kappa_{S/G2/M}}{\gamma}(\sigma - \sigma^\tau)}$ | $\frac{\Theta - \Theta_{deg}/\sigma}{\frac{\kappa_{S/G2/M}}{\gamma}(1 - \sigma^{\tau-1}) + \frac{\kappa_{G1}}{\gamma}(\sigma^\tau - 1)}$ | $\frac{\ln \sigma (1 - \tau)}{\sigma^\tau(\sigma - 1)} \frac{\kappa_{S/G2/M}}{\gamma k_{thres}}$ |

**Table S2.** Expressions for the slopes between first-order cell-size fluctuations away from the mean at consecutive G2/Ms and consecutive G1/Ss in the absence of noise (Methods, SI). When the absolute values of these slopes exceed 1, as shown in the colored regions of Figs. 2A and 3A-C, size homeostasis is lost (Methods, Fig. S1). In these figures, cell cycles are controlled by a two-phase master regulator or an inhibitor dilutor of G1/S or G2/M combined with an independently regulated phase, for example, timer, adder, or sizer regulation corresponding to *f*_phase_ = *σ*_phase_, 1, or 0, respectively. Here, we assumed: for two-phase master regulators, production and growth have the same size-dependencies through both phases (Δ*λ* = *λ_g_* – *λ*_*c*,G1_ = *λ_g_* – *λ*_*c*,S/G2/M_); growth is exponential, so *λ_g_* = 1 and *σ*_G1_ = *σ^τ^* (Methods, Fig. 1F). For two-phase master regulators, expressions are for no effect of gene copy number on regulator production (*r*_S/G2/M_ = 1, Fig. 1A), or a strong limiting effect of gene copy number (*r*_S/G2/M_ = 2). The fully general expressions are in SI.

**Linear slopes between consecutive G2/M sizes**
| S/G2/M<br>sizer | S/G2/M adder | S/G2/M timer<br>(exponential growth) |
| --- | --- | --- |
| 0 | As for G1/S | As for G1/S |

**Linear slopes between consecutive G1/S sizes**
| S/G2/M sizer | S/G2/M adder | S/G2/M timer<br>(exponential growth) |
| --- | --- | --- |
| $\frac{1 - \frac{(\sigma - 1)(1 - \Delta\lambda)}{\sigma^{(1-\tau)(1-\Delta\lambda)} - 1}}{\sigma}$ | $\frac{1 + (\sigma^{-(1-\tau)\Delta\lambda} - 1) \frac{(\sigma - 1)(1 - \Delta\lambda)}{\sigma^{(1-\tau)(1-\Delta\lambda)} - 1}}{\sigma}$ | $\frac{\sigma - \Delta\lambda(\sigma - 1)}{\sigma}$ |

**Linear slopes between consecutive G2/M sizes**
|  | G1 sizer | G1 adder | G1 timer (exponential growth) |
| --- | --- | --- | --- |
| $r_{S/G2/M} = 2$ | $\frac{1}{2\sigma^{1-\Delta\lambda}}$ | $\frac{1 + \sigma^{-\tau\Delta\lambda}}{2\sigma^{1-\Delta\lambda}}$ | $\frac{1 + \sigma^{\tau(1-\Delta\lambda)}}{2\sigma^{1-\Delta\lambda}}$ |
| $r_{S/G2/M} = 1$ | $\frac{1}{\sigma^{1-\Delta\lambda}}$ | $\frac{1}{\sigma^{1-\Delta\lambda}}$ | $\frac{1}{\sigma^{1-\Delta\lambda}}$ |

**Linear slopes between consecutive G1/S sizes**
|  | G1 sizer | G1 adder | G1 timer (exponential growth) |
| --- | --- | --- | --- |
| $r_{S/G2/M} = 2$ | 0 | As for G2/M | As for G2/M |
| $r_{S/G2/M} = 1$ | 0 | As for G2/M | As for G2/M |

**Linear slopes between consecutive G2/M sizes**
|  | S/G2/M<br>sizer | S/G2/M adder | S/G2/M timer<br>(exponential growth) |
| --- | --- | --- | --- |
| $r_{S/G2/M} = 2$ | 0 | As for G1/S | As for G1/S |
| $r_{S/G2/M} = 1$ | 0 | As for G1/S | As for G1/S |

**Linear slopes between consecutive G1/S sizes**
|  | S/G2/M<br>sizer | S/G2/M adder | S/G2/M timer<br>(exponential growth) |
| --- | --- | --- | --- |
| $r_{S/G2/M} = 2$ | $\frac{2}{\sigma}$ | $\frac{2 - \sigma^{\tau\Delta\lambda}(2\sigma^{-\Delta\lambda} - 1)}{\sigma}$ | $\frac{2 - \sigma^{\tau(\Delta\lambda-1)+1}(2\sigma^{-\Delta\lambda} - 1)}{\sigma}$ |
| $r_{S/G2/M} = 1$ | $\frac{1}{\sigma}$ | $\frac{1 - \sigma^{\tau\Delta\lambda}(\sigma^{-\Delta\lambda} - 1)}{\sigma}$ | $\frac{1 - \sigma^{\tau(\Delta\lambda-1)+1}(\sigma^{-\Delta\lambda} - 1)}{\sigma}$ |

## Methods

Below, the model description of average cellular behaviors included in the Results is extended, then noise, simulations, and analytical derivations are explained.

### Models

We study two classes of regulators, with total intracellular level *C*, that trigger G1/S or G2/M progression and are produced in a potentially size-dependent manner with negligible degradation. Parameters *λ*_*c*,phase_ in phases G1 and S/G2/M determine the cell size (*S*) dependencies of regulator production 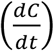 according to

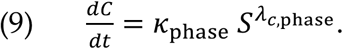

Master regulators are produced through one or two phases to trigger G1/S or G2/M progression upon reaching a total intracellular threshold level; degradation follows to a fixed level (e.g. zero; analyses show the degraded level has no impact on size homeostasis behaviors, see SI) (Fig. 1A). Inhibitor dilutors are produced throughout one phase and then diluted out in the subsequent phase to trigger G1/S or G2/M at a minimum threshold concentration (Fig. 1A). G2/M and division are assumed to be coincident. Upon division, cells divide symmetrically (*σ* = 2) or asymmetrically (*σ* ≠ 2) in a ratio 1:(*σ* − 1) and any regulator that persists is inherited in proportion to daughter cell size (Fig. 1B). Hence, at steady state population dynamics, the overall fold-size increase is *σ*, and division-plane positioning is independent of the preceding birth and G1/S sizes. Analyses and simulations assume steady state population dynamics of one cell type: following each division, only one daughter cell, corresponding to an average portion size of 1 and not *σ* − 1, is retained for analyses or simulations. The cellular growth rate 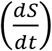 depends on cell size according to

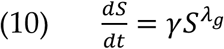

where *λ_g_* determines the growth type (exponential for *λ_g_* = 1 and linear for *λ_g_* = 0) and *γ* sets the average timescale for growth (Fig. 1C).

Master regulators or inhibitor dilutors are often considered in combination with an independently regulated S/G2/M or G1 phase: cell size at the end of the phase (*S*_e,phase_) depends on cell size at the beginning of the phase (*S*_i,phase_) according to

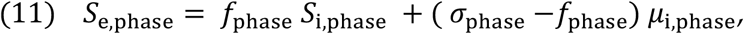

where *f*_phase_ is the mode of control (*f*_phase_ = 0, 1, or depends on growth behavior for “sizer”, “adder”, or “timer” control, respectively), *σ*_phase_ >1 is the average fold-size increase, and μ_O_,*_phase_ is the average initial size at steady state (Fig. 1D). The steady state fold-size increase over G1 and S/G2/M are related to the fraction of the cell cycle spent in G1 (*τ*) by *σ*_G1_ ≈ *σ^τ^* and *σ*_S/G2/M_ ≈ *σ*^1−*τ*^ because *σ*_S/G2/M_ = *σ*/*σ*_G1_. (The approximations are exact for exponential growth where the cell cycle duration is ln *σ/γ*, so 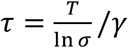 if *T* is the average G1 duration and 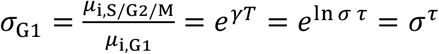.) A convenient aspect of the model is that the natural choice for free parameters changes as the control type of the independently regulated phase changes. For example, for an independently regulated phase under timer control, the natural free parameter is the duration of the phase or the fraction of the cell cycle spent in the phase (*τ*), whereas for critical size or equivalently sizer control, the natural choice is the average cell size at the transition (*μ*_e,phase_). Regardless of the natural choice, at steady state an equation connecting *τ* and *μ*_i,phase_ or *μ*_e,phase_ (SI) allows us to work in terms of the parameter *τ* (or *σ*_G1_). Then parameter sets that fail to implement cell cycles with two checkpoints on average are straightforward to exclude by enforcing 0 ≤ *τ* ≤ 1 (Fig. 2A, 3A-C).

More general analyses in SI allow growth and production rates to continually depend on cell size at the beginning of the phase, growth parameters *λ_g_* and *γ* to differ in G1 vs. S/G2/M, and the threshold mechanisms for cell cycle checkpoint progression to vary.

### Simulations and noise

Cells were initialized to the steady-state birth size and regulator level plus noise (SI). Then, cell sizes and regulator levels at G1/S and G2/M were simulated according to specified average cell-cycle control and growth parameters (Eq. 9–11) with 4 noise terms in cell-cycle regulation. We present a specific example; the general case is described in detail in SI. If an inhibitor dilutor triggers G1/S at a threshold concentration (*k*_thres_), then at the end of the G1 dilution phase the regulator level (*C*_G1/S_) and cell size (*S*_G1/S_) depend on the regulator level at birth (*C_b_*) as

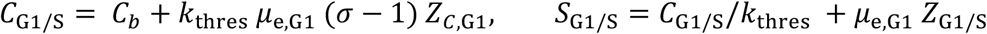

where *Z*_*C*,G1_ and *Z*_G1/S_ are zero-mean Gaussian noise terms that perturb the level of inhibitor over the G1 dilution phase and the G1/S threshold concentration, respectively. The noise terms’ coefficients give convenient interpretations for the corresponding standard deviations (s.d.). For example, *ξ*_G1/S_, the s.d. of *Z*_G1/S_ (equal to the typical G1/S error, *e*_G1/S_, in Results), approximates the coefficient of variation (CV) of the threshold concentration (SI). If S/G2/M is independently regulated, at G2/M, the cell size (*S*_G2/M_) and regulator level (*C*_G2/M_) are

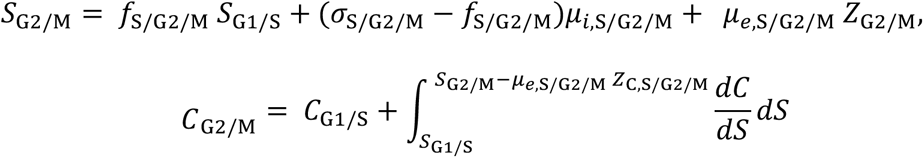

where *Z*_G2/M_ is a zero-mean Gaussian noise term that perturbs the G2/M checkpoint, ∫(*dC*/*dS*) *dS* is found explicitly from Eq. 9 divided by Eq. 10, and *Z*_*C*,S/G2/M_ is a zero-mean Gaussian noise term in inhibitor production compared with growth through S/G2/M. The interpretation of *ξ*_G2/M_, the s.d. of *Z*_G2/M_, depends on the mode of S/G2/M control (SI); for example, if the control is sizer (*f*_S/G2/M_ = 0), *ξ*_G2/M_ is the CV in the G2/M critical cell size. The regulator level and cell size at birth of the retained daughter in the next generation are

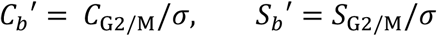

where no noise in division was assumed. Analytical derivations (SI) show that size homeostasis behaviors depend on just two noise-related parameters, *η*_G1/S_ = *ξ*_G1/S_/CV(cell size at G1/S) and *η*_G2/M_ = *ξ*_G2/M_/CV(cell size at G1/S) (Fig. 1E). In general, the noise terms that cause deviation from the average coupling between inhibitor dynamics and growth, *Z*_*C*,G1_ and *Z*_*C*,S/G2/M_, contribute to cell size at G1/S without affecting *ξ*_G1/S_ and *ξ*_G2/M_ and thus reduce *η*_G1/S_ and *η*_G2/M_.

In simulations, growth is forced to be non-negative over each phase, so *S*_b_ < *S*_G1/S_ < *S*_G2/M_, and regulator levels are forced to be non-negative (SI). This forcing is not included in analytical derivations, yet the derivations are in excellent agreement with simulations, indicating that it has minimal impact on results. Cells were simulated for 30+ generations until steady states were patently reached.

### Analysis

Throughout analyses, linear regression slopes between cell-size variables (Eqs. 1–8) were derived as follows: scaled cell-size fluctuations at each transition (Δ*S*_G1/S_ = *S*_G1/S_/*μ*_*e*,G1_ − 1 and Δ*S*_G2/M_ = *S*_G2/M_/*μ*_*e*,S/G2/M_ −1) were expressed in terms of scaled size fluctuations at earlier transitions, then only linear terms from a Taylor expansion and noise terms were retained for analyses, because cell size fluctuations are small (in most measurements, the coefficient of variation in cell size is ~0.13 (Cadart, 2018; Cadart et al., 2018; Taheri-Araghi et al., 2015; Willis et al., 2016)) and noise terms are comparable in magnitude to cell size fluctuations (Amir, 2014). Indeed, analytically derived linear regression slopes are in excellent agreement with simulations with realistic noise levels, indicating that the linear approximation is appropriate (Fig. 2–4).

We present two examples; the general case is detailed in SI. For a one-phase master regulator produced from a fixed level at G1/S through S/G2/M to trigger G2/M when it is degraded, the linearized relationship between Δ*S*_G1/S_ and Δ*S*_G2/M_ is found by solving Eqs. 9 and 10 followed by a Taylor expansion,

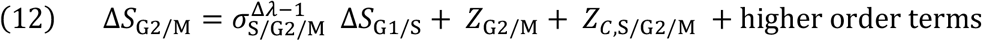

where Δ*λ* = *λ_g_* − *λ*_*c*,S/G2/M_. By definition, the linear regression slope between G1/S and G2/M sizes (*l*_S/G2/M_) is

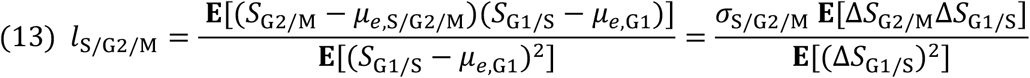

where **E**[·] denotes the average (SI). So, the slope *l*_S/G2/M_ is computed by multiplying Eq. 12 by *σ*_S/G2/M_Δ*S*_G1/S_, taking averages of each side of the equation, and dividing by **E**[(Δ*S*_G1/S_)^2^]. Since G1/S size fluctuations (Δ*S*_G1/S_) are independent of noise in the subsequent G2/M threshold (*Z*_G2/M_) and regulator dynamics compared with growth over S/G2/M (*Z*_*C*,S/G2/M_), upon taking averages, the noise terms disappear. Thus, the linear regression slope between G1/S and G2/M sizes is

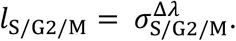

From the definition of independently regulated phases (Eq. 11) with noise, after rearrangement, we have

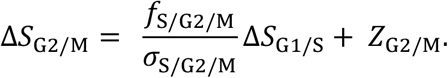

Again, the linear regression slope between G1/S and G2/M sizes is computed according to Eq. 13, to give

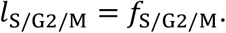

Eqs. 1–8 were derived similarly but often the dependence of size fluctuations on noise terms in preceding phases causes noise to affect size homeostasis behaviors (SI). Importantly, throughout the analytical derivations, no assumptions were made about the distributions of *Z*_G1/S_, *Z*_C,G1_, *Z*_G2/M_, and *Z*_C,S/G2/M_ beyond the values of their standard deviations, indicating that, for small fluctuations in cell size, further properties of the distributions (e.g. skewness) have no effect on size homeostasis behaviors.

### Conditions for the loss of size homeostasis

Derivations of the type above led to first-order expressions connecting size fluctuations at G1/S and G2/M in consecutive cell cycles

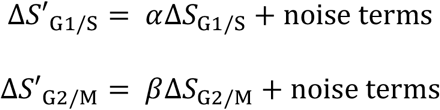

where denotes the subsequent cell cycle, and *α* and *β* are functions of parameters (SI). These equations establish whether fluctuations from the average cell size diverge, so that size homeostasis is lost, even in the absence of noise according to whether |*α*| ≥ 1 or |*β*| ≥ 1 for G1/S or G2/M, respectively (Fig. S1). The colored regions of Fig. 2A and 3A-C show where |*α*| ≥ 1 or |*β*| ≥ 1 for the parameters specified in each plot.

Simulations of single-cell trajectories with |*α*| ≈ 1 and |*β*| ≈ 1, i.e. close to the boundary of size homeostasis, show that size homeostasis is compromised (Fig. 2B, 3D, S2), thus supporting our analyses.

### General expressions for size-homeostasis statistics

The full model features up to 22 parameters (SI). This space is too large to explore computationally. Analytical derivations effectively shrunk the parameter space, because only certain parameter combinations affect size homeostasis statistics. Table S2 and Eqs. 1–8 exemplify specific cases where size homeostasis statistics depend on a parameter set that is effectively strongly reduced, with fully general expressions derived in SI.

Regardless of the nature of cell cycle control, the linear regression slope between scaled fluctuations in the duration of the phase and scaled size fluctuations at the beginning of the phase is

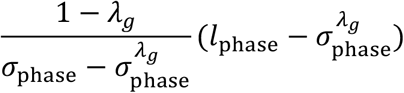

where *l*_phase_ is the linear regression slope between cell size at the beginning vs. cell size at the end of the phase (SI). This slope must be zero for independently regulated timer phases, thus, 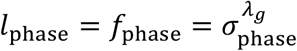.

## References

Amir, A. (2014). Cell size regulation in bacteria. Phys Rev Lett 112, 208102.

Barber, F., Ho, P.Y., Murray, A.W., and Amir, A. (2017). Details matter: noise and model structure set the relationship between cell size and cell cycle timing. Front Cell Dev Biol 5.

Cadart, C. (2018). Personal communication.

Cadart, C., Monnier, S., Grilli, J., Attia, R., Terriac, E., Baum, B., Cosentino-Lagomarsino, M., and Piel, M. (2018). Size control in mammalian cells involves modulation of both growth rate and cell cycle duration. Nat Commun 9, 3275.

Campos, M., Surovtsev, I.V., Kato, S., Paintdakhi, A., Beltran, B., Ebmeier, S.E., and Jacobs-Wagner, C. (2014). A constant size extension drives bacterial cell size homeostasis. Cell 159, 1433–1446.

Chandler-Brown, D., Schmoller, K.M., Winetraub, Y., and Skotheim, J.M. (2017). The adder phenomenon emerges from independent control of pre-and post-start phases of the budding yeast cell cycle. Curr Biol 27, 2774–2783.

Coudreuse, D., and Nurse, P. (2010). Driving the cell cycle with a minimal CDK control network. Nature 468, 1074–1079.

Cross, F.R. (1988). DAF1, a mutant gene affecting size control, pheromone arrest, and cell cycle kinetics of *Saccharomyces cerevisiae*. Mol Cell Biol 8, 4675–4684.

Dewitte, W., Riou-Khamlichi, C., Scofield, S., Healy, J.M., Jacqmard, A., Kilby, N.J., and Murray, J.A. (2003). Altered cell cycle distribution, hyperplasia, and inhibited differentiation in Arabidopsis caused by the D-type cyclin CYCD3. Plant Cell 15, 79–92.

Di Talia, S., Skotheim, J.M., Bean, J.M., Siggia, E.D., and Cross, F.R. (2007). The effects of molecular noise and size control on variability in the budding yeast cell cycle. Nature 448, 947–951.

Eun, Y.-J., Ho, P.-Y., Kim, M., LaRussa, S., Robert, L., Renner, L.D., Schmid, A., Garner, E., and Amir, A. (2018). Archaeal cells share common size control with bacteria despite noisier growth and division. Nat Microbiol 3, 148–154.

Facchetti, G., Knapp, B., Flor-Parra, I., Chang, F., and Howard, M. (2019). Reprogramming cdr2-dependent geometry-based cell size control in fission yeast. Curr Biol 29, 350–358.

Fantes, P.A. (1977). Control of cell size and cycle time in *Schizosaccharomyces pombe*. J Cell Sci 24, 51–67.

Ginzberg, M.B., Chang, N., D’Souza, H., Patel, N., Kafri, R., and Kirschner, M.W. (2018). Cell size sensing in animal cells coordinates anabolic growth rates and cell cycle progression to maintain cell size uniformity. ELife 11, e26957.

Harashima, H., Dissmeyer, N., and Schnittger, A. (2013). Cell cycle control across the eukaryotic kingdom. Trends in Cell Biology 23, 345–356.

Harashima, H., and Sugimoto, K. (2016). Integration of developmental and environmental signals into cell proliferation and differentiation through RETINOBLASTOMA-RELATED 1. Curr Opin Plant Biol 29, 95–103.

Heldt, F.S., Lunstone, R., Tyson, J.J., and Novák, B. (2018). Dilution and titration of cell-cycle regulators may control cell size in budding yeast. PLOS Comput Biol 14, e1006548.

Hilfinger, A., and Paulsson, J. (2011). Separating intrinsic from extrinsic fluctuations in dynamic biological systems. Proc Natl Acad Sci USA 108, 12167–12172.

Ho, P.-Y., and Amir, A. (2015). Simultaneous regulation of cell size and chromosome replication in bacteria. Front Microbiol 6, 662.

Hochegger, H., Takeda, S., and Hunt, T. (2008). Cyclin-dependent kinases and cell-cycle transitions: does one fit all? Nat Rev Mol Cell Biol 9, 910–916.

Jones, A.R., Forero-Vargas, M., Withers, S.P., Smith, R.S., Traas, J., Dewitte, W., and Murray, J.A.H. (2017). Cell-size dependent progression of the cell cycle creates homeostasis and flexibility of plant cell size. Nat Commun 8, 15060.

Keifenheim, D., Sun, X., D’Souza, E., Ohira, M.J., Magner, M., Mayhew, M.B., Marguerat, S., and Rhind, N. (2017). Size-dependent expression of the mitotic activator Cdc25 suggests a mechanism of size control in fission yeast. Curr Biol 27, 1491–1497.

Lin, J., and Amir, A. (2018). Homeostasis of protein and mRNA concentrations in growing cells. Nat Commun 9, 4496.

Logsdon, M.M., Ho, P.Y., Papavinasasundaram, K., Richardson, K., Cokol, M., Sassetti, C.M., Amir, A., and Aldridge, B.B. (2017). A parallel adder coordinates mycobacterial cell-cycle progression and cell-size homeostasis in the context of asymmetric growth and organization. Curr Biol 27, 3367–3374.

Micali, G., Grilli, J., Marchi, J., Osella, M., and Cosentino Lagomarsino, M. (2018a). Dissecting the control mechanisms for DNA replication and cell division in *E. coli*. Cell Rep 25, 761.

Micali, G., Grilli, J., Osella, M., and Lagomarsino, M.C. (2018b). Concurrent processes set E. coli cell division. Sci Adv 4, eaau3324.

Newman, J.R., Ghaemmaghami, S., Ihmels, J., Breslow, D.K., Noble, M., DeRisi, J.L., and Weissman, J.S. (2006). Single-cell proteomic analysis of *S. cerevisiae* reveals the architecture of biological noise. Nature 441, 840–846.

Nordholt, N., van Heerden, J.H., and Bruggeman, F.J. (2019). Growth-rate and protein-synthesis dynamics of single Bacillus subtilis cells along their cell-cycle. bioRxiv.

Osella, M., Nugent, E., and Cosentino Lagomarsino, M. (2014). Concerted control of *Escherichia coli* cell division. Proc Natl Acad Sci USA 111, 3431–3435.

Padovan-Merhard, O., Nair, G.P., Biaesch, A.G., Mayer, A., Scarfone, S., Foley, S.W., Wu, A.R., Churchman, L.S., Singh, A., and Raj, A. (2015). Single mammalian cells compensate for differences in cellular volume and DNA copy number through independent global transcriptional mechanisms. Mol Cell 58, 339–352.

Pan, K.Z., Saunders, T.E., Flor-Parra, I., Howard, M., and Chang, F. (2014). Cortical regulation of cell size by a sizer cdr2p. eLife 3.

Patterson, J.O., Rees, P., and Nurse, P. (2019). Noisy cell-size correlated expression of Cyclin B drives probabilistic cell-size homeostasis in fission yeast. Curr Biol 29, 1379–1386.

Schmoller, K.M., and Skotheim, J.M. (2015). The biosynthetic basis of cell size control. Trends Cell Biol 25, 793–802.

Schmoller, K.M., Turner, J.J., Koivomagi, M., and Skotheim, J.M. (2015). Dilution of the cell cycle inhibitor Whi5 controls budding-yeast cell size. Nature 526, 268–272.

Scofield, S., Jones, A., and Murray, J.A.H. (2014). The plant cell cycle in context. J Exp Bot 65, 2557–2562.

Sekar, K., Rusconi, R., Sauls, J.T., Fuhrer, T., Noor, E., Nguyen, J., Fernandez, V.I., Buffing, M.F., Berney, M., Jun, S., et al. (2018). Synthesis and degradation of FtsZ quantitatively predict the first cell division in starved bacteria. Mol Syst Biol 14, e8623.

Shi, H., Colavin, A., Bigos, M., Tropini, C., Monds, R.D., and Huang, K.C. (2017). Deep phenotypic mapping of bacterial cytoskeletal mutants reveals physiological robustness to cell size. Curr Biol 27, 3419–3429.

Si, F., Treut, G.L., Sauls, J.T., Vadia, S., Levin, P.A., and Jun, S. (2019). Mechanistic origin of cell-size control and homeostasis in bacteria. Curr Biol 29, 1–11.

Soifer, I., Robert, L., and Amir, A. (2016). Single-cell analysis of growth in budding yeast and bacteria reveals a common size regulation strategy. Curr Biol 26, 356–361.

Swaffer, M.P., Jones, A.W., Flynn, H.R., Snijders, A.P., and Nurse, P. (2016). CDK substrate phosporylation and ordering in the cell cycle. Cell 167, 1750–1761.

Taheri-Araghi, S., Bradde, S., Sauls, J.T., Hill, N.S., Levin, P.A., Paulsson, J., Vergassola, M., and Jun, S. (2015). Cell-size control and homeostasis in bacteria. Curr Biol 25, 385–391.

Turner, J.J., Ewald, J.C., and Skotheim, J.M. (2012). Cell size control in yeast. Curr Biol 22, 350–359.

Varsano, G., Wang, Y., and Wu, M. (2017). Probing Mammalian Cell Size Homeostasis by Channel-Assisted Cell Reshaping. Cell Rep 20, 397–410.

Wallden, M., Fange, D., Lundius, E.G., Baltekin, Ö., and Elf, J. (2016). The synchronization of replication and division cycles in individual *E. coli* cells. Cell 166, 729–739.

Wang, P., Robert, L., Pelletier, J., Dang, W.L., Taddei, F., Wright, A., and Jun, S. (2010). Robust growth of *Escherichia coli*. Curr Biol 20, 1099–1103.

Willis, L., and Huang, K. (2017). Sizing-up the bacterial cell cycle. Nat Rev Microbiol 15, 606–620.

Willis, L., Refahi, Y., Wightman, R., Landrein, B., Teles, J., Huang, K.C., Meyerowitz, E.M., and Jönsson, H. (2016). Cell size and growth regulation in the *Arabidopsis thaliana* apical stem cell niche. Proc Nat Acad Sci 113, E8238–8246.

