## Supplementary material for "Limits and constraints on mechanisms of cell-cycle regulation imposed by cell sizehomeostasis measurements": SI

### 1. Models

In this section, we extend the model in the main manuscript and specify additional notation that is used throughout SI. Parameter and variable definitions are summarized in Table 1. The simpler model presented in the main manuscript allowed the corresponding notation to be simplified. Mostly, the relationships between parameters in main manuscript and the SI are self-explanatory; Table 2 is provided as an additional reference.

We present mechanistic models for cell cycles with two checkpoints, G1/S and G2/M, where either G1/S or G2/M progression is triggered by the attainment of a threshold level or concentration of a particular regulator. The models allow for: (i) variation in the nature of the production of the regulator; (ii) variation in division patterns, for example asymmetric division or binary fission; (iii) variation in growth kinetics during each phase, for example, growth can be linear or exponential in either phase at a rate that varies with cell size at the beginning of the phase; (iv) different phenomenological mechanisms of control of the other phase, either sizer, timer, or adder behavior, or intermediate between these cases—such phases are said to be *independently regulated*; (v) different levels of noise in cell cycle processes over different phases; (vi) in later sections, different criteria for G1/S or G2/M progression. We derive expressions for the linear regression slopes relating initial cell size, final cell size, and phase duration for each cell cycle phase in terms of model parameters. These statistics have been or can be experimentally determined for various cell types.

Two mechanisms of cell cycle regulation are considered. In the first mechanism, a *master regulator* takes a fixed size-independent initial value then triggers cell cycle progression upon reaching a threshold total intracellular level when it is degraded. When a master regulator is produced through both cell-cycle phases rather than through only the phase it regulates, it is a *two-phase master regulator*. In the second mechanism, an *inhibitor dilutor* is produced during one phase then is diluted-out in the next without degradation, triggering cell cycle progression at a minimum threshold concentration. Growth and inhibitor dilutor or master regulator production are modeled by the following equations which describe the average evolution of cell size ($S$) and total intracellular regulator level ($C_i$) during phase $j$:

$$\frac{dS}{dt} = \gamma_j(1+\alpha_{g,j}\Delta S_j)\, S^{\lambda_{g,j}} \tag{1}$$

$$\frac{dC_i}{dt} = \kappa_{ij}(1+\alpha_{c,ij}\Delta S_j)\, S^{\lambda_{c,ij}}. \tag{2}$$

where $C_i$ triggers phase $i$ progression according to a particular criterion. Throughout the text, phase 1 corresponds to G1 and phase 2 corresponds to S/G2/M, and parameters are defined in Table 1. (In the main manuscript, the subscripts $i$ and $j$ are omitted to simplify the exposition, and whether the regulator controls G1/S or G2/M and the relevant growth phase ($j$) are stated explicitly.) Further extending the model in the main manuscript, growth and regulator production through phase $j$ may have a constant dependence on cell size at the beginning of the phase, which is denoted $S_j$ with steady state mean $\mu_{S,j}$ and scaled deviation $\Delta S_j = S_j/\mu_{S,j} - 1$, according to parameters $\alpha_{g,j}$ and $\alpha_{c,ij}$, respectively.

Together equations (1) and (2) imply that the overall change in cell size during phase $j$ is related to the overall change in regulator $i$ by $\Delta(S^{\lambda_{c,ij}-\lambda_{g,j}+1})/(\lambda_{c,ij}-\lambda_{g,j}+1) \approx (1+(\alpha_{g,j}-\alpha_{c,ij})\Delta S_j)\,\frac{\gamma_j}{\kappa_{ij}}\Delta C_i$, so three key parameter combinations govern the relationship between average regulator and growth evolution during phase $j$: $\Delta\alpha_{ij} = \alpha_{g,j} - \alpha_{c,ij}$ encapsulates the initial cell size dependence that is constant throughout the phase, $\Delta\lambda_{ij} = \lambda_{g,j} - \lambda_{c,ij}$ encapsulates the dependence on cell size as cell size increases throughout the phase, while $\gamma_j \Delta(C_i)/\kappa_{ij}$ is independent of cell size. We note that growth is proportional to regulator production through phase $j$ when $\Delta\lambda_{ij} = 0, \Delta\alpha_{ij} = 0$. A master regulator of phase $i$ triggers phase $i$ progression at a threshold level $\Theta_i$, when it is degraded to level $\Theta_{\text{deg}}$. An inhibitor dilutor triggers phase $i$ progression at a threshold minimum concentration $k_i$.

Upon G2/M or equivalently division (phase 2 exit), cells divide into sisters of size ratio 1:$\sigma - 1$. Only the sister corresponding to size ratio 1 is retained in the subsequent generation. Thus, $\sigma = 2$ corresponds to binary fission, while $\sigma > 2$ or $\sigma < 2$ corresponds, for example, to cells repeatedly budding-off larger or smaller sisters, respectively, and at steady state where G1/S and G2/M sizes and regulator levels fluctuate about mean fixed values over generations, the average fold size increase is $\sigma$. Regulators that persist through divisions are assumed to be inherited in proportion to daughter cell size. The average fold increase in cell size over phase $j$ is denoted $\sigma_j = \mu_{S,j+1}/\mu_{S,j}$. Thus, at steady state, the total average fold increase in cell size over the cell cycle is $\sigma = \sigma_1\sigma_2$, and if $\tau$ is the fraction of the cell cycle spent in G1 and growth is exponential ($\lambda_g = 1$), $\sigma_1 = \sigma^\tau$.

We consider master regulators or inhibitor dilutors in combination with various phenomenological controls over phase 1 or phase 2, including sizer, adder, or timer control, meaning that over the phase in question cells reach a critical size, add a fixed size increment, or a fixed time period elapses. Specifically, cell size at the end of phase $i$ ($S_{i+1}$) is determined by cell size at the beginning of the phase ($S_i$) according to

$$S_{i+1} = f_{S,i}S_i + (\sigma_i - f_{S,i})\mu_{S,i}$$

where $f_{S,i}$ is the mode of control (in the main text, $f_{\text{phase}} = f_{S,i} = 0$). Such phases, where cell size at the end of the phase depends only on cell size at the beginning of the phase and on no other preceding transition sizes, are said to be *independently regulated.*

Through each phase of the regulator's production, noise terms cause deviations from the average growth–regulator relationship specified by equations (1) and (2) and from the criteria for cell cycle checkpoint progression according to the simulations and analyses described below.

### 2. Relationships among parameters and average output variables

A convenient but confusing aspect of the model is that the natural choices for free parameters change as cell cycle control patterns change. For example, if an independently regulated phase is under timer control, a natural free parameter is the duration of the phase, while depending on the control type of the other phase, average cell size is an output variable. By contrast, if the independently regulated phase were critical size or equivalently sizer, a natural free parameter is the average cell size at the transition, while the duration of the phase is an output. Regardless of which parameters are natural to consider as free, at steady state, certain expressions connecting parameters related to average cellular behaviors must be satisfied, and these expressions are used to simplify the simulations and analyses.

The expressions are as follows. Equations (1) and (2) imply:

$$\begin{aligned} C_i(T_j) - C_i(T_{j-1}) &= \frac{\kappa_{ij}}{\gamma_j} \cdot \frac{1+\alpha_{c,ij}\Delta S_j}{1+\alpha_{g,j}\Delta S_j} \cdot \frac{S_{j+1}^{\lambda_{c,ij}-\lambda_{g,j}+1} - S_j^{\lambda_{c,ij}-\lambda_{g,j}+1}}{\lambda_{c,ij}-\lambda_{g,j}+1} \\ &= \frac{\kappa_{ij}\,\mu_{S,j}^{1-\Delta\lambda_{ij}}}{\gamma_j} \cdot \frac{1+\alpha_{c,ij}\Delta S_j}{1+\alpha_{g,j}\Delta S_j} \cdot \frac{(S_{j+1}/\mu_{S,j})^{1-\Delta\lambda_{ij}} - (S_j/\mu_{S,j})^{1-\Delta\lambda_{ij}}}{1-\Delta\lambda_{ij}} \end{aligned}$$

where $T_j$ is the time of exit from phase $j$. Taking expectations on either side and approximating we have

$$\mu_{\Delta,ij} \approx \frac{\kappa_{ij}\,\mu_{S,j}^{1-\Delta\lambda_{ij}}(\sigma_j^{1-\Delta\lambda_{ij}}-1)}{\gamma_j(1-\Delta\lambda_{ij})}, \tag{3}$$

where $\mu_{\Delta,ij}$ is the mean change over phase $j$ in the regulator triggering phase $i$. By definition,

$$\text{for master regulators:}\quad \Theta_1 - \Theta_{\rm deg}/\sigma = \mu_{\Delta,12}/\sigma + \mu_{\Delta,11} \quad\text{or}\quad \Theta_2 - \Theta_{\rm deg} = \mu_{\Delta,21} + \mu_{\Delta,22}, \tag{4}$$

or

$$\text{for inhibitor dilutors:}\quad k_1 = \frac{\mu_{\Delta,12}}{(\sigma-1)\mu_{S,2}}, \quad k_2 = \frac{\mu_{\Delta,21}}{(\sigma-1)\mu_{S,1}} \tag{5}$$

and recall that

$$\sigma = \sigma_1\sigma_2, \quad \mu_{S,2} = \sigma_1\mu_{S,1}. \tag{6}$$

These equations can be used to find the output variables, for example either $\mu_{S,1}$ or $\sigma_1$, in terms of the free parameters. They allow us to work in terms of $\sigma_1 \approx \sigma^\tau$, where $\tau$ is the fraction of the cell cycle taken up by G1 and the approximation is exact for exponential growth, regardless of the the modes of cell cycle regulation and thus to simulate cell lineage trajectories in Figs. 2 and 3 of the main manuscript. Regardless of whether $\sigma_1$ is a free parameter or an output variable, it is a convenient parameter to work with: it is required that $\sigma_1 > 1$ and $\sigma_1 < \sigma$ in order for the corresponding cell cycle to have two phases of non-zero duration. Thus, exploring parameter space only where $\sigma_1$ satisfies these inequalities allows us to automatically exclude parameter sets which fail to implement cell cycles with two checkpoints on average. For example, the following scenario is excluded: critical size control of phase 2 combined with inhibitor dilutor control of phase 1 where inhibitor production and the threshold minimum concentration are such that the average cell size at the end of phase 1 is less than the average birth size.

### 3. Stochastic simulations

Here, we describe how noise is incorporated into simulations of cell lineages with average patterns of growth and cell cycle regulation as described in the preceding section. In later sections, we compute corresponding analytical results to obtain explicit expressions for cell size-homeostasis behaviors in terms of model parameters.

In all simulations of cell lineages, cell sizes and regulator levels are initialized at their steady state mean values plus noise. At consecutive cell cycle transitions, cell sizes and regulator levels are simulated according to whether the transition is triggered by a master regulator (one-phase or two-phase), an inhibitor dilutor, or an independently regulated phase, as described below. Upon division, one daughter cell is retained with birth size $S_1 = S_3'/\sigma$ where $S_3'$ is cell size at division in the preceding generation, and if the regulator persists through the division (e.g. inhibitor dilutors or two-phase master regulators triggering phase 1 exit), it is split between daughters in proportion to their respective sizes. Thus, we assume no noise in division-plane positioning. Given cell sizes at the beginning and end of phase $j$, $S_j$ and $S_{j+1}$, respectively, the duration of phase $j$, $t_j$, is simulated by solving (1):

$$t_j = \frac{1}{1+\alpha_{g,j}\Delta S_j}\,\frac{S_{j+1}^{1-\lambda_{g,j}} - S_j^{1-\lambda_{g,j}}}{\gamma_i(1-\lambda_{g,j})} \rightarrow \frac{1}{1+\alpha_{g,j}\Delta S_j}\,\frac{\ln(S_{j+1}/S_j)}{\gamma_j} \text{ as } \lambda_{g,j} \rightarrow 1.$$

assuming no noise in growth coefficient $\gamma_j$ beyond the fluctuations due to deviations in initial cell size ($\Delta S_j$). Cell lineages are simulated over many generations until cell sizes and regulator levels have patently reached steady states.

If a *one-phase master regulator* triggers exit from phase $i$, cell size at the end of the phase ($S_{i+1}$) is simulated by solving equations (1) and (2) (which represent the average growth–regulator relationship) over phase $i$, adding a noise term $\mu_{S,i+1}Z_i$ into this relationship where $Z_i \sim N(0,\zeta_i)$ has the coefficient $\mu_{S,i+1}$ to simplify the exposition:

$$S_{i+1} = \Big(S_i^{1-\Delta\lambda_{ii}} + \frac{(\Theta_i - \Theta_{\text{deg}} + \mu_{\Delta,i\,i-1}Z_{CL})\,\gamma_i(1-\Delta\lambda_{ii})}{\kappa_{ii}}\frac{H(1+\alpha_{g,i}\Delta S_i)}{H(1+\alpha_{c,ii}\Delta S_i)}\Big)^{1/(1-\Delta\lambda_{ii})} + \mu_{S,i+1}Z_i$$

$$\to \quad S_{i+1} = S_i\,\exp\Big(\frac{(\Theta_i - \Theta_{\text{deg}} + \mu_{\Delta,i\,i-1}Z_{CL})\,\gamma_i}{\kappa_{ii}}\frac{H(1+\alpha_{g,i}\Delta S_i)}{H(1+\alpha_{c,ii}\Delta S_i)}\Big) + \mu_{S,i+1}Z_i \text{ as } \Delta\lambda_{ii}\to 1$$

where $\mu_{\Delta,i\,i-1}Z_{CL}$ is the noise in the threshold level, $Z_{CL} \sim N(0,\zeta_{CL})$, and

$$H(x) = \begin{cases} c & \text{if } x < c \\ x & \text{if } c \le x \ge 2-c \\ 2-c & \text{if } x > 2-c \end{cases}$$

with $c = 10^{-6}$ prevents growth and regulator production rates ($\gamma_j(1+\alpha_{g,i}\Delta S_i)$ and $\kappa_{ij}(1+\alpha_{g,i}\Delta S_i)$, respectively) being negative. Since the regulator is not simulated explicitly, there is no initial regulator value in generation 1. For one-phase master regulators, $\mu_{\Delta,i\,i} = \Theta_i - \Theta_{\text{deg}}$ while $\mu_{\Delta i\,i-1}$ is arbitrary; here, $\mu_{\Delta i\,i-1}$ is used as the coefficient of $Z_{CL}$ to simplify the exposition later. Throughout, the coefficients of noise terms were chosen to simplify the exposition.

If a *two-phase master regulator* triggers exit from phase $i$, then solving equations (1) and (2) and adding a noise term $\mu_{S,i+1}Z_i$, where $Z_i \sim N(0,\zeta_i)$, we have

$$S_{i+1} = \Big(S_i^{1-\Delta\lambda_{ii}} + \frac{(\Theta_i - C_{ii} + \mu_{\Delta,i\,i-1}Z_{CL})\,\gamma_i(1-\Delta\lambda_{ii})}{\kappa_{ii}}\frac{H(1+\alpha_{g,i}\Delta S_i)}{H(1+\alpha_{c,ii}\Delta S_i)}\Big)^{1/(1-\Delta\lambda_{ii})} + \mu_{S,i+1}Z_i$$

$$\to \quad S_{i+1} = S_i\,\exp\Big(\frac{(\Theta_i - C_{ii} + \mu_{\Delta,i\,i-1}Z_{CL})\,\gamma_i}{\kappa_{ii}}\frac{H(1+\alpha_{g,i}\Delta S_i)}{H(1+\alpha_{c,ii}\Delta S_i)}\Big) + \mu_{S,i+1}Z_i \text{ as } \Delta\lambda_{ii}\to 1$$

where $C_{ii}$ is the regulator's level at the beginning of phase $i$, and other notation is as for one-phase master regulators. To determine the value of $C_{ii}$, the master regulator's dynamics must be simulated for the preceding phase. For a master regulator that triggers phase 2 exit, the regulator's value at the beginning of phase 2 is determined by solving (1) and (2) with initial regulator value at the beginning of phase 1 being $C_{21} = \Theta_{\text{deg}}$:

$$C_{22} = \Theta_{\text{deg}} + \frac{\kappa_{21}}{\gamma_1} \cdot \frac{H(1 + \alpha_{c,21}\Delta S_1)}{H(1 + \alpha_{g,1}\Delta S_1)} \cdot \frac{(S_2 - \mu_{S,2}Z_1)^{1-\Delta\lambda_{21}} - S_1^{1-\Delta\lambda_{21}}}{1 - \Delta\lambda_{21}}, \quad Z_1 \sim N(0, \zeta_1)$$

where $S_2$ is the cell size at the end of the phase 1 or beginning of phase 2, and again noise in the average growth and regulator relationship is according to $Z_1 \sim N(0, \zeta_1)$. For a two-phase master regulator that triggers exit from phase 1, the regulator's value at the beginning of phase 1 is determined similarly but taking into account the intervening division whereupon the regulator is split between daughters in proportion to their respective sizes, with the initial regulator value at the beginning of the phase 2 being $C_{12} = \Theta_{\text{deg}}$:

$$C_{11} = 1/\sigma \times \Big(\Theta_{\text{deg}} + \frac{\kappa_{12}}{\gamma_2} \cdot \frac{H(1 + \alpha_{c,12}\Delta S_2')}{H(1 + \alpha_{g,2}\Delta S_2')} \cdot \frac{(S_3' - \mu_{S,3}Z_2')^{1-\Delta\lambda_{12}} - S_2'^{1-\Delta\lambda_{12}}}{1 - \Delta\lambda_{12}}\Big)$$

where $S_3'$ is the cell size at the end of phase 2 at the preceding cell cycle, and $\mu_{S,3} = \mu_{S,1}\sigma$ is the average cell size at this stage.

If an *inhibitor-dilutor* triggers exit from phase $i$,

$$S_{i+1} = (C_{ii} + \mu_{\Delta,i\,i-1}Z_{CL})/k_i + \mu_{S,i+1}Z_{ID}, \quad Z_{ID} \sim N(0, \zeta_{ID})$$

where the inhibitor is diluted-out during phase $i$ to reach a threshold minimum concentration $k_i$ with noise $Z_{ID}$, $C_{ii}$ is the inhibitor's level at the beginning of phase $i$, and $C_{ii} + \mu_{\Delta i\,i-1}Z_{CL}$ is the inhibitor's level at the end of phase $i$. For an inhibitor-dilutor that triggers phase 2 exit, the inhibitor's value accumulates through phase 1 so that at the beginning of phase 2 it is determined by solving (1) and (2) with initial value at the beginning of phase 1 being determined by its value at the previous cell cycle, that is, $C_{21} = C_{23}'/\sigma = (C_{22}' + \mu_{\Delta,21}Z_{CL}')/\sigma$ (since we assume no degradation on average during the dilution), thus at the beginning of phase 2

$$\begin{aligned} C_{22} &= (C'_{22} + \mu_{\Delta,21} Z'_{CL})/\sigma + \frac{\kappa_{21}}{\gamma_1} \cdot \frac{H(1+\alpha_{c,21}\Delta S_1)}{H(1+\alpha_{g,1}\Delta S_1)} \cdot \frac{(S_2 - \mu_{S,2} Z_1)^{1-\Delta\lambda_{21}} - S_1^{1-\Delta\lambda_{21}}}{1-\Delta\lambda_{21}} \\ &= k_2(S'_3 - \mu_{S,3} Z'_{ID})/\sigma + \frac{\kappa_{21}}{\gamma_1} \cdot \frac{H(1+\alpha_{c,21}\Delta S_1)}{H(1+\alpha_{g,1}\Delta S_1)} \cdot \frac{(S_2 - \mu_{S,2} Z_1)^{1-\Delta\lambda_{21}} - S_1^{1-\Delta\lambda_{21}}}{1-\Delta\lambda_{21}} \\ &= k_2(S_1 - \mu_{S,1} Z'_{ID}) + \frac{\kappa_{21}}{\gamma_1} \cdot \frac{H(1+\alpha_{c,21}\Delta S_1)}{H(1+\alpha_{g,1}\Delta S_1)} \cdot \frac{(S_2 - \mu_{S,2} Z_1)^{1-\Delta\lambda_{21}} - S_1^{1-\Delta\lambda_{21}}}{1-\Delta\lambda_{21}} \end{aligned}$$

where $S_2$ is the cell size at the beginning of phase 2, $S'_3$ is the cell size at the end of phase 2 of the previous cell cycle, and $Z_1$ is the noise between growth and regulator production over phase 1. For an inhibitor-dilutor that triggers exit from phase 1, the inhibitor's value at the beginning of phase 1 is determined similarly:

$$\begin{aligned} C_{11} &= 1/\sigma \times \Big( C'_{11} + \mu_{\Delta,12} Z'_{CL} + \frac{\kappa_{12}}{\gamma_2} \cdot \frac{H(1+\alpha_{c,12}\Delta S'_2)}{H(1+\alpha_{g,2}\Delta S'_2)} \cdot \frac{(S'_3 - \mu_{S,3} Z'_2)^{1-\Delta\lambda_{12}} - S'^{1-\Delta\lambda_{12}}_2}{1-\Delta\lambda_{12}} \Big) \\ &= 1/\sigma \times \Big( k_1(S'_2 - \mu_{S,2} Z'_{ID}) + \frac{\kappa_{12}}{\gamma_2} \cdot \frac{H(1+\alpha_{c,12}\Delta S'_2)}{H(1+\alpha_{g,2}\Delta S'_2)} \cdot \frac{(S'_3 - \mu_{S,3} Z'_2)^{1-\Delta\lambda_{12}} - S'^{1-\Delta\lambda_{12}}_2}{1-\Delta\lambda_{12}} \Big). \end{aligned}$$

If phase $i$ is *independently regulated* as a timer, adder, sizer, or intermediate between these cases, then cell size upon exit from phase $i$ is simulated by

$$S_{i+1} = f_{S,i} S_i + \mu_{S,i+1} - f_{S,i}\, \mu_{S,i} + \mu_{S,i+1}\, Z_O \quad Z_O \sim N(0, \zeta_O),$$

where $f_{S,i}$ determines the mode of regulation: $f_{S,i} = 0$ or 1 produces sizer or adder regulation, and from Eq. (29), $f_{S,i} = \sigma_i^{\lambda_{g,i}} + \alpha_{g,i}(\sigma_i - \sigma_i^{\lambda_{g,i}})/(1-\lambda_{g,i})$ produces approximate timer regulation. The noise term $Z_O$ is the noise in cell size acquired over the independent phase.

In all simulations $\mu_{S,1} = 1, \gamma_1 = 1$, and $\mu_{\Delta,ij} = 1$ for a specific $i, j$ sets the units of size, time, and regulator level, respectively. Parameters $\Delta\lambda_{ij}, \frac{\kappa_{ij}}{\gamma_j(1-\Delta\lambda_{ij})}, \sigma, k_i, \mu_{\Delta,ij}, \mu_{S,i}$ are interrelated according to Eqs. (3)–(6). Unless otherwise stated, master regulators are assumed to be completely degraded after triggering a transition, i.e. $\Theta_{\text{deg}} = 0$. Throughout, simulation errors are avoided by forcing cells to increase their size above a minimum of $10^{-6}$ and forcing regulator levels to stay above $10^{-6}$ over each phase. Our analytical derivations, which assume small cell size fluctuations while ignoring this enforced minimum growth, are closely matched by simulations with realistic noise terms, which strongly implies that the enforced minimum growth has negligible influence on simulated statistics for realistic parameter values.

In summary, noise terms cause: (1) deviations from the average growth—regulator relationship prescribed by a given parameter set ($\zeta_j$); (2) deviations in the threshold minimum concentration for checkpoint progression of inhibitor dilutors ($\zeta_{ID}$); (3) deviation in final level of regulators upon checkpoint progression, and thus in the threshold level for checkpoint progression for master regulators, or in the dilution phase for inhibitor dilutors ($\zeta_{CL}$); (4) deviations in the criterion for checkpoint progression in independently regulated phases ($\zeta_O$). Noise terms are assumed to be *independent* and thus are not affected by cell sizes or regulator levels at preceding transitions.

#### 3.1. **Nomenclature.** See Tables 1 and 3.

### 4. Interpretations of noise terms and corresponding measurements

Two noise terms, one associated with independently regulated phases ($Z_O$, standard deviation $\zeta_O$) and one with inhibitor dilutors ($Z_{ID}$, standard deviation $\zeta_{ID}$), have particularly important implications for the main manuscript. The terms

$$\eta_{\text{checkpoint}} = \frac{\text{Noise in checkpoint mechanism}}{\text{Coefficient of variation (CV) in G1/S size}}$$

where 'Noise in checkpoint mechanism' corresponds either to $\zeta_O$ or $\zeta_{ID}$. We discuss the meanings of these noise terms and how they can be measured, under the assumptions of the model.

For independently regulated phases, the interpretation of the noise term depends on the mode of control of the independently regulated phase. Recall that $\Delta S_i = S_i/\mu_{S,i} - 1$, then from the definition of an independently regulated phase:

$$\Delta S_{i+1} = \frac{f_{S,i}}{\sigma_i}\Delta S_i + Z_O.$$

Table 1. Notation.

| | |
|---|---|
| $\mu_{S,j}$ | Average cell size at the beginning of phase $j$. |
| $S$, $S_j$, $\Delta S_j$ | Cell size, at the beginning of phase j, scaled fluctuations $S_j/\mu_{S,j} - 1$ at the beginning of phase j. |
| $C_i$, $C_{ij}$ | Regulator that triggers exit from phase i, its level at the beginning of phase j. |
| $\mu_{\Delta,ij}$ | Average change in regulator $C_i$ over phase $j$. |
| $\Delta_{ij}$ | Scaled fluctuations in the change in regulator $C_i$ over phase $j$, $(C_{i,j+1} - C_{ij})/\mu_{\Delta,ij} - 1$. |
| $\sigma_j$, $\sigma$ | Average fold increase in size over phase $j$, average fold increase in size over the cell cycle. |
| $\Theta_i$, $\Theta_{\text{deg}}$ | Master regulators: threshold level for regulator $C_i$ that triggers phase $i$ progression, initial fixed value of regulator. |
| $k_i$ | Inhibitor-dilutors: the minimum threshold concentration that triggers progression through phase $i$. |
| $t_j$, $T_j$ | Duration, time of exit of phase $j$. |
| $Z_j$, $\zeta_j$ | Noise and standard deviation in coordination of growth and regulator production over phase $j$. |
| $Z_O$, $\zeta_O$ | Noise and standard deviation in the independently regulated phase, e.g. the sizer, adder, or timer mechanism. |
| $Z_{CL}$, $\zeta_{CL}$ | Noise and standard deviation in the regulator's final level that triggers phase progression. |
| $Z_{ID}$, $\zeta_{ID}$ | Noise and standard deviation in an inhibitor dilutor's threshold minimum concentration for phase progression. |
| $\Delta\alpha_{ij} = \alpha_{g,j} - \alpha_{c,ij}$ | Deviation from proportional growth and regulator $C_i$ production via initial cell size in phase $j$. |
| $\Delta\lambda_{ij} = \lambda_{g,j} - \lambda_{c,ij}$ | Deviation from proportional growth and regulator $C_i$ production via cell size in phase $j$. |
| $\kappa_{ij}, \gamma_j$ | Constants of proportionality for regulator $C_i$ production, growth rate in phase $j$. |

By taking variances either side of the preceding equation, we have

$$\text{Var}(\Delta S_{i+1}) = \left(\frac{f_{S,i}}{\sigma_i}\right)^2 \text{Var}(\Delta S_i) + \zeta_O^2$$

where the variance of $\Delta S_i$ equals the variance of $S_i/\mu_{S,i}$, which equals the square of the coefficient of variation of $S_i$. If control is 'sizer' or critical size, then $f_{S,i} = 0$ and the standard deviation $\zeta_O$ equals *the coefficient of cell size at the end of the independently regulated phase.* This is a measurable statistic. For exponential growth and timer regulation, it is more appropriate for the

TABLE 2. Notation that is labelled differently in the manuscript compared with the supplement (SI). LSLS stands for least-square linear slope, equivalent to the linear regression slope.

| Manuscript | SI | |
|---|---|---|
| $l_{\mathrm{G1}}$, $f_{\mathrm{G1}}$ | $f_{S,1}$ | LSLS between birth size vs. size at the G1/S transition, which for independently regulated phases is equivalent to the mode of control e.g. sizer, adder, etc. |
| $l_{\mathrm{S/G2/M}}$, $f_{\mathrm{S/G2/M}}$ | $f_{S,2}$ | LSLS between size at G1/S vs. size at G2/M or division, which for independently regulated phases is equivalent to the mode of control. |
| $\sigma_{\mathrm{G1}}, \sigma_{\mathrm{S/G2/M}}$ | $\sigma_1, \sigma_2$ | The average fold-size increase over G1 or S/G2/M. |
| $\mu_{i,\mathrm{G1}}$, $\mu_{i,\mathrm{S/G2/M}}$ | $\mu_{S,1}$, $\mu_{S,2}$ | The average cell size at the beginning of G1 or S/G2/M. |
| $\zeta_{C,G1}, \zeta_{C,S/G2/M}$ | $\zeta_1, \zeta_2$ | The standard deviations in coordination of growth and regulator production over G1 or S/G2/M. |
| $\zeta_{G1/S}, \zeta_{G2/M}$ | $\zeta_{ID}$ or $\zeta_O$ | The standard deviations in checkpoint noise, e.g. $\zeta_{G1/S} = \zeta_{ID}$ and $\zeta_{G2/M} = \zeta_O$ if an inhibitor dilutor controls G1/S while an independently regulated phase controls S/G2/M. |

TABLE 3. Statistics. LSLS stands for least-square linear slope, equivalent to the linear regression slope.

| | |
|---|---|
| $f_{ii}$ | LSLS between cell sizes at the beginning of phase $i$ in consecutive cell cycles. |
| $f_{S,i}$ | LSLS between cell size at the beginning vs. cell size at the end of phase $i$. |
| $f_{t,i}$ | LSLS between scaled cell size fluctuations at the beginning of phase $i$ and scaled fluctuations in the duration of phase $i$. |
| $\mathrm{Var}(\Delta S_i)$ | Variance in scaled fluctuations in cell size at the beginning of phase $i$, equal to the square of the coefficient of variation of $S_i$. |

independent random variable to perturb the duration of the phase. Then, assuming no noise in cell growth, cell size at the end of phase $i$ would be written:

$$S_{i+1} = S_i\, \sigma_i^{1+Z_T} \tag{7}$$

where $Z_T$ with standard deviation $\zeta_T$ is called a multiplicative noise term because its effect multiplies with $S_i$. Equation (7) can be written

$$\Delta S_{i+1} = \Delta S_i + (1 + \Delta S_i) \ln \sigma_i Z_T \; + \text{ h.o.t. } = \Delta S_i + \ln \sigma_i Z_T \; + \text{ h.o.t.}$$

so to first order, independently regulated phases encompass the multiplicative noise case. The normalized standard deviation of the phase duration, $\zeta_T$, that is, *the coefficient of variation of the phase's duration*, determines the parameter $\zeta_O = \ln \sigma_i \zeta_T$. For example, if $\sigma_i = 2^{0.5}$ and $\zeta_T = 0.2$, then $\zeta_O = 0.07$. Then, for a typical CV in G1/S size of 0.14, we have $\eta_{\text{checkpoint}} = 0.07/0.14 = 0.5$.

For inhibitor dilutors triggering exit from phase $i$ at a threshold concentration $k_i$, to first order in cell size and inhibitor level fluctuations, the standard deviation of the noise term $Z_{ID}$, $\zeta_{ID}$, is equivalent to the coefficient of variation in $k_i$, as we will now show. From the preceding section, recall that:

$$S_{i+1} = (C_{ii} + \mu_{\Delta,i\,i-1} Z_{CL})/k_i + \mu_{S,i+1} Z_{ID}, \quad Z_{ID} \sim N(0, \zeta_{ID})$$

where by definition at steady state $\mu_{S,i+1} = \mathbb{E}[C_{ii}]/k_i$ where $\mathbb{E}[C_{ii}]$ is the average value of the inhibitor at the beginning of the phase and therefore also at the end of the phase, given no inhibitor degradation. Below $\Delta C_{ii}$ is the scaled deviation from the average of the inhibitor level $C_{ii}$ at the beginning of phase $i$. Since

$$\begin{aligned}
\frac{C_{ii} + \mu_{\Delta,i\,i-1} Z_{CL}}{k_i(1 - Z_{ID})} &= \frac{C_{ii} + \mu_{\Delta,i\,i-1} Z_{CL}}{k_i}(1 + Z_{ID} + \text{h.o.t.}) \\
&= \frac{C_{ii} + \mu_{\Delta,i\,i-1} Z_{CL}}{k_i} + \frac{\mathbb{E}[C_{ii}](1 + \Delta C_{ii}) + \mu_{\Delta,i\,i-1} Z_{CL}}{k_i} Z_{ID} + \text{h.o.t.} \\
&= \frac{C_{ii} + \mu_{\Delta,i\,i-1} Z_{CL}}{k_i} + \frac{\mathbb{E}[C_{ii}]}{k_i} Z_{ID} + \text{h.o.t.} \\
&= \frac{C_{ii} + \mu_{\Delta,i\,i-1} Z_{CL}}{k_i} + \mu_{S,i+1} Z_{ID}
\end{aligned}$$

where all terms corresponding to fluctuations of order 2 or above were dropped. Therefore, the noise term $Z_{ID}$ corresponds to fluctuations in the threshold concentration triggering phase $i$ exit, and regardless of the distribution of $Z_{ID}$, $\zeta_{ID}$ is the CV of the threshold concentration. Therefore $\zeta_{ID}$ is potentially measurable.

### 5. Analytical derivations

In this section, we derive analytical expressions for the least-square linear slopes (LSLS) or equivalently the linear regression slopes between cell sizes and phase durations at different stages of the cell cycle. Throughout we assume small fluctuations about average cell sizes. Simulations with realistic fluctuations, i.e. noise levels that produce cell size coefficients of variation of

0.1–0.2, are in excellent agreement with these derivations, see figures in the main manuscript. Importantly, the analytical derivations *do not depend on how the random variables, representing noise terms, are distributed, but depend only on the standard deviations of the random variables.*

5.1. **Definition of least-square linear slope (LSLS).** We derive a well-known expression for the least-square linear slope (LSLS), or linear regression slope, between two variables $X$ and $Y$, which will be used continually throughout this section. The regression coefficients $a$ and $b$ minimize the value of $\mathbb{E}[(Y-(a+bX))^2]$, so differentiating with respect to $a$, the value of $a$ that gives zero derivative is:

$$a = \mu_Y - b\mu_X$$

where $\mu_X$ and $\mu_Y$ are the means of $X$ and $Y$, respectively. Substituting this value of $a$ into $\mathbb{E}[(Y-(a+bX))^2]$, then differentiating with respect to $b$ to find the value of $b$ that gives zero derivative:

$$b = \frac{\mathbb{E}[(Y-\mu_Y)(X-\mu_Y)]}{\mathbb{E}[(X-\mu_X)^2]},$$

which is by definition the LSLS or linear regression slope.

5.2. **LSLSs between cell sizes at different cell cycle stages.** First we derive an approximation that will be used multiple times in the following sections. Let $\Delta\lambda_{ij} = \lambda_{g,j} - \lambda_{c,ij}$, $\Delta\alpha_{ij} = \alpha_{g,j} - \alpha_{c,ij}$, and because cell size fluctuations are assumed small ($\Delta S_j \ll 1$), together equations (1) and (2) imply:

$$\begin{aligned} C_i(T_j) - C_i(T_{j-1}) &= \frac{\kappa_{ij}}{\gamma_j} \cdot \frac{1+\alpha_{c,ij}\Delta S_j}{1+\alpha_{g,j}\Delta S_j} \cdot \frac{S_{j+1}^{\lambda_{c,ij}-\lambda_{g,j}+1} - S_j^{\lambda_{c,ij}-\lambda_{g,j}+1}}{\lambda_{c,ij}-\lambda_{g,j}+1} \\ &= \frac{\kappa_{ij}\,\mu_{S,j}^{1-\Delta\lambda_{ij}}}{\gamma_j}\left(1-\Delta\alpha_{ij}\Delta S_j + o(\Delta S_j)\right)\frac{(S_{j+1}/\mu_{S,j})^{1-\Delta\lambda_{ij}} - (S_j/\mu_{S,j})^{1-\Delta\lambda_{ij}}}{1-\Delta\lambda_{ij}} \end{aligned}$$

where $T_j$ is the time of exit from phase $j$. This is prior to the addition of noise at the end of phase $j$. Using the approximation,

$$\mu_{\Delta,ij} \approx \frac{\kappa_{ij}\,\mu_{S,j}^{1-\Delta\lambda_{ij}}(\sigma_j^{1-\Delta\lambda_{ij}}-1)}{\gamma_j(1-\Delta\lambda_{ij})}, \qquad (8)$$

if $\mu_{\Delta,ij} \neq 0$, we have

$$\begin{aligned}\frac{C_i(T_j) - C_i(T_{j-1})}{\mu_{\Delta,ij}} &\approx (1 - \Delta\alpha_{ij}\Delta S_j)\frac{(\sigma_j S_{j+1}/\mu_{S,j+1})^{1-\Delta\lambda_{ij}} - (S_j/\mu_{S,j})^{1-\Delta\lambda_{ij}}}{\sigma_j^{1-\Delta\lambda_{ij}} - 1} \\ &= (1 - \Delta\alpha_{ij}\Delta S_j)\,(1 + (1 - \Delta\lambda_{ij})\frac{\sigma_j^{1-\Delta\lambda_{ij}}\Delta S_{j+1} - \Delta S_j}{\sigma_j^{1-\Delta\lambda_{ij}} - 1} + \text{h.o.t.}) \\ &= 1 - \Big(\Delta\alpha_{ij} + \frac{1 - \Delta\lambda_{ij}}{\sigma_j^{1-\Delta\lambda_{ij}} - 1}\Big)\Delta S_j + \frac{(1 - \Delta\lambda_{ij})\sigma_j^{1-\Delta\lambda_{ij}}}{\sigma_j^{1-\Delta\lambda_{ij}} - 1}\Delta S_{j+1} + \text{h.o.t.}\end{aligned}$$

and the scaled deviation of the increment of regulator $i$ is related to the scaled deviations from average cell sizes at the beginning and end of phase $j$ by:

$$\Delta_{ij} = \frac{C_i(T_j) - C_i(T_{j-1})}{\mu_{\Delta,ij}} - 1 \approx -\Big(\Delta\alpha_{ij} + \frac{1 - \Delta\lambda_{ij}}{\sigma_j^{1-\Delta\lambda_{ij}} - 1}\Big)\Delta S_j + \frac{(1 - \Delta\lambda_{ij})\sigma_j^{1-\Delta\lambda_{ij}}}{\sigma_j^{1-\Delta\lambda_{ij}} - 1}\Delta S_{j+1}$$

or, now taking into account the addition of noise $Z_j$ in stochastic simulations,

$$\Delta_{ij} \approx -\Big(\Delta\alpha_{ij} + \frac{1 - \Delta\lambda_{ij}}{\sigma_j^{1-\Delta\lambda_{ij}} - 1}\Big)\Delta S_j + \frac{(1 - \Delta\lambda_{ij})\sigma_j^{1-\Delta\lambda_{ij}}}{\sigma_j^{1-\Delta\lambda_{ij}} - 1}(\Delta S_{j+1} - Z_j). \tag{9}$$

For phase $i$ inhibitor-dilutors, $\mu_{\Delta,ii} = 0$ and

$$C_i(T_i) - C_i(T_{i-1}) = \mu_{\Delta,i\,i-1} Z_{CL}. \tag{10}$$

Expressions (9) and (10) are used in the following derivations. These expressions and all following derivations do not depend on the distributions of random variables in the simulation.

5.2.1. *One-phase master regulators.* For one-phase master regulators governing phase $i$, $1+\Delta_{ii} = (C_i(T_i) - C_i(T_{i-1}))/\mu_{\Delta,ii} = (\Theta_i - \Theta_{\text{deg}} + \mu_{\Delta,i\,i-1}Z_{CL})/\mu_{\Delta,ii} = 1 + \mu_{\Delta,i\,i-1}Z_{CL}/\mu_{\Delta,ii}$, so from (9) we have a relation between scaled fluctuations from the average cell size at the beginning and end of the phase

$$\Delta S_{i+1} = \Big(\sigma_i^{\Delta\lambda_{ii}-1} + \Delta\alpha_{ii}\frac{1 - \sigma_i^{\Delta\lambda_{ii}-1}}{1 - \Delta\lambda_{ii}}\Big)\,\Delta S_i + \text{h.o.t.} + Z_i + \frac{\mu_{\Delta,i\,i-1}}{\mu_{\Delta,ii}}\frac{1 - \sigma_i^{\Delta\lambda_{ii}-1}}{1 - \Delta\lambda_{ii}} Z_{CL}.$$

Therefore, the LSLS between $\Delta S_i$ and $\Delta S_{i+1}$, which is by definition $\mathbb{E}[\Delta S_{i+1}\,\Delta S_i]/\mathbb{E}[\Delta S_i^2]$, is to first order in cell size fluctuations:

$$\text{LSLS of } \Delta S_i \text{ vs. } \Delta S_{i+1} \quad\approx\quad \sigma_i^{\Delta\lambda_{ii}-1} + \Delta\alpha_{ii}\frac{1 - \sigma_i^{\Delta\lambda_{ii}-1}}{1 - \Delta\lambda_{ii}}$$

because $Z_i$ and $Z_{CL}$ are by assumption independent of $\Delta S_i$ so $\mathbb{E}[\Delta S_i Z_i] = \mathbb{E}[\Delta S_i Z_{CL}] = 0$. Multiplying this expression for the LSLS by $\sigma_i = \mu_{S,i+1}/\mu_{S,i}$ gets the LSLS between cell size at the beginning vs. the end of phase $i$ ($S_i$ vs. $S_{i+1}$, denoted $f_{S,i}$):

$$f_{S,i} \approx \sigma_i^{\Delta\lambda_{ii}} + \Delta\alpha_{ii}\frac{\sigma_i - \sigma_i^{\Delta\lambda_{ii}}}{1 - \Delta\lambda_{ii}}. \tag{11}$$

5.2.2. *Independently regulated phases.* For one-phase master regulators, cell size fluctuations depend only on the fluctuation at the previous phase and not on fluctuations at earlier phase transitions. They are examples of *independently regulated phases.* Independently regulated phases are simulated as

$$\Delta S_{i+1} \quad = \quad \frac{f_{S,i}}{\sigma_i}\Delta S_i + Z_i.$$

Multiplying this expression by $\Delta S_i$, taking expectations, dividing by $\mathbb{E}[\Delta S_i^2]$, and multiplying by $\sigma_i$, we find

$$\text{LSLS between cell size at the beginning vs. end of phase } i = f_{S,i}.$$

If the subsequent or preceding phase is also triggered by an independently regulated mechanism, then the LSLS relating size at the beginning of phase $i$ vs. size at the beginning of phase $i$ in the subsequent generation is $f_{S,ii} \approx f_{S,1}f_{S,2}/\sigma$, because, ignoring non-linear terms in cell size fluctuations:

$$\begin{aligned} \Delta S_{i+1} &= \frac{f_{S,i}}{\sigma_i}\Delta S_i + Z_i \\ \Rightarrow \Delta S_{i+2} &= \frac{f_{S,i+1}}{\sigma_{i+1}}\Big(\frac{f_{S,i}}{\sigma_i}\Delta S_i + Z_i\Big) + Z_{i+1} \\ \Rightarrow f_{S,ii} = \frac{\mathbb{E}[\Delta S_{i+2}\Delta S_i]}{\mathbb{E}[\Delta S_i^2]} &= \frac{f_{S,i}f_{S,i+1}}{\sigma_i\sigma_{i+1}} = \frac{f_{S,1}f_{S,2}}{\sigma}. \end{aligned}$$

Multiplying this expression by $\sigma$, we have:

$$\text{LSLS of birth vs. division size} = \sigma\frac{\mathbb{E}[\Delta S_3\,\Delta S_1]}{\mathbb{E}[\Delta S_1^2]} = f_{S,1}\,f_{S,2}.$$

Again we assumed that $Z_i, Z_{i+1}$ are zero-mean random variables independent of $\Delta S_i$.

5.2.3. *Two-phase master regulators.* For a two-phase master regulator that triggers G2/M or division (exit from phase 2), we have:

$$\begin{aligned}\mu_{\Delta,21}\Delta_{21}+\mu_{\Delta,22}\Delta_{22} &= \mu_{\Delta,21}\Big(\frac{C_2(T_1)-C_2(T_0)}{\mu_{\Delta,21}}-1\Big)+\mu_{\Delta,22}\Big(\frac{C_2(T_2)-C_2(T_1)}{\mu_{\Delta,22}}-1\Big)\\ &= \Theta_2+\mu_{\Delta,21}Z_{CL}-\Theta_{\text{deg}}-(\mu_{\Delta,21}+\mu_{\Delta,22})\\ &= \mu_{\Delta,21}Z_{CL}\end{aligned}$$

$$\Rightarrow \qquad \Delta_{22} = -\frac{\mu_{\Delta,21}}{\mu_{\Delta,22}}\Delta_{21}+\frac{\mu_{\Delta,21}}{\mu_{\Delta,22}}Z_{CL} \tag{12}$$

because $C_2(T_0)=\Theta_{\text{deg}}$ and $C_2(T_2)=\Theta_2+\mu_{\Delta,21}Z_{CL}$. Similarly for a two-phase master regulator that triggers G1/S (exit from phase 1):

$$\begin{aligned}\mu_{\Delta,12}\Delta_{12}+\sigma\mu_{\Delta,11}\tilde{\Delta}_{11} &= \mu_{\Delta,12}\Big(\frac{C_1(T_2)-C_1(T_1)}{\mu_{\Delta,12}}-1\Big)+\sigma\mu_{\Delta,11}\Big(\frac{C_1(\tilde{T}_1)-C_1(\tilde{T}_0)}{\mu_{\Delta,11}}-1\Big)\\ &= \sigma(\Theta_1+\mu_{\Delta,12}Z_{CL})-\Theta_{\text{deg}}-(\mu_{\Delta,12}+\sigma\mu_{\Delta,11})\\ &= \sigma\mu_{\Delta,12}Z_{CL}\end{aligned}$$

$$\Rightarrow \qquad \tilde{\Delta}_{11} = -\frac{\mu_{\Delta,12}}{\sigma\mu_{\Delta,11}}\Delta_{12}+\frac{\mu_{\Delta,12}}{\mu_{\Delta,11}}Z_{CL}. \tag{13}$$

where $\tilde{X}$ denotes the value of a variable $X$ at the next generation, $C_1(T_1)=\Theta_{\text{deg}}$, $C_1(\tilde{T}_1)=\Theta_1+\mu_{\Delta,12}Z_{CL}$, and $C_1(\tilde{T}_0)=C_1(T_2)/\sigma$ because upon division the regulator is partitioned in proportion to daughter cell sizes.

Recall that no noise in division-plane positioning implies that the scaled size fluctuation upon division equals the scaled size fluctuation upon birth in the next generation (i.e. $\Delta S_3=S_3/\mu_{S,3}-1=S_3/\sigma/(\mu_{S,3}/\sigma)-1=\tilde{S}_1/\mu_{S,1}-1=\Delta\tilde{S}_1$), and expressing (9) as

$$\Delta_{ij}\approx L_{ij}\Delta S_j+M_{ij}(\Delta S_{j+1}-Z_j) \tag{14}$$

where

$$L_{ij}=-\Big(\Delta\alpha_{ij}+\frac{1-\Delta\lambda_{ij}}{\sigma_j^{1-\Delta\lambda_{ij}}-1}\Big) \quad\text{and}\quad M_{ij}=\frac{(1-\Delta\lambda_{ij})\sigma_j^{1-\Delta\lambda_{ij}}}{\sigma_j^{1-\Delta\lambda_{ij}}-1} \tag{15}$$

then from (12) and (13) we have:

two-phase master regulator of G2/M or division (phase 2 exit):

$$L_{22}\Delta S_2 + M_{22}(\Delta\tilde{S}_1 - Z_2) = -\frac{\mu_{\Delta,21}}{\mu_{\Delta,22}}(L_{21}\Delta S_1 + M_{21}(\Delta S_2 - Z_1) - Z_{CL}) \tag{16}$$

two-phase master regulator of G1/S (phase 1 exit):

$$L_{11}\Delta\tilde{S}_1 + M_{11}(\Delta\tilde{S}_2 - \tilde{Z}_1) = -\frac{\mu_{\Delta,12}}{\sigma\mu_{\Delta,11}}(L_{12}\Delta S_2 + M_{12}(\Delta\tilde{S}_1 - Z_2) - \sigma Z_{CL}). \tag{17}$$

From (16), for a two-phase master regulator of G2/M or division (phase 2 exit) we can derive the LSLS between birth size and birth size at the subsequent generation ($f_{S,11}$) and the LSLS between cell size at the beginning vs. the end of phase 2 ($f_{S,2}$) in terms of the LSLS between cell size at the beginning vs. the end of phase 1 ($f_{S,1}$), which is determined by the mechanism of control of phase 1 exit and other model parameters. Given $f_{S,11} = \mathbb{E}[\Delta\tilde{S}_1\Delta S_1]/\mathbb{E}[\Delta S_1^2]$, and $f_{S,1}/\sigma_1 = \mathbb{E}[\Delta S_2\Delta S_1]/\mathbb{E}[\Delta S_1^2]$, multiplying (16) by $\Delta S_1$, taking the expectation and dividing by $\mathbb{E}[\Delta S_1^2]$, we have

$$L_{22}\frac{f_{S,1}}{\sigma_1} + M_{22}f_{S,11} = -\frac{\mu_{\Delta,21}}{\mu_{\Delta,22}}(L_{21} + M_{21}\frac{f_{S,1}}{\sigma_1})$$

because again noise terms in the coordination of regulator production and growth ($Z_1, Z_2$) and the final level of regulator ($Z_{CL}$) are assumed to be zero-mean random variables that are independent of $\Delta S_1$. This equation gives

$$f_{S,11} = -\frac{\mu_{\Delta,21}}{\mu_{\Delta,22}}\frac{L_{21}}{M_{22}} - \frac{f_{S,1}}{\sigma_1}\Big(\frac{L_{22}}{M_{22}} + \frac{\mu_{\Delta,21}}{\mu_{\Delta,22}}\frac{M_{21}}{M_{22}}\Big). \tag{18}$$

Similarly, given $f_{S,2}/\sigma_2 = \mathbb{E}[\Delta\tilde{S}_1\Delta S_2]/\mathbb{E}[\Delta S_2^2]$, multiplying (16) by $\Delta S_2$, taking the expectation and dividing by $\mathbb{E}[\Delta S_2^2]$, and assuming steady state so $\mathrm{Var}(\Delta S_i) = \mathrm{Var}(\Delta\tilde{S}_i)$

$$\frac{f_{S,2}}{\sigma_2} = -\frac{L_{22}}{M_{22}} - \frac{\mu_{\Delta,21}}{\mu_{\Delta,22}}\frac{M_{21}}{M_{22}} - \frac{f_{S,1}}{\sigma_1}\frac{\mu_{\Delta,21}}{\mu_{\Delta,22}}\frac{L_{21}}{M_{22}}\frac{\mathrm{Var}(\Delta S_1)}{\mathrm{Var}(\Delta S_2)} \tag{19}$$

where $\mathrm{Var}(\Delta S_i)$ equals the square of the coefficient of variation of $S_i$.

For two-phase master regulator of G1/S (phase 1 exit) we can similarly derive the LSLS between cell size at the beginning of phase 2 vs. cell size at the beginning of phase 2 at the subsequent generation ($f_{S,22}$) and the LSLS between cell size at the beginning vs. the end of phase 1 ($f_{S,1}$) in terms of the LSLS between cell size at the beginning vs. the end of phase 2

($f_{S,2}$). Given $f_{S,22} = \mathbb{E}[\Delta\tilde{S}_2 \Delta S_2]/\mathbb{E}[\Delta S_2^2]$, and $f_{S,2}/\sigma_2 = \mathbb{E}[\Delta S_2 \Delta\tilde{S}_1]/\mathbb{E}[\Delta S_2^2]$, multiplying (17) by $\Delta S_2$, taking the expectation and dividing by $\mathbb{E}[\Delta S_2^2]$, we have:

$$f_{S,22} = -\frac{\mu_{\Delta,12}}{\sigma\mu_{\Delta,11}}\frac{L_{12}}{M_{11}} - \frac{f_{S,2}}{\sigma_2}\Big(\frac{L_{11}}{M_{11}} + \frac{\mu_{\Delta,12}}{\sigma\mu_{\Delta,11}}\frac{M_{12}}{M_{11}}\Big), \tag{20}$$

and similarly:

$$\frac{f_{S,1}}{\sigma_1} = -\frac{L_{11}}{M_{11}} - \frac{\mu_{\Delta,12}}{\sigma\mu_{\Delta,11}}\frac{M_{12}}{M_{11}} - \frac{f_{S,2}}{\sigma_2}\frac{\mu_{\Delta,12}}{\sigma\mu_{\Delta,11}}\frac{L_{12}}{M_{11}}\frac{\mathrm{Var}(\Delta S_2)}{\mathrm{Var}(\Delta S_1)}. \tag{21}$$

If G1 is an independently regulated phase, then:

$$\Delta\tilde{S}_2 \quad = \quad \frac{f_{S,1}}{\sigma_1}\Delta\tilde{S}_1 + \tilde{Z}_O,$$

and, to derive $f_{S,22}$ in terms of $f_{S,1}$, multiply this expression by $\Delta S_2$, take the expectation, and divide by $\mathbb{E}[\Delta S_2^2]$ to get:

$$f_{S,22} = \frac{f_{S,1}}{\sigma_1}\frac{f_{S,2}}{\sigma_2}$$

since $\tilde{Z}_O$ is a zero-mean random variable that is independent of $\Delta S_2$. If S/G2/M is an independently regulated phase, then similarly:

$$f_{S,11} = \frac{f_{S,1}}{\sigma_1}\frac{f_{S,2}}{\sigma_2}$$

implying:

$$\text{LSLS of birth vs. division size} = f_{S,1}f_{S,2}. \tag{22}$$

5.2.4. *Inhibitor-dilutors.* Recall that for an inhibitor-dilutor that triggers exit from phase $i$, we have

$$\Delta_{ij} \approx L_{ij}\Delta S_j + M_{ij}(\Delta S_{j+1} - Z_j) \text{ for } j \neq i, \quad \text{and} \quad C_i(T_i) - C_i(T_{i-1}) = \mu_{\Delta,i\,i-1}Z_{CL}$$

where as above

$$L_{ij} = -\Big(\Delta\alpha_{ij} + \frac{1-\Delta\lambda_{ij}}{\sigma_j^{1-\Delta\lambda_{ij}} - 1}\Big) \quad \text{and} \quad M_{ij} = \frac{(1-\Delta\lambda_{ij})\sigma_j^{1-\Delta\lambda_{ij}}}{\sigma_j^{1-\Delta\lambda_{ij}} - 1}.$$

If an inhibitor-dilutor triggers G2/M (phase 2 exit),

$$\begin{aligned}\Delta_{21} &= \frac{C_2(T_1) - C_2(T_0)}{\mu_{\Delta,21}} - 1 = \frac{C_2(T_2) - \mu_{\Delta,21} Z_{CL} - C_2(T_0)}{\mu_{\Delta,21}} - 1 \\ &= \frac{k_2(S_3 - S_1 - \mu_{S,3} Z_{ID} + \mu_{S,1} Z'_{ID})}{\mu_{\Delta,21}} - 1 - Z_{CL} \\ &= \frac{\sigma}{\sigma - 1}(\Delta S_3 - Z_{ID}) - \frac{1}{\sigma - 1}(\Delta S_1 - Z'_{ID}) - Z_{CL}\end{aligned}$$

where $Z'_{ID}$ is the noise in the inhibitor's threshold concentration in the previous cell cycle and given $\mu_{\Delta,21} = k_2 \mu_{S,1}(\sigma - 1)$ from equation (5). Since $\Delta S_3 = \Delta \tilde{S}_1$, for a phase 2 inhibitor-dilutor we have

$$\frac{\sigma}{\sigma - 1}(\Delta \tilde{S}_1 - Z_{ID}) - \frac{1}{\sigma - 1}(\Delta S_1 - Z'_{ID}) - Z_{CL} = L_{21} \Delta S_1 + M_{21}(\Delta S_2 - Z_1) \tag{23}$$

and similarly for a phase 1 inhibitor-dilutor, we have

$$\frac{\sigma}{\sigma - 1}(\Delta \tilde{S}_2 - \tilde{Z}_{ID}) - \frac{1}{\sigma - 1}(\Delta S_2 - Z_{ID}) - Z_{CL} = L_{12} \Delta S_2 + M_{12}(\Delta \tilde{S}_1 - Z_2). \tag{24}$$

From (23), for an inhibitor-dilutor regulating phase 2 exit we can derive the LSLS between birth size and birth size at the subsequent generation ($f_{S,11}$) and the LSLS between cell size at the beginning vs. the end of phase 2 ($f_{S,2}$) in terms of the LSLS between cell size at the beginning vs. the end of phase 1 ($f_{S,1}$). Given $f_{S,11} = \mathbb{E}[\Delta \tilde{S}_1 \Delta S_1]/\mathbb{E}[\Delta S_1^2]$, and $f_{S,1}/\sigma_1 = \mathbb{E}[\Delta S_2 \Delta S_1]/\mathbb{E}[\Delta S_1^2]$, multiplying (23) by $\Delta S_1$, taking the expectation and dividing by $\mathbb{E}[\Delta S_1^2] = \mathrm{Var}(\Delta S_1)$, we have

$$\frac{\sigma}{\sigma - 1} f_{S,11} - \frac{1}{\sigma - 1}\Big(1 - \frac{\mathbb{E}[Z'_{ID} \Delta S_1]}{\mathrm{Var}(\Delta S_1)}\Big) = L_{21} + M_{21} \frac{f_{S,1}}{\sigma_1}$$

which implies

$$f_{S,11} = \frac{\sigma - 1}{\sigma}\Big(L_{21} + M_{21} \frac{f_{S,1}}{\sigma_1}\Big) + \sigma^{-1}\Big(1 - \frac{\zeta_{ID}^2}{\mathrm{Var}(\Delta S_1)}\Big).$$

Except for the $Z'_{ID}$ noise term, again other noise terms disappear because they are zero-mean random variables that are independent of $\Delta S_1$. By contrast, by definition $\Delta S_1$ depends on the value of $Z'_{ID}$, that is, $\Delta S_1 = g(\Delta S'_2, \Delta S'_1, \ldots) + Z'_{ID}$ where $g(\cdot)$ is a function of previous transition sizes that are independent of $Z'_{ID}$, so we have $\mathbb{E}[\Delta S_1 Z'_{ID}] = \zeta_{ID}^2$.

Similarly, given $f_{S,2}/\sigma_2 = \mathbb{E}[\Delta\tilde{S}_1\Delta S_2]/\mathbb{E}[\Delta S_2^2]$, multiplying (23) by $\Delta S_2$, taking the expectation and dividing by $\mathbb{E}[\Delta S_2^2]$,

$$\frac{f_{S,2}}{\sigma_2} = \frac{1}{\sigma}\frac{f_{S,1}}{\sigma_1}\frac{\mathrm{Var}(\Delta S_1)}{\mathrm{Var}(\Delta S_2)}(1+(\sigma-1)L_{21}) + \frac{\sigma-1}{\sigma}M_{21} - \frac{1}{\sigma}\frac{\mathbb{E}[Z'_{ID}\Delta S_2]}{\mathrm{Var}(\Delta S_2)}.$$

If the phase 2 inhibitor dilutor is combined with an independently regulated phase 1, then

$$\Delta S_2 = \frac{f_{S,1}}{\sigma_1}\Delta S_1 + Z_1^O = \frac{f_{S,1}}{\sigma_1}(g(\Delta S'_2, \Delta S'_1, \ldots) + Z'_{ID}) + Z_1^O \quad \Rightarrow \quad \mathbb{E}[Z'_{ID}\Delta S_2] = \frac{f_{S,1}}{\sigma_1}\zeta_{ID}^2$$

and so

$$\frac{f_{S,2}}{\sigma_2} = \frac{1}{\sigma}\frac{f_{S,1}}{\sigma_1}\frac{\mathrm{Var}(\Delta S_1)}{\mathrm{Var}(\Delta S_2)}(1+(\sigma-1)L_{21}) + \frac{\sigma-1}{\sigma}M_{21} - \frac{1}{\sigma}\frac{f_{S,1}}{\sigma_1}\frac{\zeta_{ID}^2}{\mathrm{Var}(\Delta S_2)}.$$

Furthermore, because phase 1 is independently regulated, as derived above,

$$f_{S,22} = \frac{f_{S,1}}{\sigma_1}\frac{f_{S,2}}{\sigma_2}.$$

From (24), for an inhibitor-dilutor regulating G1/S (phase 1 exit) similar derivations give the LSLS between cell size at the beginning of phase 2 vs. cell size at the beginning of phase 2 in the subsequent generation ($f_{S,22}$) and the LSLS between cell size at the beginning vs. the end of phase 1 ($f_{S,2}$) in terms of the LSLS between cell size at the beginning vs. the end of phase 2 ($f_{S,2}$):

$$f_{S,22} = \frac{\sigma-1}{\sigma}\Big(L_{12} + M_{12}\frac{f_{S,2}}{\sigma_2}\Big) + \sigma^{-1}\Big(1 - \frac{\zeta_{ID}^2}{\mathrm{Var}(\Delta S_2)}\Big). \tag{25}$$

If G2/M (phase 2 exit) is triggered by an independently regulated mechanism, then:

$$\frac{f_{S,1}}{\sigma_1} = \frac{1}{\sigma}\frac{f_{S,2}}{\sigma_2}\frac{\mathrm{Var}(\Delta S_2)}{\mathrm{Var}(\Delta S_1)}(1+(\sigma-1)L_{12}) + \frac{\sigma-1}{\sigma}M_{12} - \frac{1}{\sigma}\frac{f_{S,2}}{\sigma_2}\frac{\zeta_{ID}^2}{\mathrm{Var}(\Delta S_1)} \tag{26}$$

and

$$f_{S,11} = \frac{f_{S,1}}{\sigma_1}\frac{f_{S,2}}{\sigma_2}.$$

5.3. **LSLSs between cell cycle transition sizes and phase durations.** To derive the LSLS $f_{t,j}$ relating cell size fluctuations at the beginning of phase $j$ ($\Delta S_j = S_j/\mu_{S,j} - 1$) to fluctuations in the duration of phase $j$ ($t_j/\mu_{t,j} - 1$ where $\mu_{t,j}$ is the average duration of phase $j$), we first solve equation (1):

$$(1+\alpha_{g,j}\Delta S_j)\,\gamma_j t_j = \frac{S_j^{1-\lambda_{g,j}} - S_j^{1-\lambda_{g,j}}}{1-\lambda_{g,j}}.$$

Taking means of either side

$$\gamma_j\,\mu_{t,j} \approx \frac{\mu_{S,j}^{1-\lambda_{g,j}}(\sigma_j^{1-\lambda_{g,j}}-1)}{1-\lambda_{g,j}} \tag{27}$$

then expanding in powers of cell size fluctuations and with some rearrangement, we have:

$$(1+\alpha_{g,j}\Delta S_j)\,\gamma_j t_j = \frac{\mu_{S,j}^{1-\lambda_{g,j}}}{1-\lambda_{g,j}}\big(\sigma_j^{1-\lambda_{g,j}} - 1 + (1-\lambda_{g,j})(\sigma_j^{1-\lambda_{g,j}}\Delta S_{j+1} - \Delta S_j) + \text{h.o.t.}\big).$$

Now dividing both sides by $\gamma_j\mu_{t,j}(1+\alpha_{g,j}\Delta S_j)$ and substituting (27):

$$t_j/\mu_{t,j} - 1 = (1-\lambda_{g,j})\frac{\sigma_j^{1-\lambda_{g,j}}}{\sigma_j^{1-\lambda_{g,j}}-1}\Delta S_{j+1} - \Big(\frac{1-\lambda_{g,j}}{\sigma_j^{1-\lambda_{g,j}}-1} + \alpha_{g,j}\Big)\Delta S_j + \text{h.o.t.} \tag{28}$$

So, assuming cell size fluctuations are small and terms other than first-order can be ignored, the LSLS $f_{t,j}$ between $t_j/\mu_{t,j} - 1$ and $\Delta S_j$, which is by definition $\mathbb{E}[(t_j/\mu_{t,j} - 1)\,\Delta S_j]/\mathbb{E}[\Delta S_j^2]$, is:

$$f_{t,j} \approx \frac{1-\lambda_{g,j}}{\sigma_j - \sigma_j^{\lambda_{g,j}}}\,(f_{S,j} - \sigma_j^{\lambda_{g,j}}) - \alpha_{g,j} \quad \to \quad \frac{f_{S,j}-\sigma_j}{\sigma_j \ln\sigma_j} - \alpha_{g,j} \text{ as } \lambda_{g,j}\to 1. \tag{29}$$

since $f_{S,j}/\sigma_j = \mathbb{E}[\Delta S_{j+1}\,\Delta S_j]/\mathbb{E}[\Delta S_j^2]$. For example, assuming $\alpha_{g,j} = 0$, *timer regulation of phase $j$ implies that $f_{t,j} = 0$, so we must have $f_{S,j} = \sigma_j^{\lambda_{g,j}}$*. If $f_{S_1} = 1/f_{S,2}$ and $f_{S,2} > 1$, a situation that can arise naturally (see special cases below), then $f_{t,1} < 0$ and the correlation between birth size and G1 duration is negative.

### 6. Special cases for two-phase master regulators

The expressions for relationships among size variables for two-phase master regulators are complicated but they simplify dramatically under specific assumptions. For example, if we assume: (1) the same dependence of growth and master regulator dynamics on initial cell size ($\Delta\alpha_{ij} = 0$); and (2) the master regulator has the same size dependence relative to growth during both phases ($\Delta\lambda_{ii} = \Delta\lambda_{ij} = \Delta\lambda_i$), then a two-phase master regulator that triggers G1/S (exit from phase 1), from definitions of $\mu_{\Delta,ij}$ (equation (8)), $L_{ij}$, and $M_{ij}$ (equations (15)), satisfies:

$$\frac{\mu_{\Delta,12}}{\sigma\mu_{\Delta,11}} = \frac{\gamma_1\kappa_{12}}{\gamma_2\kappa_{11}}\frac{1}{\sigma}\Big(\frac{\sigma_2^{1-\Delta\lambda_1}-1}{1-\sigma_1^{\Delta\lambda_1-1}}\Big), \quad \frac{L_{12}}{M_{11}} = \frac{1-\sigma_1^{\Delta\lambda_1-1}}{\sigma_2^{1-\Delta\lambda_1}-1}$$

$$\frac{L_{11}}{M_{11}} = -\sigma_1^{\Delta\lambda_1-1}, \quad \frac{M_{12}}{M_{11}} = \frac{1-\sigma_1^{\Delta\lambda_1-1}}{1-\sigma_2^{\Delta\lambda_1-1}}, \tag{30}$$

which upon substitution into (20) and (21) give:

$$f_{S,22} = \sigma^{-1}\frac{\gamma_1\kappa_{12}}{\gamma_2\kappa_{11}} - \sigma_1^{-1}\Big(\frac{\gamma_1\kappa_{12}}{\gamma_2\kappa_{11}}\sigma_2^{-\Delta\lambda_1} - \sigma_1^{\Delta\lambda_1}\Big)\frac{f_{S,2}}{\sigma_2} \tag{31}$$

$$\frac{f_{S,1}}{\sigma_1} = \Big(1 - \frac{\gamma_1\kappa_{12}}{\gamma_2\kappa_{11}}\sigma^{-\Delta\lambda_1}\Big)\sigma_1^{\Delta\lambda_1-1} + \frac{f_{S,2}}{\sigma_2}\sigma^{-1}\frac{\gamma_1\kappa_{12}}{\gamma_2\kappa_{11}}\frac{\mathrm{Var}(\Delta S_2)}{\mathrm{Var}(\Delta S_1)}. \tag{32}$$

From the equation for $f_{S,22}$, assuming $\gamma_1 = \gamma_2$, we see that, regardless of noise and further restrictions on $\Delta\lambda_i$ or $\sigma_1$, $f_{22} \geq 1$ for phase 2 sizer regulation so $f_{S,2} = 0$, and $\kappa_{12}/\kappa_{11} = 2$ (i.e. gene copy-number limited production), and $\sigma \leq 2$. For $f_{S,2} > 0$ and $\Delta\lambda_i \ll -1$, the term $\sigma_2^{-\Delta\lambda_1}$ becomes large and thus $f_{S,22} < -1$, regardless of other parameters. If we further assume either that the G1/S master regulator is produced in proportion to growth rate throughout the cell-cycle ($\Delta\lambda_i = 0, \kappa_{ij} = \text{constant } \times\gamma_j$), or that growth is exponential ($\gamma_1 = \gamma_2$ and $\lambda_{g,1} = \lambda_{g,2} = 1$) while the regulator is produced proportionally to cell size ($\Delta\lambda_i = 0$, but possibly $\kappa_{12} \neq \kappa_{11}$), then from (31) and (32) (case for $\kappa_{12} \neq \kappa_{11}$):

$$\begin{aligned} f_{S,22} &= \sigma^{-1} & &\Big(= \sigma^{-1}\frac{\kappa_{12}}{\kappa_{11}} - \sigma_1^{-1}(\frac{\kappa_{12}}{\kappa_{11}} - 1)\frac{f_{S,2}}{\sigma_2}\Big) \\ f_{S,1}/\sigma_1 &= \sigma^{-1}\frac{f_{S,2}}{\sigma_2}\frac{\mathrm{Var}(\Delta S_2)}{\mathrm{Var}(\Delta S_1)} & &\Big(= \sigma_1^{-1}(1 - \frac{\kappa_{12}}{\kappa_{11}}) + \sigma^{-1}\frac{\kappa_{12}}{\kappa_{11}}\frac{f_{S,2}}{\sigma_2}\frac{\mathrm{Var}(\Delta S_2)}{\mathrm{Var}(\Delta S_1)}\Big) \end{aligned}$$

and, if the second phase (S/G2/M) is independently regulated, from equation (22):

$$\text{LSLS of birth size vs. division size} \quad = \quad f_{S,1}f_{S,2}.$$

Substituting the expression for $f_{S,1}$ in the case when regulator production is proportional to growth ($\kappa_{12} = \kappa_{11}$), we get a condition for apparent 'adder' regulation:

$$\text{LSLS of birth size vs. division size } = (f_{S,2}/\sigma_2)^2\,\mathrm{Var}(\Delta S_2)/\mathrm{Var}(\Delta S_1) \approx 1.$$

The independent regulation of phase 2 implies

$$\Delta\tilde{S}_1 = \frac{f_{S,2}}{\sigma_2}\Delta S_2 + Z_O$$

so, at steady state, taking variances of both sides,

$$\mathrm{Var}(\Delta S_1) = (f_{S,2}/\sigma_2)^2\mathrm{Var}(\Delta S_2) + \zeta_O^2$$

where $\mathrm{Var}(\Delta S_2)$ is equivalent to the square of the coefficient of variation in cell size at G1/S. Therefore, substituting this expression for $\mathrm{Var}(\Delta S_1)$,

$$\text{LSLS of birth size vs. division size } = \frac{(f_{S,2}/\sigma_2)^2}{(f_{S,2}/\sigma_2)^2 + \eta_{\mathrm{G2/M}}^2}$$

where $\eta_{\mathrm{G2/M}} = \zeta_O/$coefficient of variation of G1/S size. Consequently we see that, if $\eta_{\mathrm{G2/M}} \ll f_{S,2}/\sigma_2$, then apparent adder regulation is observed between birth and division regardless of the mechanism of S/G2/M control (adder, timer, etc.), and further then $f_{S,1} \approx 1/f_{S,2}$. For larger values of $\eta_{\mathrm{G2/M}}$, this scenario produces sub-adder regulation between birth and division. Between consecutive G1/Ss, true adder regulation is executed, where a constant amount of material is added between DNA replication initiations ignoring the intervening division regardless of noise levels ($\sigma f_{S,22} = 1$).

If rather than controlling G1/S the two-phase master regulator triggers G2/M or division (phase 2 exit), the results are analogous. As above, assuming $\Delta\alpha_{ij} = 0$ and $\Delta\lambda_{ii} = \Delta\lambda_{ij} = \Delta\lambda_i$, from definitions of $\mu_{\Delta,ij}$ (equation (8)), $L_{ij}$, and $M_{ij}$ (equations (15)), we have:

$$\begin{aligned}
&\frac{\mu_{\Delta,21}}{\mu_{\Delta,22}} = \frac{\gamma_2\kappa_{21}}{\gamma_1\kappa_{22}}\Big(\frac{1-\sigma_1^{\Delta\lambda_2-1}}{\sigma_2^{1-\Delta\lambda_2}-1}\Big), \quad \frac{L_{21}}{M_{22}} = \frac{1-\sigma_2^{\Delta\lambda_2-1}}{1-\sigma_1^{1-\Delta\lambda_2}} \\
&\frac{L_{22}}{M_{22}} = -\sigma_2^{\Delta\lambda_2-1}, \quad \frac{M_{21}}{M_{22}} = \frac{1-\sigma_2^{\Delta\lambda_2-1}}{1-\sigma_1^{\Delta\lambda_2-1}},
\end{aligned} \tag{33}$$

which upon substitution into (18) and (19) give:

$$\begin{aligned}
f_{S,11} &= \sigma^{\Delta\lambda_2-1}\frac{\gamma_2\kappa_{21}}{\gamma_1\kappa_{22}} - \sigma_2^{\Delta\lambda_2-1}\Big(\frac{\gamma_2\kappa_{21}}{\gamma_1\kappa_{22}}-1\Big)\frac{f_{S,1}}{\sigma_1} \\
\frac{f_{S,2}}{\sigma_2} &= \Big(1-\frac{\gamma_2\kappa_{21}}{\gamma_1\kappa_{22}}\Big)\sigma_2^{\Delta\lambda_2-1} + \frac{f_{S,1}}{\sigma_1}\sigma^{\Delta\lambda_2-1}\frac{\gamma_2\kappa_{21}}{\gamma_1\kappa_{22}}\frac{\mathrm{Var}(\Delta S_1)}{\mathrm{Var}(\Delta S_2)}.
\end{aligned}$$

Now for regulator production proportional to growth ($\Delta\lambda_i = 0, \kappa_{ij} =$ constant $\times \gamma_j$), or for exponential growth ($\gamma_1 = \gamma_2$ and $\lambda_{g,1} = \lambda_{g,2} = 1$) while the regulator is produced in proportion to cell size ($\Delta\lambda_i = 0$, but possibly $\kappa_{12} \neq \kappa_{11}$), then (case for $\kappa_{12} \neq \kappa_{11}$):

$$
\begin{aligned}
f_{S,11} &= \sigma^{-1} \quad \Big(= \sigma^{-1}\frac{\kappa_{21}}{\kappa_{22}} - \sigma_2^{-1}\big(\frac{\kappa_{21}}{\kappa_{22}} - 1\big)\frac{f_{S,1}}{\sigma_1}\Big) \\
f_{S,2}/\sigma_2 &= \frac{f_{S,1}}{\sigma_1}\sigma^{-1}\frac{\mathrm{Var}(\Delta S_1)}{\mathrm{Var}(\Delta S_2)} \qquad \Big(= \sigma_2^{-1}(1 - \frac{\kappa_{21}}{\kappa_{22}}) + \frac{f_{S,1}}{\sigma_1}\sigma^{-1}\frac{\kappa_{21}}{\kappa_{22}}\frac{\mathrm{Var}(\Delta S_1)}{\mathrm{Var}(\Delta S_2)}\Big).
\end{aligned}
$$

Furthermore, assuming the first phase is regulated independently (e.g. sizer, adder, or timer) and steady state, then:

$$
\begin{aligned}
f_{S,22} &= \frac{f_{S,2}}{\sigma_2}\frac{f_{S,1}}{\sigma_1} \\
\mathrm{Var}(\Delta S_2) &= (f_{S,1}/\sigma_1)^2\mathrm{Var}(\Delta S_1) + \zeta_O^2.
\end{aligned}
$$

The conclusions are analogous to the case for a G1/S two-phase master regulator. For exponentially growing cells with $\gamma_1 = \gamma_2$ and size-independent regulator production ($\Delta\lambda_i = 0$),

$$
\begin{aligned}
f_{S,11} &= \frac{\kappa_{21}}{\kappa_{22}} - \Big(\frac{\kappa_{21}}{\kappa_{22}} - 1\Big)\frac{f_{S,1}}{\sigma_1} \\
\frac{f_{S,2}}{\sigma_2} &= \Big(1 - \frac{\kappa_{21}}{\kappa_{22}}\Big) + \frac{f_{S,1}}{\sigma_1}\frac{\kappa_{21}}{\kappa_{22}}\frac{\mathrm{Var}(\Delta S_1)}{\mathrm{Var}(\Delta S_2)}.
\end{aligned}
$$

For $\kappa_{21}/\kappa_{22} = 1/2$ (i.e. gene copy-number limited production), $f_{S,11} = 1$ for timer control of phase 1 when $f_{S,1} = \sigma_1$, but $f_{S,11} < 1$ when $f_{S,1} < \sigma_1$, e.g. for adder or sizer regulation of phase 1.

### 7. Special cases for inhibitor-dilutors

In the following we assume: (1) an inhibitor-dilutor triggers G1/S (phase 1 exit); (2) growth is exponential ($\lambda_{g,1} = \lambda_{g,2} = 1$); (3) the inhibitor-dilutor is produced at a constant size-independent rate during S/G2/M ($\lambda_{c,12} = 0$); (4) there is no difference between initial cell size dependence of exponential growth and inhibitor production during S/G2/M ($\Delta\alpha_{12} = 0$); (5) G2/M (phase 2 exit) is triggered by an independently regulated mechanism, e.g. sizer, adder, or timer. Then from equations (15):

$$M_{12} = (\ln \sigma_2)^{-1}, L_{12} = -(\ln \sigma_2)^{-1}$$

and, substituting these values into (25) and (26), the LSLS between cell sizes at consecutive G1/Ss is:

$$f_{S,22} = (\ln \sigma_2)^{-1}\frac{\sigma - 1}{\sigma}\Big(\frac{f_{S,2}}{\sigma_2} - 1\Big) + \sigma^{-1}\Big(1 - \frac{\zeta_{ID}^2}{\mathrm{Var}(\Delta S_2)}\Big)$$

while the LSLS between birth size vs. G1/S size is:

$$\frac{f_{S,1}}{\sigma_1} = \frac{\sigma - 1}{\sigma}(\ln \sigma_2)^{-1}\Big(1 - \frac{f_{S,2}}{\sigma_2}\frac{\mathrm{Var}(\Delta S_2)}{\mathrm{Var}(\Delta S_1)}\Big) + \frac{1}{\sigma}\frac{f_{S,2}}{\sigma_2}\Big(\frac{\mathrm{Var}(\Delta S_2) - \zeta_{ID}^2}{\mathrm{Var}(\Delta S_1)}\Big)$$

and the LSLS between birth and division sizes is:

$$\sigma f_{S,11} = f_{S,1}\, f_{S,2}.$$

S/G2/M is assumed to be governed by an independently regulated mechanism so $\mathrm{Var}(\Delta S_1) = \mathrm{Var}(\Delta S_2) + \zeta_O^2$. If the mechanism is timer, then from (29), assuming no dependence of growth on initial cell size ($\alpha_{g,j} = 0$), $f_{S,2} = \sigma_2$. Now substituting these relations:

$$f_{S,22} = \sigma^{-1}\Big(1 - \frac{\zeta_{ID}^2}{\mathrm{Var}(\Delta S_2)}\Big) = \sigma^{-1}(1 - \eta_{\mathrm{G1/S}}^2) \tag{34}$$

$$\begin{aligned}\text{LSLS of birth vs. division size} &= \sigma f_{S,11} \\ &= \frac{\sigma - 1}{\ln \sigma_2}\Big(1 - \frac{\mathrm{Var}(\Delta S_2)}{\mathrm{Var}(\Delta S_2) + \zeta_O^2}\Big) + \frac{\mathrm{Var}(\Delta S_2) - \zeta_{ID}^2}{\mathrm{Var}(\Delta S_2) + \zeta_O^2} \\ &= \frac{1 + \frac{\sigma - 1}{\ln \sigma_2}\eta_{\mathrm{G2/M}}^2 - \eta_{\mathrm{G1/S}}^2}{1 + \eta_{\mathrm{G2/M}}^2}. \qquad (35)\end{aligned}$$

where $\eta_{\mathrm{G1/S}} = \zeta_{ID}/$coefficient of variation of cell size at G1/S and, as above, $\eta_{\mathrm{G2/M}} = \zeta_O/$coefficient of variation of cell size at G1/S. If instead S/G2/M is regulated by an adder mechanism, then $f_{S,2} = 1$, and

$$\begin{aligned}\text{LSLS of birth vs. division size} &= \sigma f_{S,11} \\ &= \frac{1 - \frac{(\sigma - 1)(\sigma_2 - 1)}{\sigma_2 \ln \sigma_2} + \frac{\sigma - 1}{\sigma_2 \ln \sigma_2}\sigma_2^2 \eta_{\mathrm{G2/M}}^2 - \eta_{\mathrm{G1/S}}^2}{1 + \sigma_2^2 \eta_{\mathrm{G2/M}}^2}.\end{aligned}$$

If S/G2/M is regulated by a sizer mechanism, then $f_{S,2} = 0$, and

$$\begin{aligned}f_{S,22} &= \sigma^{-1} - \frac{1}{\ln \sigma_2} + \frac{\sigma^{-1}}{\ln \sigma_2} - \sigma^{-1}\eta_{\mathrm{G1/S}}^2 \\ \text{LSLS of birth vs. division size} &= 0.\end{aligned}$$

Now we consider different cases for S/G2/M independently regulated control if the inhibitor is produced with the same size-dependence as growth through S/G2/M ($\Delta\lambda_{12} = 0$). Then

$$M_{12} = \frac{\sigma_2}{\sigma_2 - 1}, L_{12} = -\frac{1}{\sigma_2 - 1}.$$

For S/G2/M timer control and assuming exponential growth, $f_{S,2} = \sigma_2$, and

$$\begin{aligned} f_{S,22} &= 1 - \sigma^{-1}\eta^2_{\mathrm{G1/S}} \\ f_{S,11} &= \frac{1 + \frac{1-\sigma^{-1}}{1-\sigma_2^{-1}}\eta^2_{\mathrm{G2/M}} - \sigma^{-1}\eta^2_{\mathrm{G1/S}}}{1 + \eta^2_{\mathrm{G2/M}}}. \\ \frac{f_{S,1}}{\sigma_1} &= f_{S,11}/(f_{S,2}/\sigma_2). \end{aligned}$$

In the absence of noise ($\eta_{\mathrm{G1/S}} = \eta_{\mathrm{G2/M}} = 0$), $f_{S,22} = f_{S,11} = 1$, and $f_{S,1} = \sigma_1$ which implies G1 is also effectively regulated by timer control (since from equation (29) there is no dependence on G1 duration on birth size). Size homeostasis is lost.

For S/G2/M sizer control with no assumptions on growth,

$$\begin{aligned} f_{S,22} &= -\sigma^{-1}\frac{\sigma - \sigma_2}{\sigma_2 - 1} - \sigma^{-1}\eta^2_{\mathrm{G1/S}} \\ f_{S,11} &= 0. \end{aligned}$$

Assuming linear growth and S/G2/M timer/adder control, then $f_{S,2} = 1$, and

$$\begin{aligned} f_{S,22} &= \sigma^{-1}(1 - \eta^2_{\mathrm{G1/S}}) \\ f_{S,11} &= \sigma^{-1}\frac{1 + \left(\frac{\sigma-1}{\sigma_2-1}\right)\sigma_2^2\eta^2_{\mathrm{G2/M}} - \eta^2_{\mathrm{G1/S}}}{1 + \sigma_2\eta^2_{\mathrm{G2/M}}}. \end{aligned}$$

### 8. Checkpoint progression triggered at a threshold local density in a cellular region that scales with size $\sim S^{\lambda_T}$

We first suppose that a master regulator triggers phase progression not at a threshold level but instead at a threshold local density, denoted by $k_{\mathrm{crit}}$, within a cellular compartment that scales with cell size according to $S^{\lambda_T}$, whereupon the regulator is degraded to a fixed level $\Theta_{\mathrm{deg}}$. We derive the corresponding LSLS between cell size at the beginning vs. cell size at the end of the phase. The analysis is also applicable to two-phase master regulators that have identical parameters through both phases ($\lambda_{i1} = \lambda_{i2}$, $\kappa_{i1} = \kappa_{i2}$) when growth parameters are also constant ($\gamma_1 = \gamma_2, \lambda_{g1} = \lambda_{g2}$).

The analysis differs beginning from equation (9). With this new criterion for phase progression, $\Delta_{ii} = (C(T_i) - C(T_{i-1}))/\mu_{\Delta,ii} - 1 = \lambda_T \Delta S_{i+1}/(1 - \Theta_{\text{deg}}/k_{\text{crit}}\mu_{S,i+1}^{\lambda_T}) + \mu_{\Delta,ii-1}/\mu_{\Delta,ii} Z_{CL}$, and from (9),

$$\Delta S_{i+1} = \frac{\sigma_i^{\Delta\lambda_{ii}-1} + \Delta\alpha_{ii}\frac{1-\sigma_i^{\Delta\lambda_{ii}-1}}{1-\Delta\lambda_{ii}}}{1 - \frac{1-\sigma_i^{\Delta\lambda_{ii}-1}}{1-\Delta\lambda_{ii}}\frac{\lambda_T}{1-\Theta_{\text{deg}}/k_{\text{crit}}\mu_{S,i+1}^{\lambda_T}}}\Delta S_i + \text{h.o.t.} + \beta_S Z_i + \beta_{CL} Z_{CL}$$

where $\beta_S$ and $\beta_{CL}$ are constants that have no affect on results. Following previous sections in deriving the LSLS between $\Delta S_i$ and $\Delta S_{i+1}$, we have:

$$\text{LSLS of } \Delta S_i \text{ vs. } \Delta S_{i+1} \quad \approx \quad \frac{\sigma_i^{\Delta\lambda_{ii}-1} + \Delta\alpha_{ii}\frac{1-\sigma_i^{\Delta\lambda_{ii}-1}}{1-\Delta\lambda_{ii}}}{1 - \frac{1-\sigma_i^{\Delta\lambda_{ii}-1}}{1-\Delta\lambda_{ii}}\frac{\lambda_T}{1-\Theta_{\text{deg}}/k_{\text{crit}}\mu_{S,i+1}^{\lambda_T}}}.$$

Assuming that the master regulator localizes to a compartment that scales directly with cell size ($\lambda_T = 1$), is degraded to zero ($\Theta_{\text{deg}} = 0$), and that $\Delta\alpha_{ii} = 0$,

$$\text{LSLS of } \Delta S_i \text{ vs. } \Delta S_{i+1} \quad \approx \frac{\sigma_i^{\Delta\lambda_{ii}-1}}{1 - \frac{1-\sigma_i^{\Delta\lambda_{ii}-1}}{1-\Delta\lambda_{ii}}}.$$

which produces a LSLS of 1 if $\Delta\lambda_{ii} = 0$. Thus, a two-phase master regulator which triggers G2/M at a threshold local density in any cellular region that scales proportionally with cell size, and that has the same size-dependencies as growth through G1 and S/G2/M, has a LSLS slope of 1 between cell-size fluctuations in consecutive phases and thus fails to achieve size-homeostasis.

We now consider G1/S (phase 1 exit) inhibitor dilutors that trigger phase progression at a threshold minimum local density, where the inhibitor localizes to a cellular compartment that scales with cell size according to $S^{\lambda_T}$ and it is the local concentration within the compartment that triggers phase progression. The analysis differs beginning from equation (24), where instead of equation (24) we have

$$\frac{\lambda_T \sigma^{\lambda_T}}{\sigma^{\lambda_T} - 1}(\Delta\tilde{S}_2 - \tilde{Z}_1) - \frac{\lambda_T}{\sigma^{\lambda_T} - 1}(\Delta S_2 - Z_1) - Z_{CL} = L_{12}\Delta S_2 + M_{12}(\Delta\tilde{S}_1 - Z_2).$$

Preceding analogously to the derivation following equation (24), we have

$$f_{S,22} = \frac{\sigma^{\lambda_T} - 1}{\lambda_T \sigma^{\lambda_T}}\Big(L_{12} + M_{12}\frac{f_2}{\sigma_2}\Big) + \sigma^{-\lambda_T}\Big(1 - \frac{\zeta_{ID}^2}{\mathrm{Var}(\Delta S_2)}\Big),$$

and

$$\frac{f_1}{\sigma_1} = \frac{1}{\lambda_T \sigma^{\lambda_T}}\frac{f_2}{\sigma_2}\frac{\mathrm{Var}(\Delta S_2)}{\mathrm{Var}(\Delta S_1)}(\lambda_T + (\sigma^{\lambda_T} - 1)L_{12}) + \frac{\sigma^{\lambda_T} - 1}{\lambda_T \sigma^{\lambda_T}} M_{12} - \sigma^{-\lambda_T}\frac{f_2}{\sigma_2}\frac{\zeta_{ID}^2}{\mathrm{Var}(\Delta S_1)}.$$

For a budding yeast-like inhibitor dilutor, where growth is exponential ($\lambda_{g,1} = \lambda_{g,2} = 1$), the inhibitor-dilutor is produced at a constant size-independent rate during S/G2/M ($\lambda_{c,12} = 0$), there is no difference between initial cell size dependence of exponential growth and inhibitor production during S/G2/M ($\Delta\alpha_{12} = 0$), and SG2/M (phase 2 exit) is a timer mechanism, then as above,

$$M_{12} = (\ln \sigma_2)^{-1}, L_{12} = -(\ln \sigma_2)^{-1}, f_2 = \sigma_2, \mathrm{Var}(\Delta S_1) = \Big(\frac{f_2}{\sigma_2}\Big)^2 \mathrm{Var}(\Delta S_2) + \zeta_O^2$$

and

$$f_{S,22} = \sigma^{-\lambda_T}(1 - \eta^2_{\mathrm{G1/S}})$$

implying a slope between G1/S size and added size until the next G1/S of $\sigma f_{S,22} - 1 = \sigma^{1-\lambda_T}(1 - \eta^2_{\mathrm{G1/S}}) - 1$. Also

$$f_{S,11} = \frac{f_1}{\sigma_1}\frac{f_2}{\sigma_2} = \sigma^{-\lambda_T}(1 + \Big(\frac{\sigma^{\lambda_T} - 1}{\lambda_T \ln \sigma_2}\Big)\eta^2_{\mathrm{G2/M}} - \eta^2_{\mathrm{G1/S}})$$

implying a slope between birth size and division size of $\sigma f_{11} = \sigma^{1-\lambda_T}(1 + \Big(\frac{\sigma^{\lambda_T}-1}{\lambda_T \ln \sigma_2}\Big)\eta^2_{\mathrm{G2/M}} - \eta^2_{\mathrm{G1/S}})$.

### 9. Special cases for two-phase master regulators with the same parameters through both phases

If a G2/M two-phase master regulator is produced throughout the cell cycle with both regulator production and growth maintaining their initial per unit size rates ($\lambda_{c,21} = \lambda_{c,22}, \lambda_{g,1} = \lambda_{g,2}, \kappa_{21} = \kappa_{22}, \gamma_1 = \gamma_2$ and any initial dependence on birth size is maintained throughout the cell cycle), then the analytical derivations for one-phase master regulators are clearly applicable. When further regulator production is proportional to growth ($\Delta\lambda_{ij} = 0$), then from equation (11), we have:

$$\text{LSLS of birth vs. division size} = 1 + \Delta\alpha(\sigma - 1)$$

where $\Delta\alpha = \alpha_g - \alpha_c$ represents the difference in dependence of regulator production vs. growth on cell birth size that persists throughout the cell cycle, and $\sigma$ is the overall fold-size change throughout the cell cycle.

### 10. The size-homeostasis behavior between birth and division and consecutive G1/Ss is identical when one phase is independently regulated with low noise

We show that if one phase is independently regulated with relatively low noise ($\zeta_O$/coefficient of variation of G1/S size $\ll 1$), then the size-homeostasis behavior between birth and division and consecutive G1/Ss are the same. We consider the case of an independently regulated S/G2/M, derivations for G1 being analogous. Recall that for an independently regulated S/G2/M, we have:

$$\mathrm{Var}(\Delta S_1) = \Big(\frac{f_{S,2}}{\sigma_2}\Big)^2 \mathrm{Var}(\Delta S_2) + \zeta_O^2 \implies \mathrm{Var}(\Delta S_1)/\mathrm{Var}(\Delta S_2) = \Big(\frac{f_{S,2}}{\sigma_2}\Big)^2 + \eta^2_{\mathrm{G2/M}},$$

and:

$$\Delta S_1' = \frac{f_{S,2}}{\sigma_2}\Delta S_2' + Z_O', \qquad \Delta S_1 = \frac{f_{S,2}}{\sigma_2}\Delta S_2 + Z_O,$$

where $'$ denotes the previous phase and $Z_O'$ and $Z_O$ are independent random variables. Therefore

$$\Delta S_1'\Delta S_1 = \Big(\frac{f_{S,2}}{\sigma_2}\Big)^2 \Delta S_2'\Delta S_2 + \frac{f_{S,2}}{\sigma_2}\Delta S_2 Z_O' + \frac{f_{S,2}}{\sigma_2}\Delta S_2' Z_O + Z_O Z_O'$$

and taking the expectation and dividing by $\mathbb{E}[\Delta S_1^2] = \mathrm{Var}(\Delta S_1)$, at steady state we have:

$$f_{11} = \Big(\frac{f_{S,2}}{\sigma_2}\Big)^2 f_{22}\frac{\mathrm{Var}(\Delta S_2)}{\mathrm{Var}(\Delta S_1)} + \frac{f_{S,2}}{\sigma_2}\frac{\mathbb{E}[\Delta S_2 Z_O']}{\mathrm{Var}(\Delta S_1)} = \frac{\Big(\frac{f_{S,2}}{\sigma_2}\Big)^2}{\Big(\frac{f_{S,2}}{\sigma_2}\Big)^2 + \eta^2_{\mathrm{G2/M}}} f_{22} + \frac{f_{S,2}}{\sigma_2}\frac{\mathbb{E}[\Delta S_2 Z_O']}{\mathrm{Var}(\Delta S_1)}$$

because $\Delta S_2'$ and $Z_O'$ are independent of $Z_O$. In the general case, $\Delta S_2 = \alpha_1\Delta S_1' + \alpha_2\Delta S_2' +$ size fluctuations from prior generations $+Z = \alpha_1(\frac{f_{S,2}}{\sigma_2}\Delta S_2' + Z_O') + \alpha_2\Delta S_2' +$ size fluctuations from prior generations $+Z$, where $Z$ is a random variable and $\alpha_i$ depend on the control type of the other phase. So, because all size fluctuations preceding $\Delta S_2'$ are independent of $Z_O'$,

$$\mathbb{E}[\Delta S_2 Z_O'] = \alpha_1\zeta_O^2$$

and hence

$$f_{11} = \left(\frac{f_{S,2}}{\sigma_2}\right)^2 f_{22} \frac{\mathrm{Var}(\Delta S_2)}{\mathrm{Var}(\Delta S_1)} + \frac{f_{S,2}}{\sigma_2} \frac{\mathbb{E}[\Delta S_2 Z'_O]}{\mathrm{Var}(\Delta S_1)} = \frac{\left(\frac{f_{S,2}}{\sigma_2}\right)^2}{\left(\frac{f_{S,2}}{\sigma_2}\right)^2 + \eta^2_{\mathrm{G2/M}}} f_{22} + \frac{f_{S,2}}{\sigma_2}\alpha_1 \eta^2_{\mathrm{G2/M}}.$$

So $f_{S,11} \approx f_{S,22}$ for supra-sizer regulation of G2/M and low G2/M checkpoint noise compared with other noise sources ($\eta_{\mathrm{G2/M}} \ll f_{S,2}/\sigma_2$), meaning that the apparent size-homeostasis behavior between consecutive G2/Ms (or birth and division) and consecutive G1/Ss is identical so long as cell size fluctuations are small and thus the linearization is appropriate, regardless of all other assumptions.

### 11. The size-homeostasis behavior between the added size over G1 vs. the added size over S/G2/M is a function of other cell size-homeostasis statistics

Recall that by definition the linear regression slope of $X$ vs. $Y$ is $\mathbb{E}[(X-\mu_X)(Y-\mu_Y)]/\mathbb{E}[(X-\mu_X)^2]$. Using the notation above, the fluctuations from the mean added-size scaled by the mean birth size over G1 and S/G2/M are $\sigma_1 \Delta S_2 - \Delta S_1$ and $\sigma \Delta S_3 - \sigma_1 \Delta S_2$, respectively. So the linear regression slope between added size over G1 vs. added size over S/G2/M is

$$\begin{aligned}\frac{\mathbb{E}[(\sigma\Delta S_3 - \sigma_1 \Delta S_2)(\sigma_1 \Delta S_2 - \Delta S_1)]}{\mathbb{E}[(\sigma_1 \Delta S_2 - \Delta S_1)^2]} &= \frac{\mathbb{E}[(\sigma\sigma_1 \Delta S_3 \Delta S_2 - \sigma_1^2 \Delta S_2^2 - \sigma \Delta S_3 \Delta S_1 - \sigma_1 \Delta S_2 \Delta S_1)]}{\mathbb{E}[\sigma_1^2 \Delta S_2^2 - 2\sigma_1 \Delta S_2 \Delta S_1 + \Delta S_1^2]} \\ &= \frac{\sigma_1^2 f_{S,2}\left(\frac{\rho_{\mathrm{G1/S}}}{\rho_{\mathrm{G2/M}}}\right)^2 - \sigma_1^2\left(\frac{\rho_{\mathrm{G1/S}}}{\rho_{\mathrm{G2/M}}}\right)^2 - f_{bd} + f_{S,1}}{\sigma_1^2\left(\frac{\rho_{\mathrm{G1/S}}}{\rho_{\mathrm{G2/M}}}\right)^2 - 2f_{S,1} + 1}.\end{aligned}$$

where $\rho_{\mathrm{G1/S}}$ and $\rho_{\mathrm{G2/M}}$ are the coefficients of variation in cell size at G1/S and G2/M, respectively, $f_{bd}$ is the linear regression slope between birth and G2/M, and $f_{S,1}$ and $f_{S,2}$ are the linear regression slopes between birth size and G1/S size, and G1/S size and G2/M size, respectively. This derivation used only algebraic manipulations and no modeling assumptions, thus it holds regardless of all modeling assumptions.
